## Supplement for "Sequence-structure-function relationships in the microbial protein universe"

#### Supplementary information

#### Table of Contents

1. Data availability on Zenodo and Github ... 3
2. MIP dataset construction ... 3
3. Protein structure predictions ... 3
   3.1. Sequence alignments ... 3
   3.2. Fragment picking ... 4
   3.3. Residue-residue contact predictions ... 4
   3.4. Rosetta model building on the IBM World Community Grid ... 6
   3.5. DMPfold model building ... 7
4. Function prediction ... 8
5. MIP dataset descriptions ... 9
   5.1. MIP dataset curation ... 10
   5.2. Sequence length distribution ... 11
   5.3. Pairwise structural comparisons in the curated dataset via CATH ... 12
6. Model quality assessment on the MIP_random5000 dataset ... 13
   6.1. Explanation of variables ... 13
   6.2. DMPFold models have higher coil content than Rosetta models ... 16
   6.3. Model quality is independent of sequence length and correlates with agreement between Rosetta and DMPFold models ... 17
   6.4. Deeper Multiple Sequence Alignment leads to higher model quality ... 18
   6.5. Transmembrane spans ... 19
   6.6. Disorder ... 20
   6.7. Model complexity ... 21
   6.8. TM-score to CATH ... 22
   6.9. Rosetta score ... 23
7. Additional results on the MIP_curated dataset ... 25
   7.1. Transmembrane spans ... 25
   7.2. Contact order ... 25

#### 1. Data availability on Zenodo and Github

All datasets are publicly accessible on Zenodo (url https://doi.org/10.5281/zenodo.6611431). This includes workflows and scripts to search the database using a query sequence, a query structure, or a query function, to find similar proteins in the MIP dataset.

Further, scripts for various data analyses are located on Github at https://github.com/microbiome-immunity-project/protein_universe.

#### 2. MIP dataset construction

The MIP dataset is constructed on the basis of GEBA1003 representative bacterial and archeal genomes from across the tree of life[1]. The dataset includes environmental samples from soil, ocean water, human gut microbiome and was designed to sample the microbial tree of life evenly. For each genome, we generated a list of predicted genes using Prodigal[2].

The raw gene catalog was processed using an incremental clustering approach, similar to the one employed by UniClust[3]. First, redundancy in the dataset was removed by using `linclust` (ie. clustering at 100% sequence identity), as implemented in MMSeqs2[4,5]. Then, the dataset was further clustered into 90%, 70% and 30% sequence identity clusters, with the last step (70% to 30% clustering) executed using the MMSeqs2 `clust` module.

The resulting dataset was sorted according to sequence length, sampling the entirety of sequences between 40 and 200 residues.

#### 3. Protein structure predictions

##### 3.1. Sequence alignments

Sequences for predictions were selected following the use of the microprot pipeline (https://github.com/biocore/microprot). Briefly, for each gene in each of the selected genomes the pipeline runs HHsearch[6] to discard fragments (domains) with detectable similarity to known structures (PDB70 database).

```
% hhsearch -i seq.fasta -d pdb70 -e 0.1 -Z 2000 -B 2000 -o seq.out -oa3m seq.a3m

% python split_search.py seq.out seq.fasta -e 0.1 -l 40 -o seq_cm.fasta
```

Then, each fragment is split further by running HHsearch against Pfam.

```
% hhsearch -i seq_cm.fasta -d pfamA -e 0.1 -Z 2000 -B 2000 -o seq_cm.out -oa3m
seq_cm.a3m
```

```
% python split_search.py seq_cm.out seq_cm.fasta -e 0.01 -p 90.0 -l 40 -o seq_pfam.fasta
```

Each putative domain is assessed as being suitable for *ab initio* predictions or not on the basis on the number of effective sequences in an alignment (Neff) following HHblits.

```
% hhblits -i seq_pfam.fasta -d UniRef30 -n 3 -e 0.01 -diff inf -cov 50 -Z 10000 -B 10000 -o
seq_pfam.out -oa3m seq_pfam.a3m

% python calculate_Neff.py -i seq_pfam.a3m -c 80 -o seq_msa.neff
```

Within the lifetime of the Microbiome Immunity Project (Aug 2017 - Jun 2021) reference databases were consistently updated to reflect up-to-date changes.

##### 3.2. Fragment picking

Rosetta's *ab initio* structure prediction protocols use a fragment assembly approach of 3 to 9 residue peptide fragments extracted from experimentally determined structures in the PDB and that have similar sequences and/or predicted properties to the target sequence. Protein fragments capture local structure and can speed up the search of conformation space of the target sequence being predicted. We created fragment libraries using the standard protocol[7]. The fragment picking protocol makes use of NCBI BLAST[8] (version 2.2.17) and the NCBI Non-Redundant database (downloaded Aug 2017) for alignments and sequence profiles, SPARKS-X[9] and PSI-PRED[10] (version 3.3) for secondary structure prediction and Rosetta (version ac6fa1beb57c70914b419b2e07450f1add8b81f4) for picking fragments. For each of the MIP sequences, 200 fragments of length 3 and 9 residue were generated per position. The fragment picking tools are part of the standard Rosetta distribution package and require only the target sequence. Fragments are generated using a command like below.

```
% perl make_fragments.pl MIP_${KEY}.fasta
```

##### 3.3. Residue-residue contact predictions

The accuracy of protein structure prediction can be improved by incorporating restraints from residue-pair interactions into the structure prediction trajectory, which are inferred from evolutionary couplings. We inferred residue-residue contacts from sequences closely-related to the target sequence using GREMLIN[11] (version 2.0.1). GREMLIN takes as input a target sequence and a multiple sequence alignment (described above) and outputs scores representative of the likelihood of a contact for each residue-pair in the target sequence. Multiple sequence alignments were filtered using hhfilter, converted to PSICOV format[12] using the HH Suite reformat script, and then GREMLIN was run with the default values using the command lines below.

```
% hhfilter -id 90 -cov 75 -i ${A3M_PATH} -o MIP_${KEY}.a3m

% perl reformat.pl a3m psi MIP_${KEY}.a3m MIP_${KEY}.psi -l 10000 -M first -r

% run_gremlin.sh /PATH/TO/MATLAB_Runtime/v717/ MIP_${KEY}.psi MIP_${KEY}.mat
```

The raw GREMLIN score matrix was converted into a format that can be used in Rosetta: Raw scores between residue-pairs with a sequence separation of less than 3 were removed. Raw scores from all remaining residue-pairs in the score matrix were averaged. Individual residue-pair scores were scaled by dividing by the average score and residue-pairs with a scaled score less than 2.00 were removed. The Rosetta *ab initio* structure prediction protocol does not require all residues to be restrained and only the most confident predictions were used. The remaining scaled scores were then sorted and only the top scoring contacts (the number of contacts used was up to 1.5 times the sequence length of the target sequence) were converted into AtomPair constraints for the Rosetta constraint framework using the form described below (https://new.rosettacommons.org/docs/latest/rosetta_basics/file_types/constraint-file).

```
AtomPair {atomname_i} {res_i} {atomname_j} {res_j} SCALARWEIGHTEDFUNC
{scaled_score_ij} SUMFUNC 2 SIGMOID {nbr_dist_ij} 3.0 CONSTANTFUNC -0.5
```

The distance between residue-pairs *i* and *j* in the structure prediction trajectory was constrained using a sigmoidal function between the “neighbor atoms” of each residue (Cβ atom except for glycine where Cα was used). The functional form of the constraint was

$$f_{sigmoid}(x) = 1/(1 + exp(-m \cdot (x - x_{0,ij}))) - 0.5$$

where *m* is the slope (fixed to 3.0 for all atom-pairs), *x* is the distance between the neighbor atoms of residues *i* and *j*, and $x_{0,ij}$, the center of the sigmoid function, is the sum of the “neighbor distance” for the two residue types. The neighbor distance is the radius of the sphere that represents the side-chain in Rosetta’s centroid residue representation (neighbor atoms and distances for all residue types are shown in table below). The sigmoid function is reduced by a constant 0.5, so that it is bounded [-1,0), and then multiplied by the scaled score of the residue-pair. In contrast to harmonic or bounded constraints, sigmoid constraints are smooth and differentiable, and allow residues to slide into contact during the gradient based minimization steps of the structure prediction protocol. The flattened tails of the sigmoid function ensure that unsatisfied constraints receive minimal penalties, satisfied constraints receive an almost constant bonus relative to the confidence in the prediction and at close distance the full atom energy function to determine fine-grained interactions.

**Table S1: Neighbor distance that represents the radius of the side-chain in centroid representation around the neighbor atom.**

| Residue | Neighbor Distance (Å) | Neighbor Atom Name |
|---|---|---|
| ALA | 3.245 | Cβ |
| ARG | 5.640 | Cβ |
| ASN | 4.290 | Cβ |
| ASP | 4.250 | Cβ |
| CYS | 4.170 | Cβ |
| GLU | 4.950 | Cβ |
| GLY | 4.900 | Cα |
| GLN | 2.400 | Cβ |
| HIS | 4.660 | Cβ |
| ILE | 4.320 | Cβ |
| LEU | 4.760 | Cβ |
| LYS | 5.300 | Cβ |
| MET | 4.950 | Cβ |
| PHE | 4.815 | Cβ |
| PRO | 3.580 | Cβ |
| SER | 3.560 | Cβ |
| THR | 3.790 | Cβ |
| TRP | 4.910 | Cβ |
| TYR | 4.715 | Cβ |
| VAL | 3.900 | Cβ |

#### 3.4. Rosetta model building on the IBM World Community Grid

We predicted the structure for each MIP sequence by generating 20,000 evolutionary contact-assisted (GREMLIN) models using Rosetta, in a project deployed on the IBM World Community Grid (WCG) called the Microbiome Immunity Project (MIP). The WCG is a citizen science initiative that allows researchers to perform calculations on volunteers' computers when they would otherwise be idle. A modified version of Rosetta (based on the 2016.32.58837 release) was used to generate all Rosetta models on the WCG. The modifications were cosmetic and provided a graphical animation of the protein structure prediction process for WCG participants that ran the graphical client program. No scientific code was modified and the equivalent command line and options file (a.k.a. "flags") are shown below.

```
% minirosetta.default.linuxgccrelease @/PATH/TO/MIP_${KEY}.flags -out::suffix MIP_${KEY}
-out::file::silent -jran ${RNG_SEED}
```

Contents of the command line flags file (MIP_${KEY}.flags) are shown below.

```
-database ./database

-in:file:fasta ./MIP_KEY.fasta
-in:file:frag3 ./MIP_KEY.1
-in:file:frag9 ./MIP_KEY.2
```

```
-abinitio:relax
-relax:fast
-abinitio::increase_cycles 10
-abinitio::rg_reweight 0.5
-abinitio::rsd_wt_helix 0.5
-abinitio::rsd_wt_loop 0.5

-use_filters true
-psipred_ss2 ./MIP_KEY.psipred_ss2

-nstruct 30
-run:constant_seed

-beta_nov15

-cst_file ./MIP_KEY.cst
-cst_weight 3

-cst_fa_file ./MIP_KEY.cst
-cst_fa_weight 3
```

The protein sequence, fragment files (described above), constraints based on predicted contacts (described above), secondary structure prediction of the target sequence (described above), and a random seed are provided as inputs. The random number generator seed was to ensure that all clients received a different seed. Identical constraints were used during both the centroid and full atom phases of the protocol. The REF2015 score function[13] was used during the full-atom phase of the protocol (called beta_nov15 in this release of Rosetta). The model with the lowest energy score according to the REF2015 score function was used for function prediction and further analysis.

##### 3.5. DMPfold model building

We additionally predicted the structures of all MIP sequences using DMPFold[14]. In contrast to fragment-based protein structure prediction techniques like Rosetta, DMPFold uses a consensus neural network to predict geometric constraints from the target sequence and uses them for structure calculations. Specifically, DMPFold takes as input a target protein sequence and generates multiple sequence alignments using HHBlits on the uniclust30 database. In this work, we instead used the alignments that were used for contact prediction for Rosetta. Then, DMPFold uses a neural network to generate geometric constraints, which are then used in an iterative structure prediction pipeline using the Crystallography and NMR System (CNS), similarly to how NMR data are used for structure calculations. DMPFold produces up to 5 ranked structural models.

Representative command lines for both steps are shown below. The “map” and “21c” files are intermediate files generated in the alignment step. DMPFold was run with the default 3 iterations and 50 models per iteration. The best ranking model, “final_1.pdb” was used for function prediction and further analysis.

```
% csh seq2maps.csh MIP_${KEY}.fasta

% bash run_dmpfold.sh MIP_${KEY}.fasta MIP_${KEY}.21c MIP_${KEY}.map models 3 50
```

#### 4. Function prediction

To further understand the sequence, structure, and functional relationships in our dataset, we predicted the function of the lowest energy Rosetta models and highest-ranking DMPFold models for all MIP sequences using DeepFRI[15]. DeepFRI uses a graph convolutional neural network to predict protein function from sequence and/or structure, yet does not use homology transfer to accomplish the predictions. It has been trained to predict both Gene Ontology (GO) and Enzyme Commision (EC) identifiers. DeepFRI's prediction performance on predicted protein models is similar to ones on experimentally determined structures. Additionally, DeepFRI can be used for class activation mapping to indicate which residues are the most salient for the predicted functions. DeepFRI takes protein structures (from which it extracts the protein sequence and constructs a residue-residue pair contact maps), and a set of trained weights as input. It outputs a vector of scores where each element in the vector is assigned to a specific function and scores range from 0 to 1. This means that each protein has multiple different functions that don’t compete with one another.

In this work, we used an updated set of weights trained on the SIFTS 2021 database release called the “newest trained models” available (https://github.com/flatironinstitute/DeepFRI). Generating class activation maps increases the runtime and was not performed on the full dataset. The command lines for batch prediction on a directory of PDB formatted protein structures are shown below. Each of the top level GO classifications, and the EC identifiers need to be run separately; molecular function (mf), cellular component (cc), biological process (bp), and enzyme commission (ec).

```
% python DeepFRI/predict.py --verbose --model_config
newest_trained_models/model_config.json --saliency --use_guided_grads --verbose --ont
{cc,mf,bp,ec} --pdb_dir /PATH/TO/PDB_DIR
```

The confidence metric for DeepFRI should be interpreted in the following way: DeepFRI scores > 0.5 have a high confidence, which is mostly due to high occurrences in the training dataset and which often happens for more general functions at the top of the GO hierarchies. DeepFRI scores < 0.2 don't necessarily indicate an incorrect prediction, but rather that the number of examples in the training dataset is low. DeepFRI is further trained such that scores propagate along the GO tree with higher scores towards the top of the tree and lower scores at the leafs, meaning that DeepFRI produces self-consistent predictions and does not "randomly" sample the GO tree. We often find that DeepFRI scores of 0.2 or even lower are perfectly adequate as indicated by very similar predictions for vastly different sequences (yet same structures), which is another indication of confidence in our predictions (see Fig. 4).

For functional similarity, we used cosine similarity between DeepFRI vectors thresholded at 0.1 (the non-zero threshold is important for denoising). DeepFRI was developed such that output scores are normalized between 0 and 1 and cosine similarity of two DeepFRI score vectors is a normalized sum of products of positive values (and therefore cannot be negative). We used cosine similarity as a similarity metric of two function prediction vectors because proteins have multiple functions and therefore DeepFRI predictions are vectors in a multi-dimensional function space. If two proteins have the same vectors, they would have the same function. If two proteins have the same functions but with slightly different scores, the function vectors would have a similar direction in that multi-dimensional space, therefore using cosine similarity as a similarity metric makes intuitive sense. In contrast, if two proteins have very different functions, the elements with non-zero scores in the two vectors would differ and therefore the cosine similarity would be zero or close to zero, meaning the directions of the score vectors in the multi-dimensional space would be different.

#### 5. MIP dataset descriptions

Datasets were created in the following order:

(1) *MIP_raw* is the starting dataset with all entries, containing both Rosetta and DMPFold models of varying quality from both high-quality to low-quality models.
(2) We used 5,000 random entries from the *MIP_raw* dataset to derive model quality metrics – this dataset is denoted *MIP_random5000_raw*.
(3) When filtering out the low-quality models from *MIP_random5000_raw*, we are left with 3,052 high-quality models, denoting *MIP_random5000_curated*
(4) When filtering out the low-quality models from the entire *MIP_raw* dataset, we get the *MIP_curated* dataset that only contains high-quality models.
(5) We also created a subset of *MIP_curated* for visualization purposes, these are 10,000 pairs of high-quality Rosetta and DMPFold models.

The tables below describe our datasets and the number of models in each dataset. Note that there is partial overlap between the datasets of Rosetta models and DMPFold models because either DMPFold didn't converge, or Rosetta models weren't completed due to interrupted runs on the World Community Grid.

**Table S2: Definitions of the MIP datasets.**

| Dataset | Description |
|---|---|
| *MIP_raw* | All MIP entries. Includes both Rosetta and DMPFold models. |
| *MIP_curated* | Highest quality dataset created by filtering *MIP_raw* by quality metrics. Used for most quantitative studies in the main paper and the supplement. |
| *MIP_visualization* | Representative dataset from *MIP_curated* for visualization purposes; randomly sampled 9839 MIP IDs and added 161 representatives of novel fold clusters (in total: 10,000 points for DMPFold + 10,000 for Rosetta + 6631 CATH representatives). |
| *MIP_random5000_raw* | 5000 random entries from *MIP_raw* with IDs common between Rosetta and DMPfold sampled by length distribution. The dataset was used to estimate the quality metric thresholds for constructing the *MIP_curated* dataset. |
| *MIP_random5000_curated* | The curated part of MIP random5000_raw was used to compute pairwise similarity measures in Fig. 2. |

**Table S3: Number of structures in the MIP datasets.**

| | Rosetta models | DMPFold models | CATH superfamilies | Common Rosetta-DMPfold models |
|---|---|---|---|---|
| *MIP_raw* | 245,443 | 241,834 | 0 | 240,703 |
| *MIP_curated* | 211,069 | 203,877 | 0 | 184,642 |
| *MIP_visualization* | 10,000 | 10,000 | 6,631 | 10,000 |
| *MIP_random5000_raw* | 5000 | 5000 | 0 | 5000 |
| *MIP_random5000_curated* | 3052 | 3052 | 0 | 3052 |

#### 5.1. MIP dataset curation

To identity high quality models in the *MIP_raw* dataset we used the following quality measures (MQA - model quality assessment):

- Rosetta: average TM-score between the 10 structures with the lowest energies
- DMPfold: quality metric of the structural model that describes the agreement between the neural-network predicted contact map and the contact map back-computed from the final model

The metrics, together with other parameters (e.g. Stride[16] output), were then analyzed (independently for Rosetta and DMPFold) using the Random5000_raw dataset - see section *Random5000 data analysis* below for details. As a result, the *MIP_curated* dataset was constructed based on the following criteria:

- Rosetta: MQA >= 0.4 and coil content ≤ 60%
- DMPFold: MQA >= 0.4 and coil content ≤ 80%

The coil content was computed as a sum of T and C Stride annotations normalized by sequence length. In Fig. S1 we show how the above criteria shift the density distribution between *MIP_raw* and curated i.e. we can notice higher agreement between Rosetta and DMPFold models for the latter.

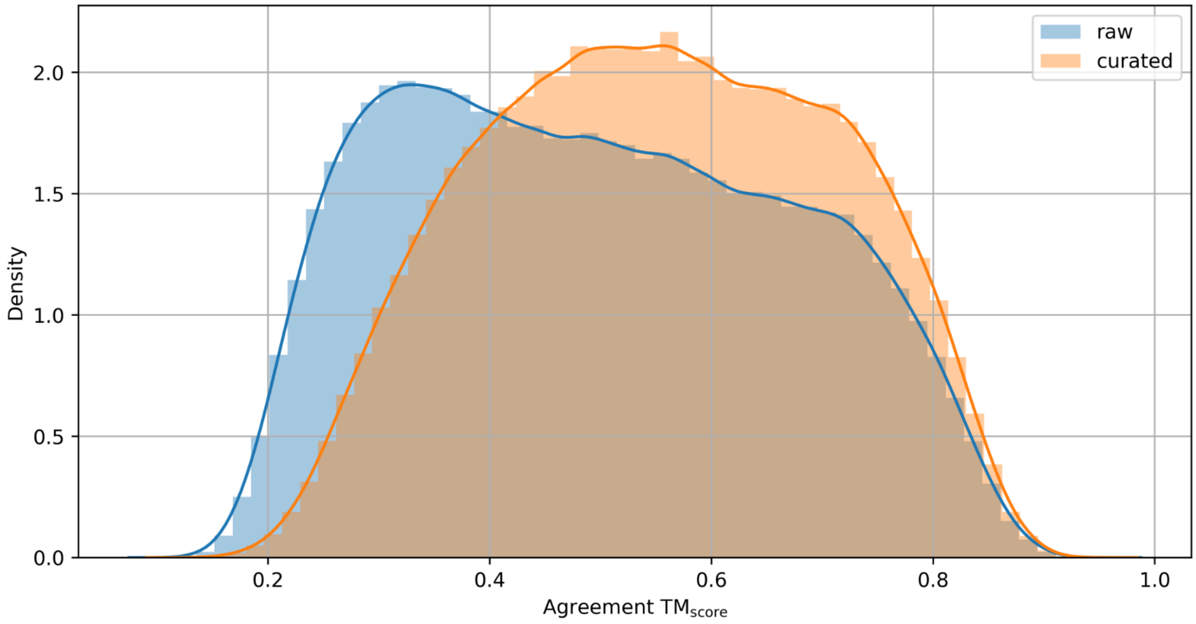

*Fig. S1. Density plot of MIP structures as a function of structural similarity (agreement TM-score between Rosetta and DMPFold models) for MIP_raw and MIP_curated.*

##### 5.2. Sequence length distribution

MIP sequences are relatively short (between 40 and 200 residues) - see upper panel in Fig. S2. Comparing sequence lengths distributions for *MIP_curated* (lower panel in Fig. S2) we see that more short DMPFold models (<100 residues) were discarded, as were more longer Rosetta models (>100 residues), compared to the *MIP_raw* dataset. However, the overlap (common part) of the models that survived is still high (89%) - see also Table S3.

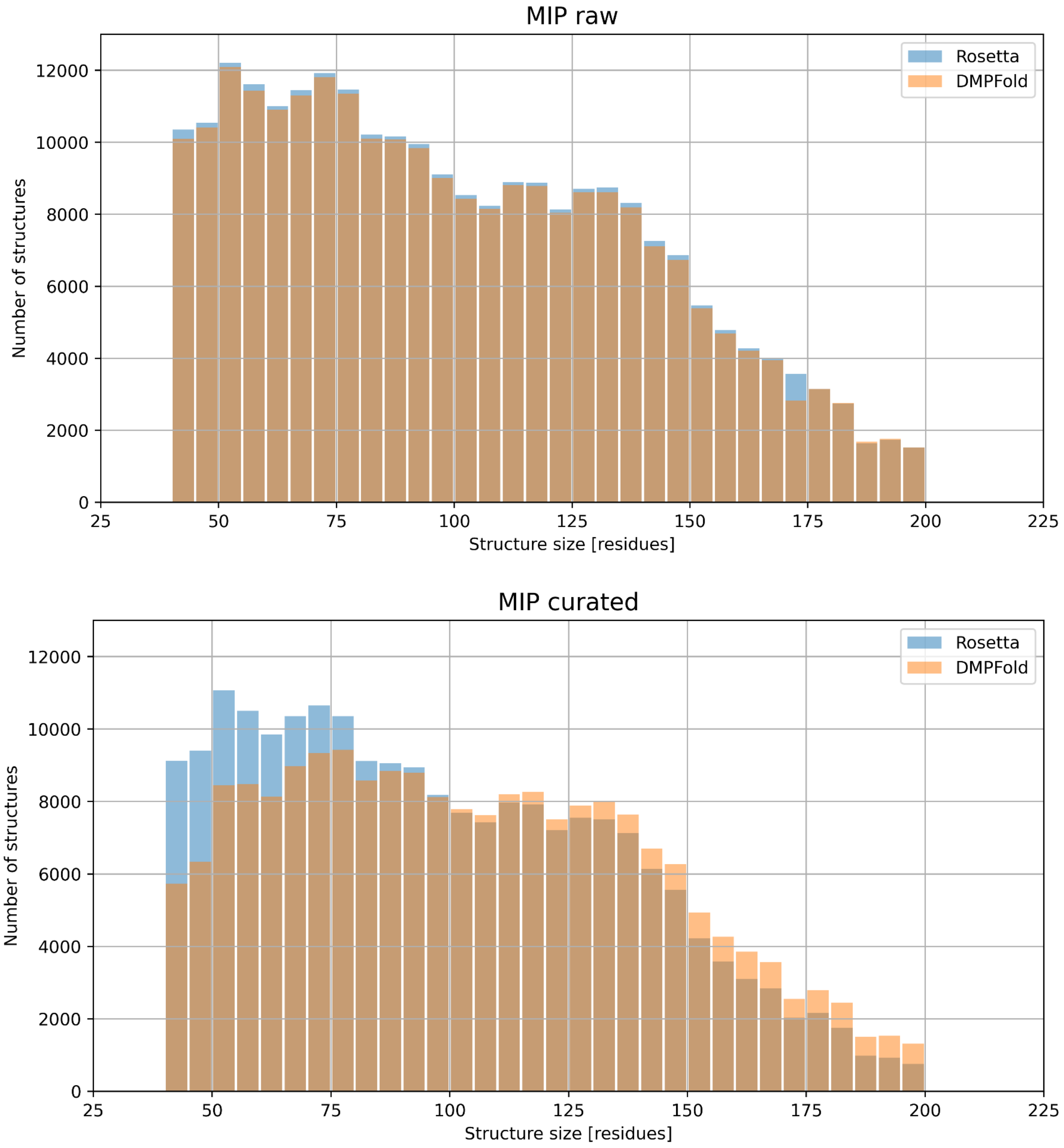

*Fig. S2. Sequence length distributions for MIP_raw and MIP_curated.*

#### 5.3. Pairwise structural comparisons in the curated dataset via CATH

Our next goal was to identify how structurally similar any two MIP models were to each other, to find clusters of similar structures and to find novel folds. However, this would have required 200,000 x 200,000 / 2 = 20 billion comparisons which is computationally prohibitive. Instead, we decided to compare each of our *MIP_curated* models to CATH superfamilies, of which there are about 6,000, and then compare CATH superfamilies to each other. This led to 6,000 x 6,000 / 2 = 18 million comparisons, which was far more tractable.

To accomplish this, all models in the *MIP_curated* dataset were superimposed with CATH 4.3.0 superfamily structures using TM-align (see Methods section in the main paper). For each model we then found the most similar CATH representative (the largest TM-score normalized by MIP sequence length) and annotated the model by CATH hierarchy levels (class, architecture, fold, superfamily) assigned to that CATH structure. We found that approx. 70% of CATH superfamilies were identified as the closest structural homologs for the Rosetta curated dataset and approx. 65% for the DMPFold curated set. The percentages drop slightly (to 68% and 62% respectively) when considering only structures with TM-scores >= 0.5.

Overall, Rosetta predicts more alpha-helical models than DMPFold and inversely for beta and alpha/beta models (left panel of Fig. S3). The difference even grows with TM-score (better quality of annotations). The above conclusion also holds on the architecture level (right panel in Fig. S3). For all major architectures in each CATH class we notice similar differences between Rosetta and DMPFold.

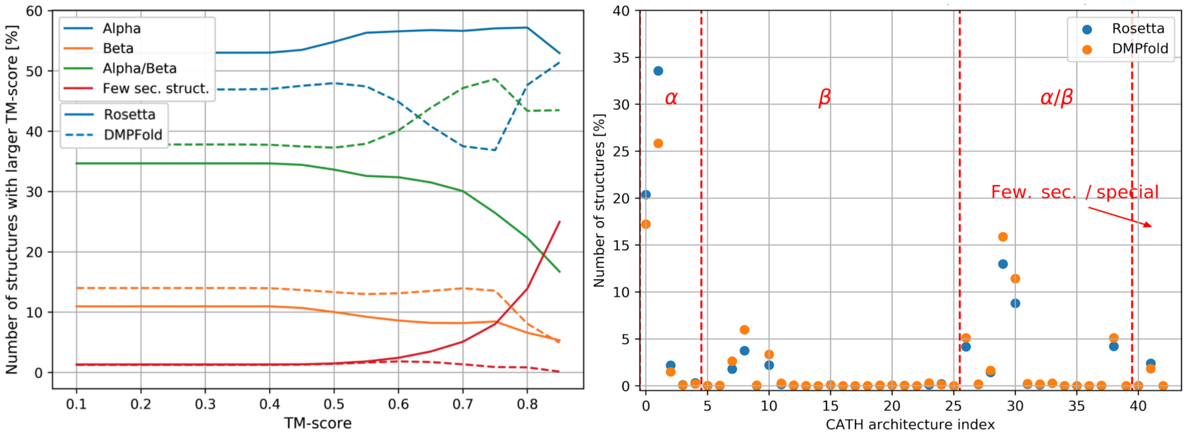

*Fig. S3: Left: Comparing the models in the MIP_curated dataset against CATH classes and plotting the TM-score over the percentage with the specific classes. Solid lines for Rosetta models, dotted lines for DMPFold models. For a TM-score around 0.7, DMPFold the percentage of models switches between alpha and alpha/beta proteins. With increasing TM-score, the percentage of Rosetta models with few secondary structure elements increases while the alpha/beta percentage decreases. Right: percentage of structures in MIP_curated datasets (Rosetta, DMPFold) for a given CATH architecture (only structures with maximum TM-score against CATH >= 0.6 were considered).*

### 6. Model quality assessment on the MIP_random5000 dataset

#### 6.1. Explanation of variables

*N_eff*

N effective is the number of sequences in the multiple sequence alignment that each MIP entry has. For contact prediction, the larger N_eff is, the higher quality the predicted contacts and therefore the models are. It has previously been suggested that N_eff has to be at least 5x the sequence length to produce high-quality contacts during prediction. However, we use the lower limit of 16 sequences required to predict contacts using GREMLIN[11].

*Sequence length*

The raw sequences that came out of sequencing were split into domains based on sequence-based predictions. These domains were given a MIP ID entry and their sequence length was based on this domain sequence.

*Secondary structure*

The sequence for each MIP ID was used for structure prediction both with DMPFold[14] and Rosetta *abinitio* modeling using contact restraints predicted from GREMLIN[11]. Both DMPFold and Rosetta models were run through Stride[16] to extract the secondary structure definitions for each residue. Stride classifies residues into the following secondary structures: H – alpha helix; G – 3-10 helix; I – PI-helix; E – extended conformation; B or b – isolated bridge; T – turn; C – coil (none of the above). To create a 3-state secondary structure definition, we grouped H, G, and I into a helix definition; E and Bb into strand definition; and defined (1 - helix - strand) as coil. Coil percentage is defined as the number of coil residues over the sequence length; similar for helix and strand.

*Membrane content*
The sequence for each MIP ID was used to predict transmembrane helices through OCTOPUS[17] and transmembrane strands through BOCTOPUS[18]. The membrane-helix percentage is the number of residues predicted as a transmembrane helix divided by the sequence length; similar for membrane-strand prediction.

*Disorder*
Protein disorder was predicted from the sequence of each MIP entry using MobiDB-lite[19] v.1.0 (March 2016) and DISOPRED3[20] (zip downloaded on September 16, 2021; dependencies: Blast v.2.2.26, Uniref90 v.20210731).

*TM-score to CATH*
Predicting the structures of millions of proteins raises the question of how many novel folds are predicted in this dataset. This requires each new model to be compared with known protein structures. CATH[21] is a database that contains all known non-redundant protein domains present in the Protein Data Bank[22]. An advantage is the small size of the database encompassing ~6000 domains, as compared to ~150,000 structures in the PDB, minimizing runtime. We use the TM-score[23] ('template modeling score') as a structural similarity measure which falls between 1 for identical folds, to 0 for different folds. From a technical perspective, the TM-score rarely falls below 0.2 and a cutoff of 0.4 is often used to distinguish novel folds (TM-score ≤ 0.4) from known folds (TM-score > 0.4). The TM-score is computed as

$$\text{TM-score} = \max\left(\frac{1}{L_{target}} \cdot \sum_{i}^{L_{aligned}} \frac{1}{1 + \left(\frac{d_i}{d_0(L_{target})}\right)^2}\right)$$

where $L_{target}$ and $L_{aligned}$ are the lengths of the protein target and the aligned regions, respectively. $d_i$ is the distance between the $i$-th pair of residues and

$$d_0(L_{target}) = 1.24 \cdot \sqrt[3]{L_{target} - 15} - 1.8$$

is a scaling factor that normalizes distances.

*Agreement TM-score*
Because we have two models for each MIP ID, a DMPFold and a Rosetta model, we can compute a TM-score of the Rosetta model (aligned) to the DMPFold model (target) and vice versa. This leaves us with two TM-scores, the average of which we define as the agreement TM-score. Like the TM-score, the agreement TM-score falls between 0 and 1, with 1 being ideal model structural agreement.

*Contact order*

The contact order is a measure of fold complexity. The absolute contact order is computed as

$$Absolute\ contact\ order = \frac{1}{N}\sum^{N} \Delta S_{ij}$$

with $N$ being the total number of contacts and $\Delta S_{ij}$ being the sequence separation between residues $i$ and $j$. The relative contact order is the absolute contact order normalized by the sequence length, therefore being length independent.

*MQA score*

We sought a model quality assessment score that is fast to compute, is ideally a single model score and tells us something about how ‘good’ the overall model is, irrespective of details such as bond lengths, bond angles, rotamers etc. We considered the VoroMQA score, yet ultimately refrained from using it for reasons mentioned below. We decided on a MQA score that is dependent on the method used. We know that Rosetta is often able to find the global minimum of an energy landscape, especially looking at the top 10 (i.e. 10 lowest-energy) models from 20,000 models (a.k.a. ‘decoys’). Even if the top model belongs to a false minimum on the energy landscape, we know from experience that some models among the top 10 are close to the ‘native’ structure if an experimental structure exists. In the vast majority of cases, clustering the top 10 models and looking at the largest cluster, will find the ‘correct’ solution. While this exact approach is somewhat cumbersome and computationally inefficient, we use a shortcut which we show below seems to work just as well: we compute the average of the pairwise TM-scores between the top 10 Rosetta models. Again, this TM-score is in the range between 0 and 1. The higher this Rosetta MQA score, the more of the top 10 models have the same structure, supporting the notion that Rosetta has found the global minimum in this case. Therefore, the MQA score also encodes our confidence in the correctness of the Rosetta model.

DMPFold is unfortunately inconsistent and outputs 1-5 final models, however the models have a confidence score associated with them. We use this confidence score as MQA score for the DMPFold model.

*Rosetta score*

Each Rosetta model has a score associated with them which represents the ‘energy’ of the model in the energy landscape of the Rosetta scoring function. The total score, which we are using here, is a weighted combination of different score terms, encoding both physics-based and knowledge-based terms. The total Rosetta score is not normalized by protein length. A detailed review of the Rosetta scoring function was recently published[13].

#### 6.2. DMPFold models have higher coil content than Rosetta models

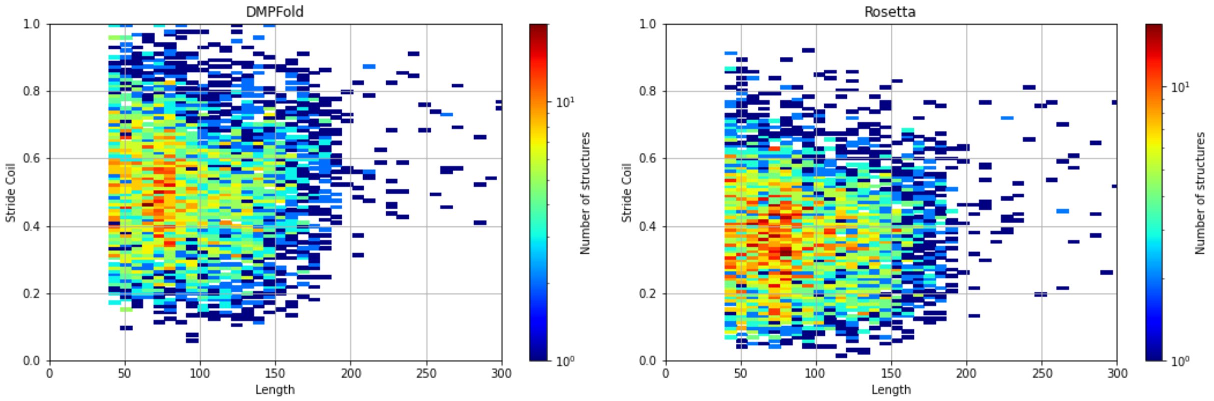

*Fig. S4: Percentage of coil residues in the models, identified by Stride, over the sequence length. DMPFold models have a higher percentage of coil residues due to local inaccuracies in backbone torsion angles. DMPFold uses evolutionary restraints from contact prediction and predicts the structure using the CNS software under the influence of a molecular dynamics force field. Visually analyzing a large number of models indicates that Rosetta's fragment-based approach (in conjunction with contact constraints) and its scoring function are able to produce better local features in these models, while DMPFold is able to produce higher quality models of larger proteins.*

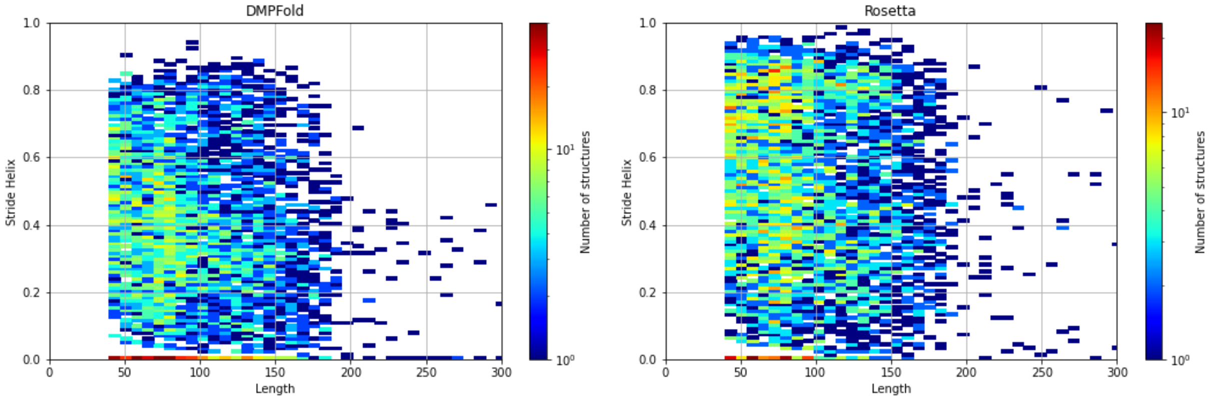

*Fig. S5: Percentage of helical residues in DMPFold and Rosetta models over the sequence length. Rosetta models generally have a higher percentage of helical residues than DMPFold models.*

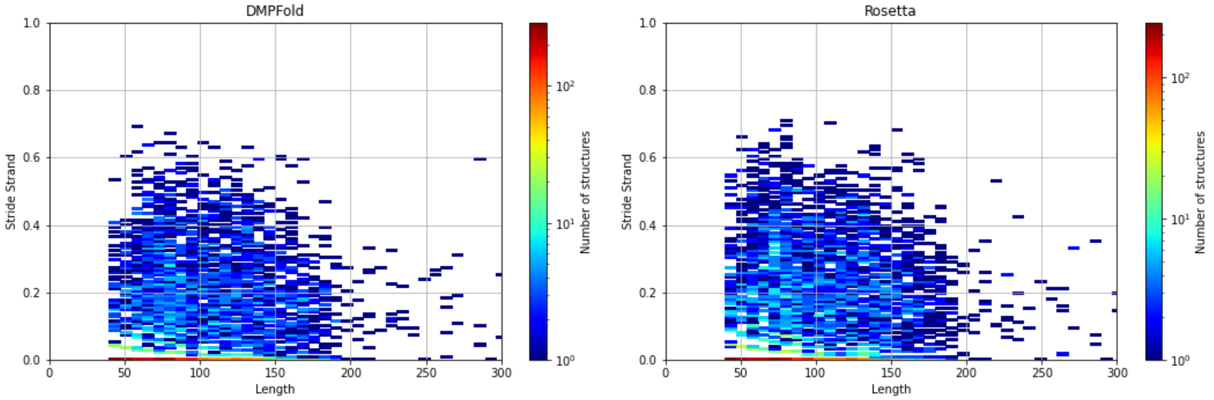

*Fig. S6: Percentage of strand residues in DMPFold and Rosetta models over the sequence length. Rosetta models have a slightly higher percentage of strand residues than DMPFold models.*

##### 6.3. Model quality is independent of sequence length and correlates with agreement between Rosetta and DMPFold models

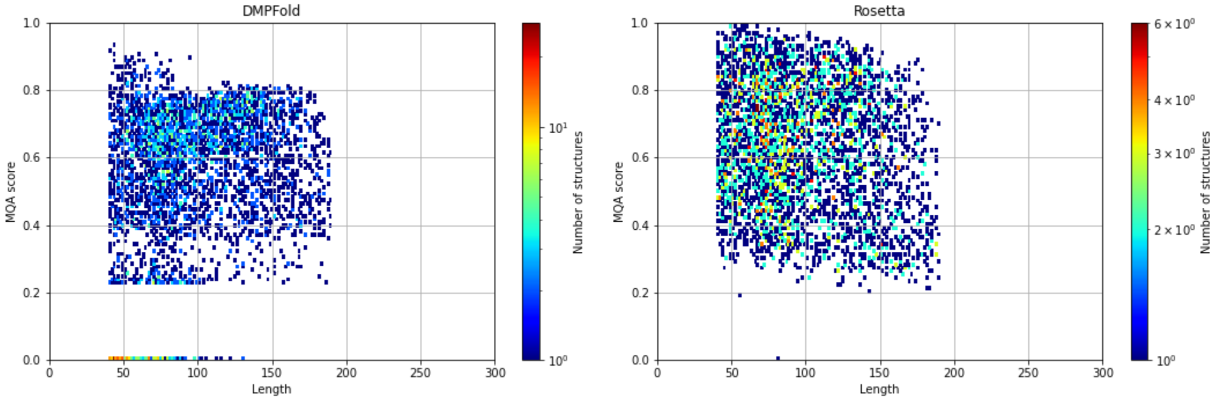

*Fig. S7: The MQA score is largely independent of protein length. While there is no correlation for DMPFold models, the MQA score limits slightly decrease for Rosetta models. Still, this lack of correlation is a major advantage for using this MQA score in model quality assessment, in contrast to, for example the VoroMQA score24, which we tested.*

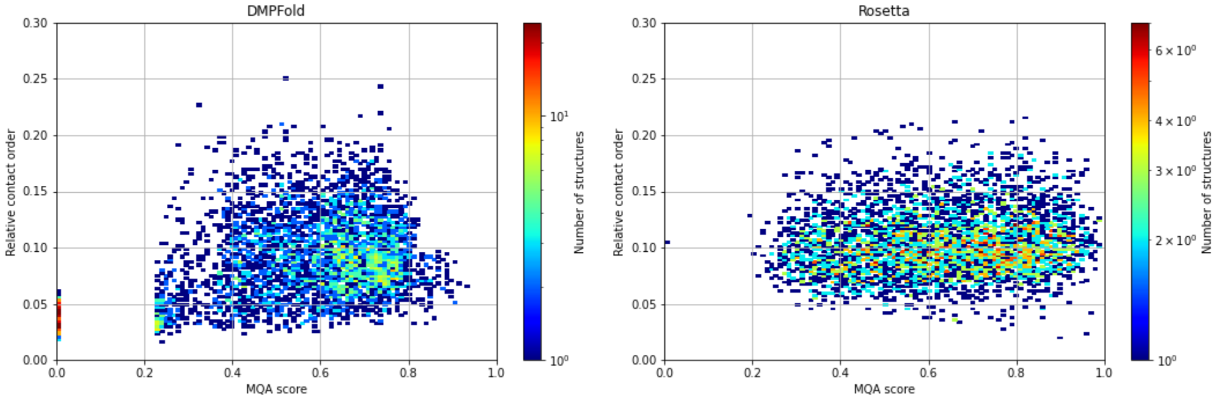

*Fig. S8: The MQA score does not correlate with relative contact order, making it independent of the protein size. This is another indication that the MQA score is a good measure to evaluate model quality.*

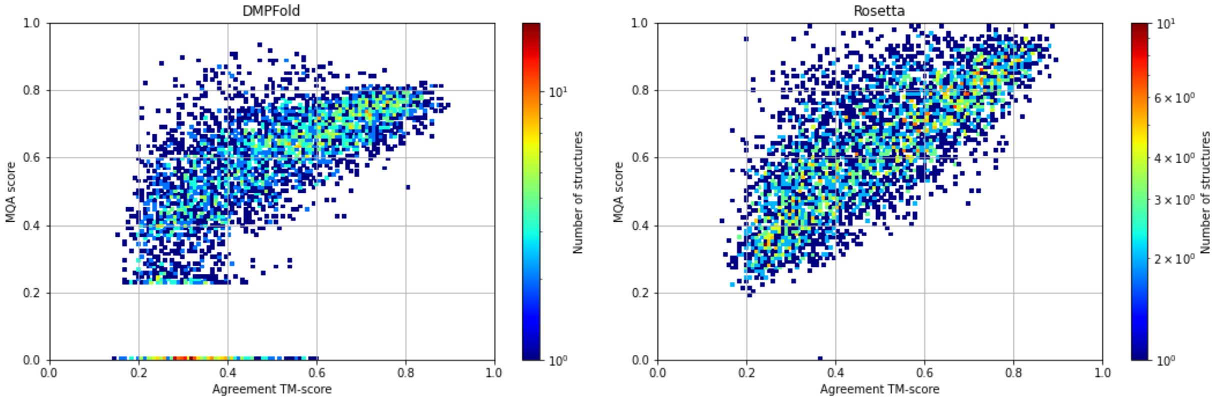

*Fig. S9: The agreement TM-score between Rosetta and DMPFold models correlates with the MQA score, making the latter a good metric for model quality. The higher the model quality is, the higher the MQA score and the more likely it is that the DMPFold and Rosetta models are similar to each other. This works especially because the structure prediction methods use contact-predicted constraints which have a high impact on the total score and model quality.*

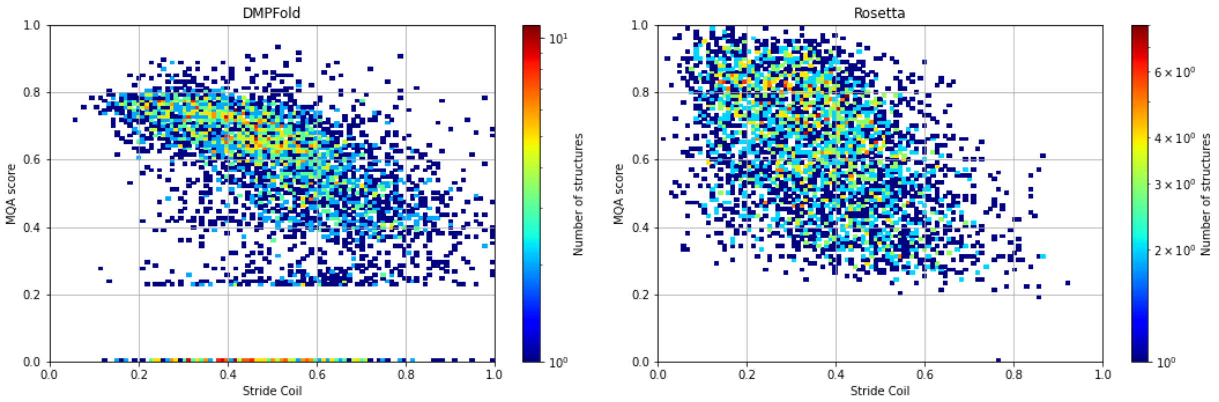

*Fig. S10: The MQA score decreases with increasing coil content because these models have less defined secondary structure.*

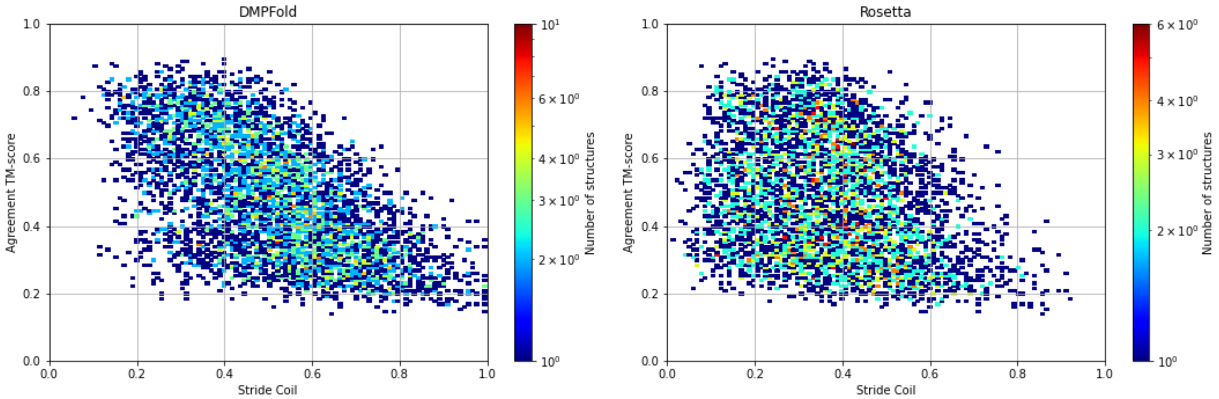

*Fig. S11: The agreement TM-score between DMPFold and Rosetta models decreases as a function of coil percentage. Models with a higher percentage of coil have worse quality and therefore less agreement between DMPFold and Rosetta.*

#### 6.4. Deeper Multiple Sequence Alignment leads to higher model quality

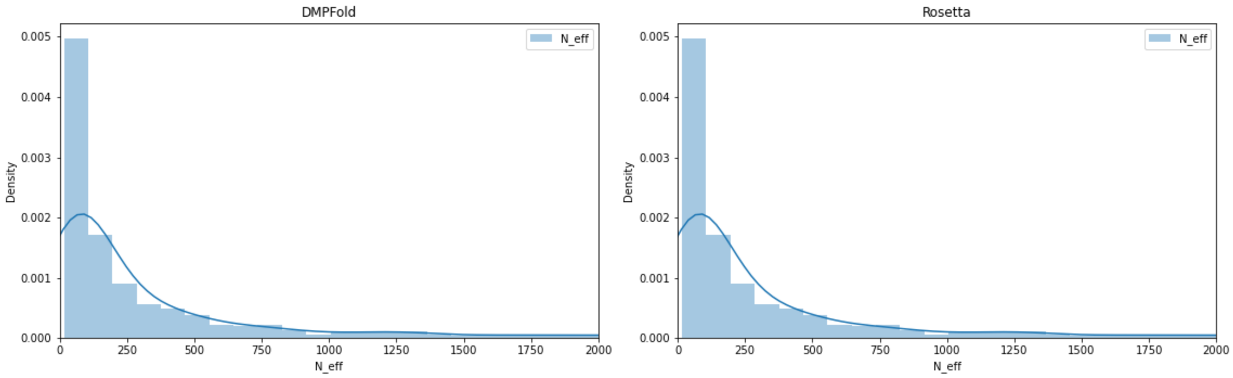

*Fig. S12: Our models are predicted de novo using restraints from contact prediction methods derived from Multiple Sequence Alignments. The majority of MIP entries has less than 250 effective sequences in their multiple sequence alignment, as indicated by N_eff.*

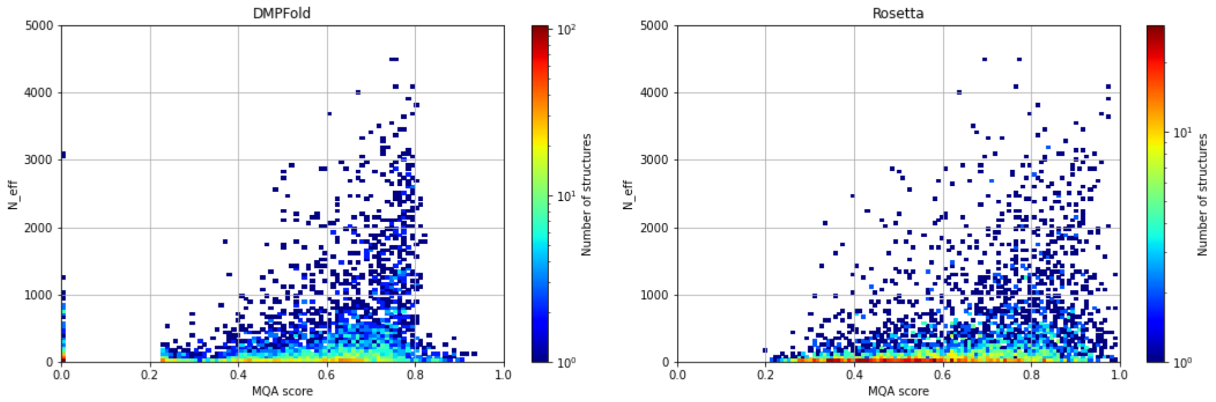

*Fig. S13: A higher N_eff leads to higher quality models for DMPFold, as indicated by its MQA score. Similarly, a higher N_eff leads to better quality models for Rosetta, using the Rosetta MQA score. Comparing both methods, the same number of sequences in the MSA leads to higher MQA scores in Rosetta as compared to DMPFold. However, MQA scores for Rosetta and DMPFold were derived independently and are therefore not perfectly comparable.*

#### 6.5. Transmembrane spans

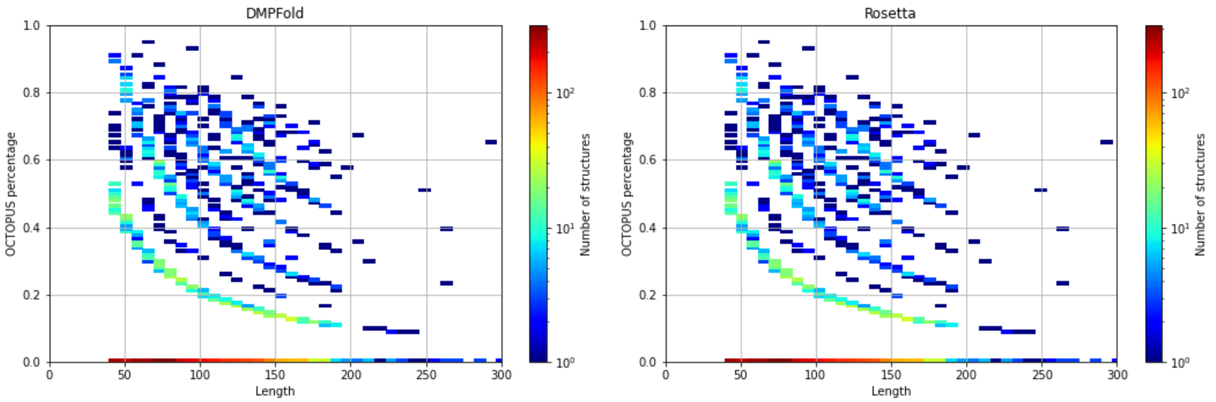

*Fig. S14: Percentage of transmembrane helix residues predicted by OCTOPUS over the sequence length. The decay of the curves is a feature of the number of transmembrane helices as normalized by the sequence length. The bottom curve is a single transmembrane helix, the curve directly above is for two helices, and so on.*

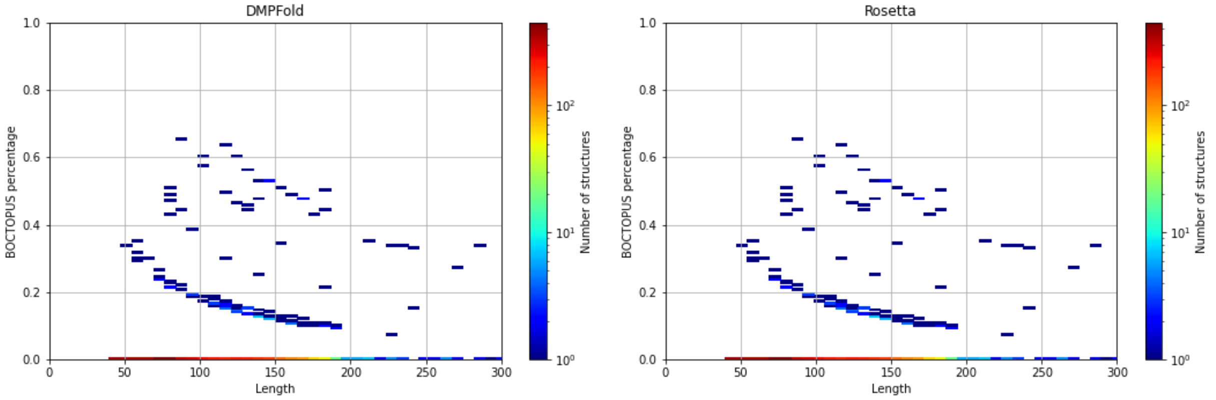

*Fig. S15: Percentage of transmembrane strand residues predicted by BOCTOPUS over the sequence length. The decay of the curves is a feature of the number of transmembrane strands as normalized by the sequence length.*

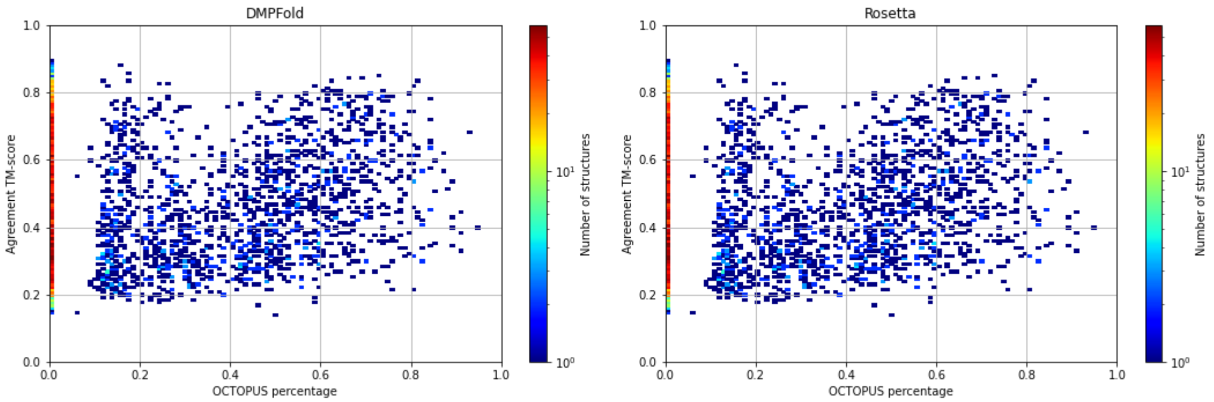

*Fig. S16: Agreement TM-score between Rosetta and DMPFold models as a function of the percentage of transmembrane helices as predicted by OCTOPUS.*

#### 6.6. Disorder

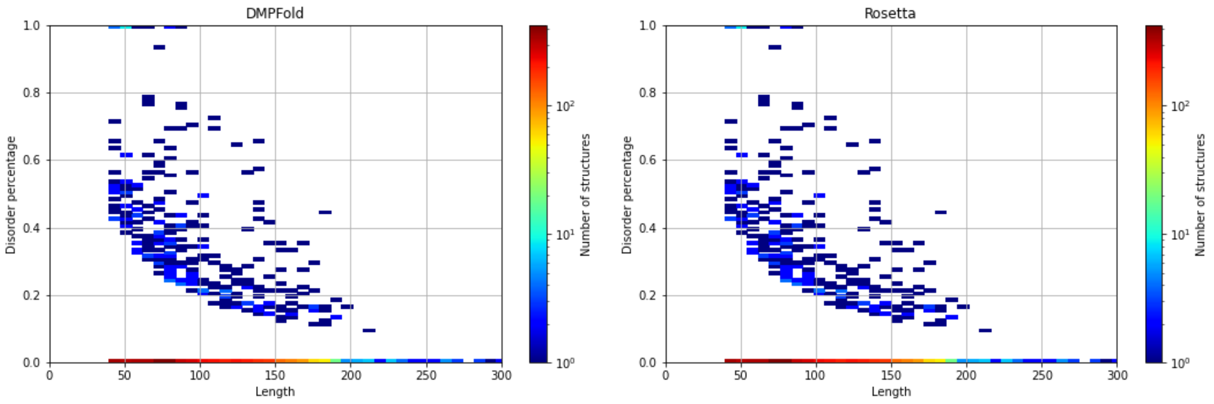

*Fig. S17: Percentage of protein disorder as predicted by DISOPRED3, plotted over the sequence length. Again, the decay of the curve is a feature over the normalization by sequence length.*

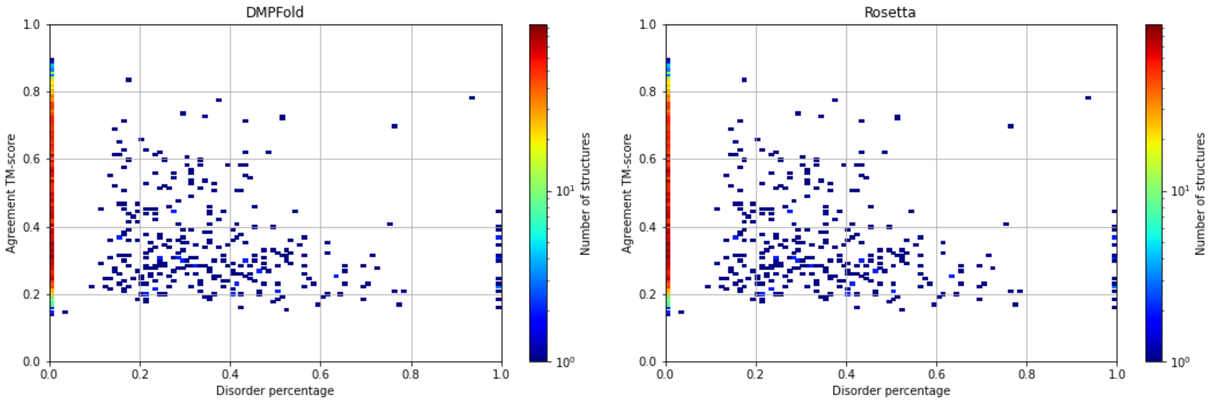

*Fig. S18: The agreement TM-score between DMPFold and Rosetta decreases with increasing disorder percentage. This originates in a larger set of possible conformations between DMPFold and Rosetta and in decreasing model quality of higher disorder conformations.*

#### 6.7. Model complexity

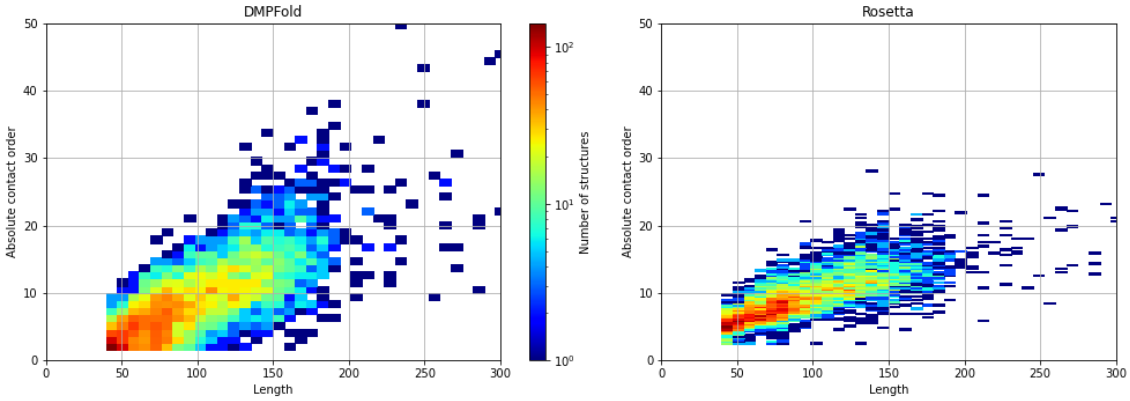

*Fig. S19: The absolute contact order correlates with sequence length. The slope is larger for DMPFold than for Rosetta because DMPFold is better at predicting larger protein models, unlike Rosetta, which was originally designed to predict small, soluble proteins. Note that the absolute contact order is length dependent.*

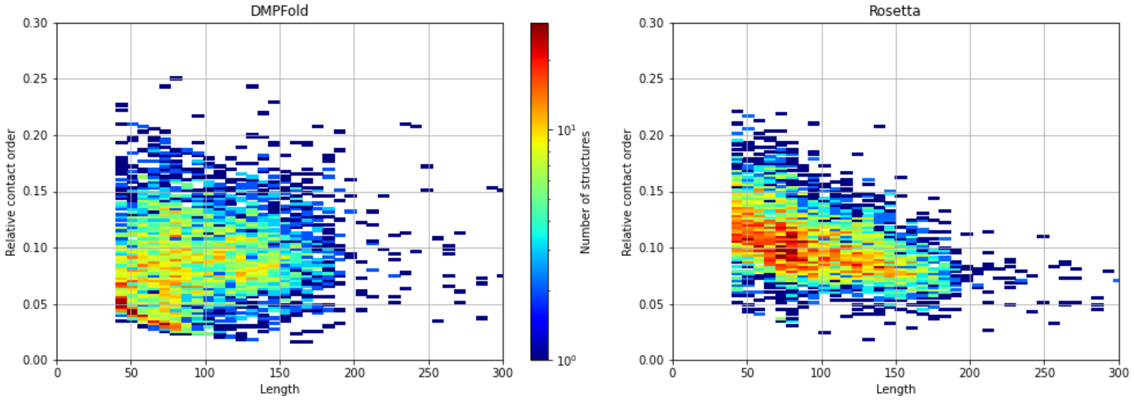

*Fig. S20: The relative contact order, which is the absolute contact order normalized by length, as a function of the sequence length. DMPFold models do not exhibit a dependence of the relative contact order on sequence length, whereas Rosetta models decrease in their relative contact order, because it is more difficult to predict larger proteins with Rosetta.*

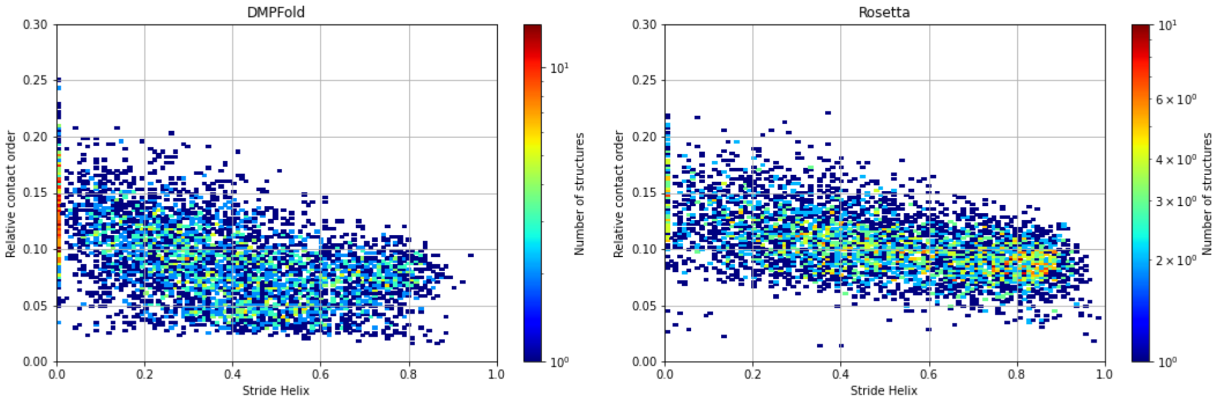

*Fig. S21: The relative contact order decreases with the percentage of helix residues. There are many proteins in the MIP dataset that have one or two helices. These have a high helical content but a small contact order. Also, contact order increases with long-range contacts, yet helices have local contacts between residues.*

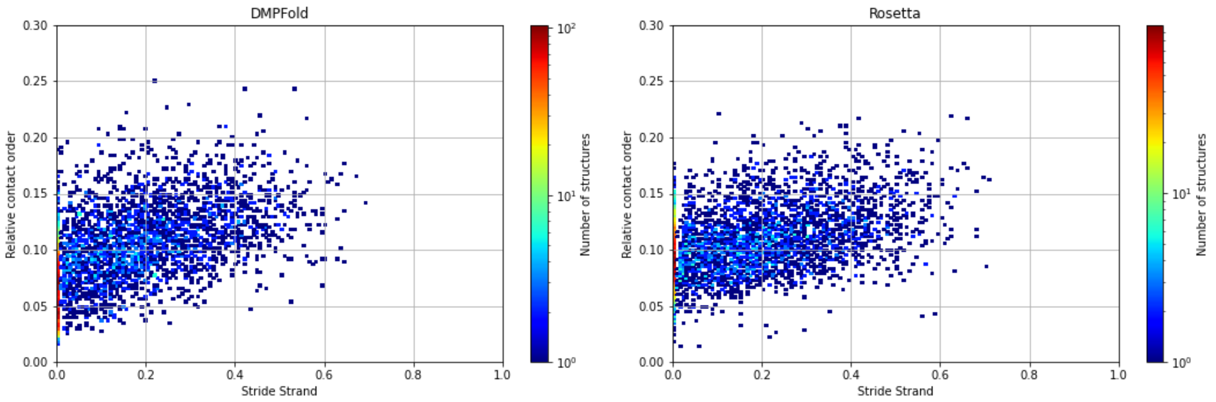

*Fig. S22: The relative contact order as a function of strand content. The higher the strand content, the higher the contact order because strands are involved in long-range contacts.*

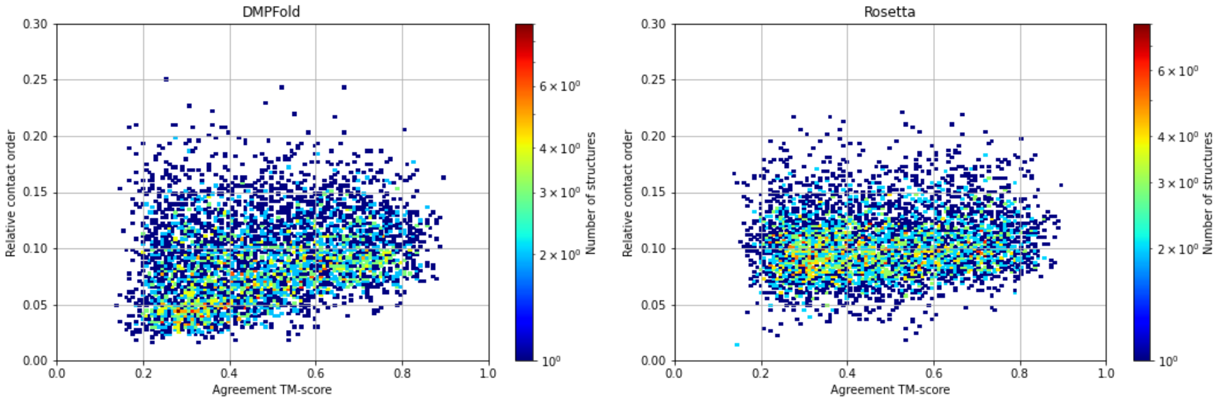

*Fig. S23: The agreement TM-score increases slightly with the relative contact order for DMPFold models, but not for Rosetta models.*

#### 6.8. TM-score to CATH

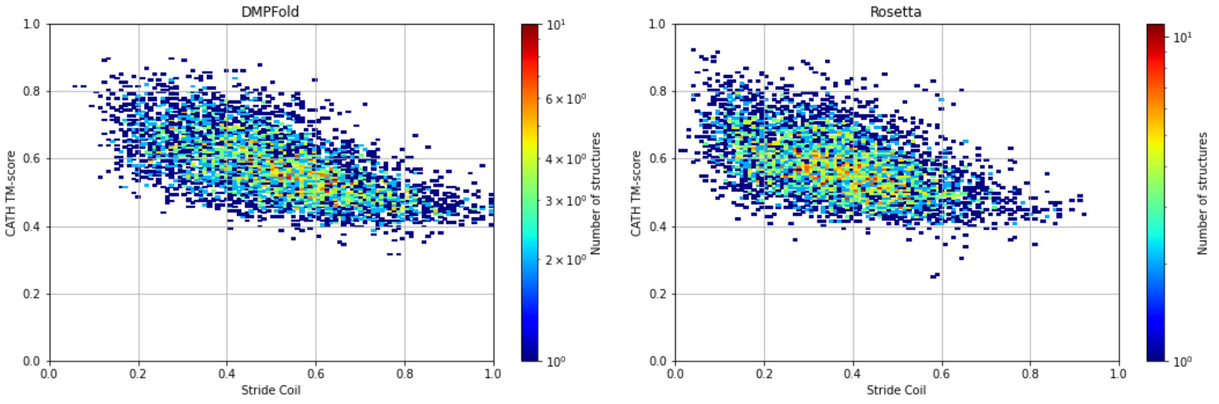

*Fig. S24: The TM-score to CATH superfamilies as a percentage of coil residues in the models, indicating a slightly negative correlation. The more unstructured the protein is, the harder it is to find a match in CATH and the lower the TM-score will be because of structural differences.*

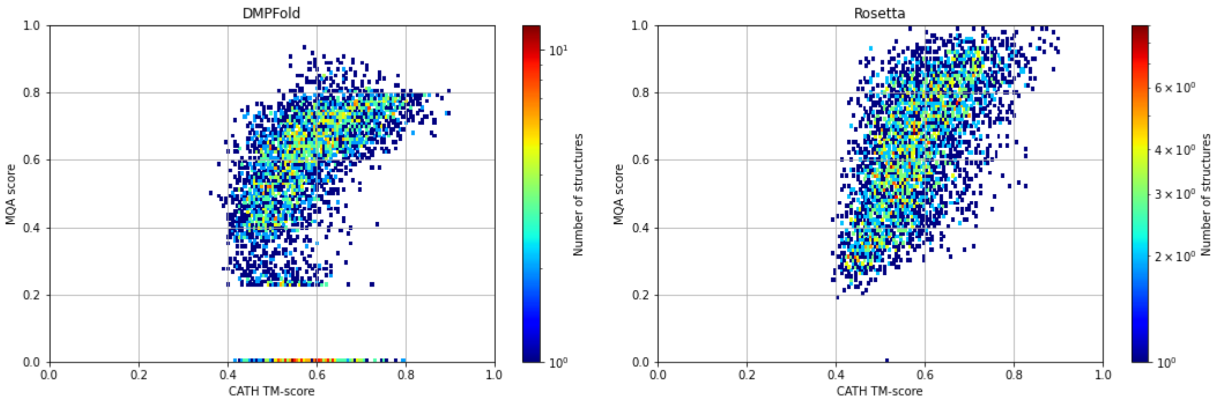

*Fig. S25: There is a small correlation between model MQA score and TM-score to CATH, meaning the higher the model quality is, the more likely it is to find a match in CATH.*

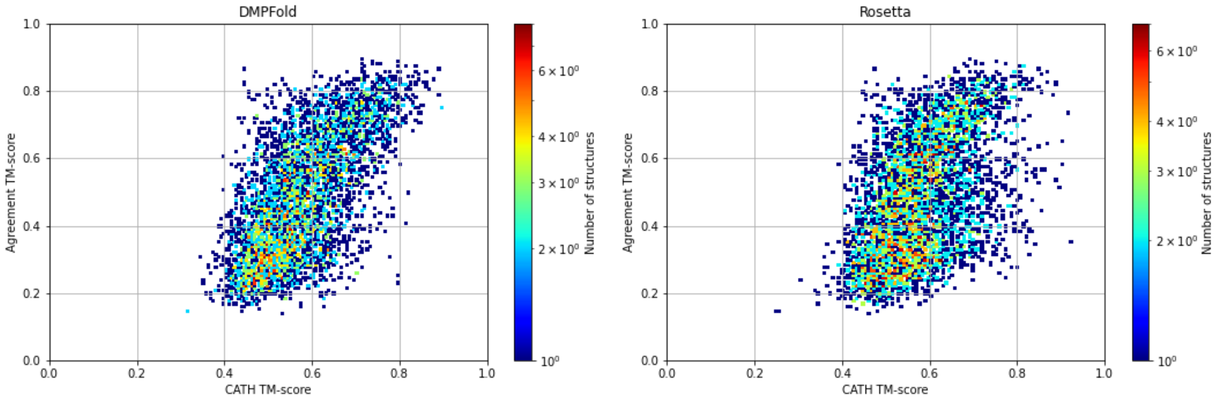

*Fig. S26: A higher agreement TM-score between Rosetta and DMPFold models correlates with a higher TM-score to a structure in the CATH database. Since agreement TM-score correlates with MQA score as well, this Fig. extends the previous one.*

#### 6.9. Rosetta score

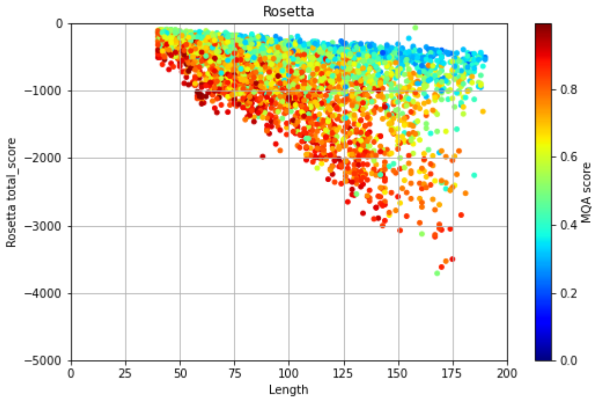

*Fig. S27: The Rosetta total score decreases for larger protein models. This is expected because the score is a sum of the scores over all residues in the protein. Further, models with the lowest (i.e. best) Rosetta scores have the highest MQA scores, which is another advantage of the MQA score.*

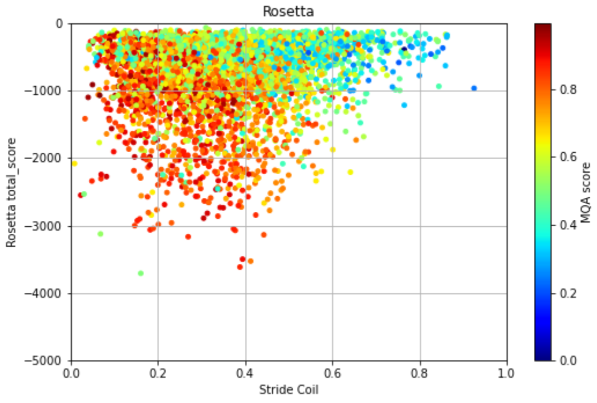

*Fig. S28: The total Rosetta score generally gets worse (i.e. increases) with increasing coil content because of lack of hydrogen bonds, optimal solvation, etc.*

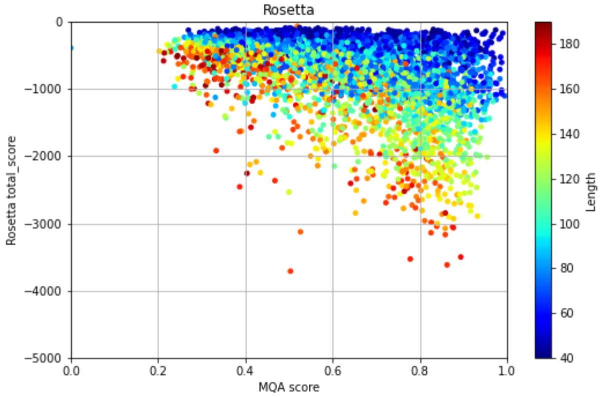

*Fig. S29: Models with higher MQA scores can achieve lower Rosetta total scores, which are typically achieved by larger proteins.*

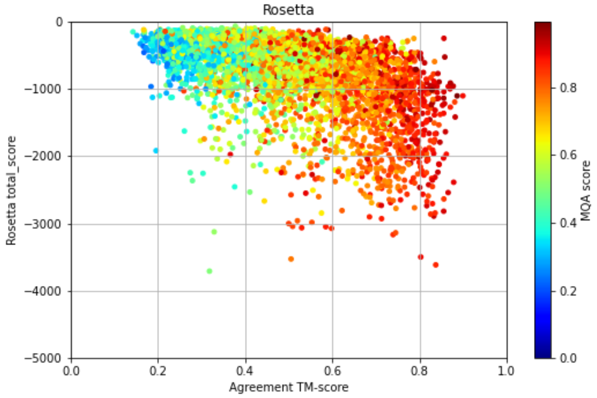

*Fig. S30: The more similar the DMPFold and Rosetta models are (high agreement TM-score), the higher the Rosetta MQA score and more likely that they can achieve a low (good) Rosetta score.*

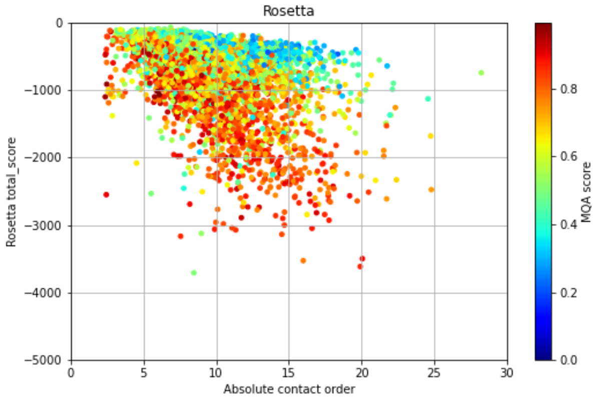

*Fig. S31: The higher the absolute contact order (i.e. the larger the proteins are), the lower the Rosetta total score these models can achieve. The lower the Rosetta total score, the better the model quality, as shown by higher MQA scores.*

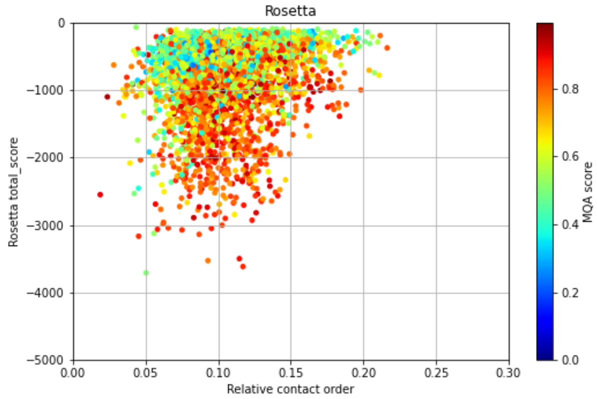

*Fig. S32: The correlation for the absolute contact order is washed out in the relative contact order, yet models on the higher end of the relative contact order typically have higher MQA scores and can have lower total Rosetta scores.*

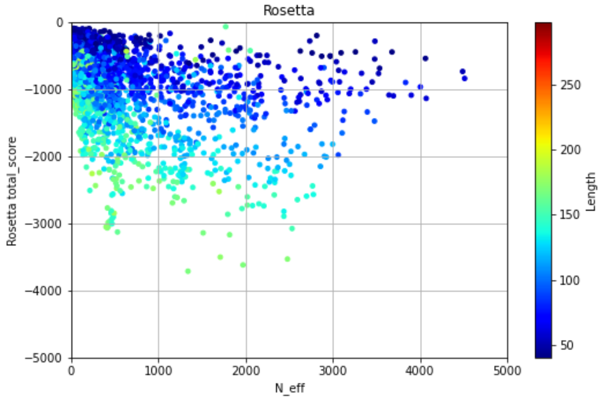

*Fig. S33: For Rosetta, a higher N_eff leads to lower total Rosetta scores, indicating higher model quality. Further, this relationship also correlates with length – longer proteins have lower Rosetta scores, because the total Rosetta score is not normalized by protein length.*

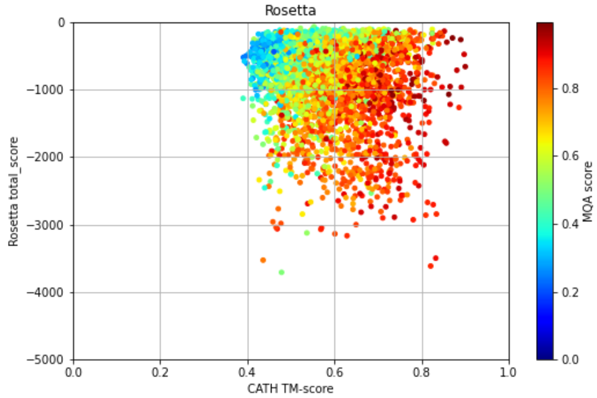

*Fig. S34: Models with a higher TM-score to CATH can achieve lower Rosetta total scores (i.e. are better models) and typically also have a higher MQA score.*

### 7. Additional results on the *MIP_curated* dataset

We performed a series of analyses including structure- and sequence-based annotations. The former include pairwise sequence and structural similarities (TM-align), function similarities (DeepFRI) and secondary structure assignments (Stride), which are already discussed in the main text and in this supplement (see MIP dataset curation, Novel folds and Sequence-structure-function relationships sections), and the contact order.

#### 7.1. Transmembrane spans

Using the protein sequences, we predicted alpha-helical transmembrane spans with OCTOPUS and beta-strand transmembrane spans with BOCTOPUS (see methods in main paper). We discovered that the percentage of alpha-helical transmembrane spans in the common *MIP_curated* proteins is equal to 33%, which is according to expectations[25,26]. In all cases the transmembrane region is located at the center of the sequence and not at the termini. As a reference, the CATH 4.3.0 superfamily dataset comprises 6.8% of helical transmembrane proteins. Interestingly, the percentage of beta barrels in *MIP_curated* common (annotated by BOCTOPUS) is much lower, around 0.3%. In this case, although still below expectations (about 3% of proteins in gram-negative bacterial genomes[27]), the number is much closer to the reference CATH 4.3.0 superfamily dataset, where we also found ~0.3% of beta barrels.

#### 7.2. Contact order

The contact order is a measure of fold complexity (see section Supplement section 6) and the relative contact order is the absolute contact order normalized by sequence length. Since our proteins are relatively small, their relative contact order is very similar to reference structures in CATH. We further observe that proteins with high beta-strand content have a larger relative contact order than alpha-helical ones (see Fig. S35).

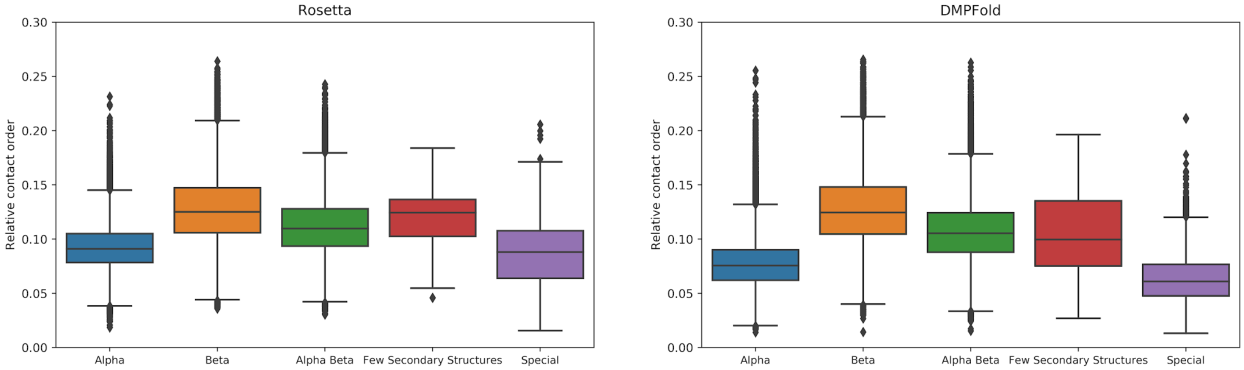

*Fig. S35: Relative contact order for MIP_curated models stratified by CATH class.*

#### 7.3. Disorder

Intrinsically disordered regions (IDRs) are important parts of proteins but their behavior is still not well understood. About 2% of archean proteins, 4% of eubacterial, and 33% of eukaryotic proteins contain IDRs longer than 30 residues[28]. To identify IDRs in the *MIP_curated* dataset we used

MobiDB-lite, which is a fast sequence-based method that, in the first step, uses 8 different predictors to derive a consensus and then filters for spurious short predictions in the second step. We found out that in the *MIP_raw* and curated datasets (considering common parts in both cases) the percentage of sequences containing at least one IDR is equal to 5.7 and 3.3 respectively, of which roughly 70% are located at the termini. The difference between raw and curated datasets is expected since the latter has less coil content and higher-quality models. However, we noticed that MobiDB-lite[19] is not sensitive enough in some cases, therefore for the putative novel folds dataset analysis we used DISOPRED3[20] (see *Novel folds* section below).

### 8. Novel folds

#### 8.1. Novel fold identification: overall procedure

Novel folds were identified in a two-step procedure: First, we compared protein models in the *MIP_curated* dataset to CATH superfamilies using the TM-score from TMalign. All resulting models without significant matches, i.e. that had a TM-score < 0.5, were then compared against the PDB90, again using the TM-score from TMalign (see below for filtering procedure and cutoff justification). All models without matches in the PDB90 and which had a high agreement score between Rosetta and DMPFold models (agreement TM-score >= 0.5) were never-seen-before structures, of which were 452. These were then clustered into 161 novel folds (see below for clustering procedure) and verified through AlphaFold2, which identified 13 false positives, resulting in 438 novel structures clustered into 148 novel folds.

#### 8.2. Novel fold identification: verifying a sensible TM-score cutoff

To choose and verify a sensible cutoff value for structural similarity using the TM-score, we computed the posterior probability that two folds within CATH 4.2.0 are similar (Fig. S36). It has been shown previously (see reference [29], Fig. 6) that a TM-score = 0.4 corresponds to at most 5% probability that two folds are similar.

Defining novel folds by a strict cut-off is debatable, but a 0.5 cut-off value is a meaningful choice given the data and is the most common definition used (see e.g. reference[30]). For 0.55 the probability of finding a known fold rises to 60-70% which still doesn't guarantee that a given fold is not novel. In these cases, a manual inspection of the models is required (see below).

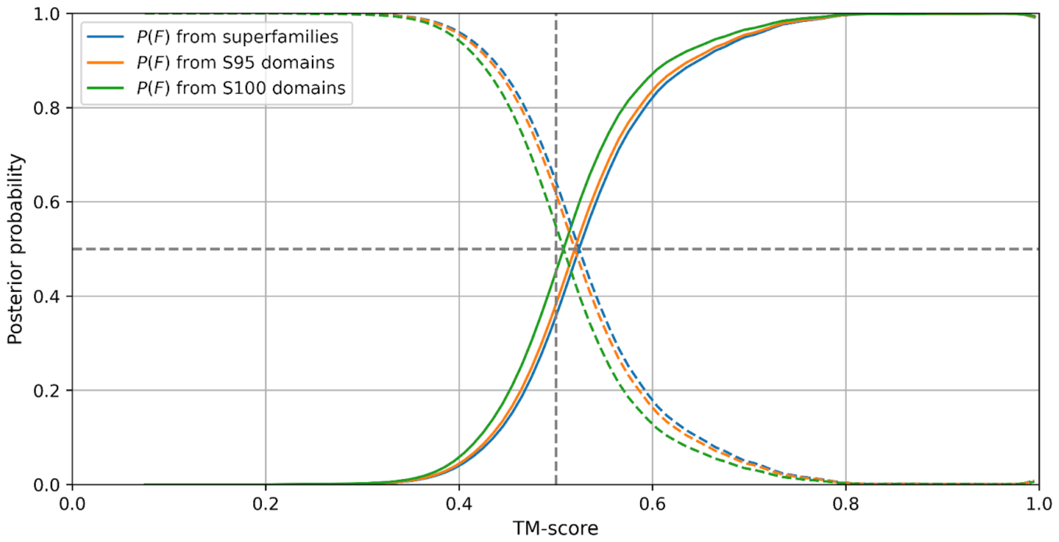

*Fig. S36: Validating the TM-score cutoff by computing the posterior probability that two proteins in CATH 4.2.0 share the same fold, verifying that a TM-score cutoff of 0.5 is a sensible choice.*

#### 8.3. Novel fold identification: comparison against CATH and PDB90

The CATH filtering step is depicted in Fig. S37 and shows maximum TM-scores for all *MIP_curated* models for both Rosetta and DMFold. Models that survived are placed in the lower left quadrant. Interestingly, the PDB filtering step is much more restrictive for Rosetta predictions - see Table S4. Coil content increases after each filtering step (see Fig. S39) but, in general, is still moderate. Such behavior is expected, since higher coil content generally translates into lower agreement with experimental (well-folded) structures deposited in CATH/PDB. The coil content is usually lower for Rosetta predictions which are constructed from fragments, in contrast to DMPFold. However, the coil content in our novel fold examples is relatively low (~30% on average).

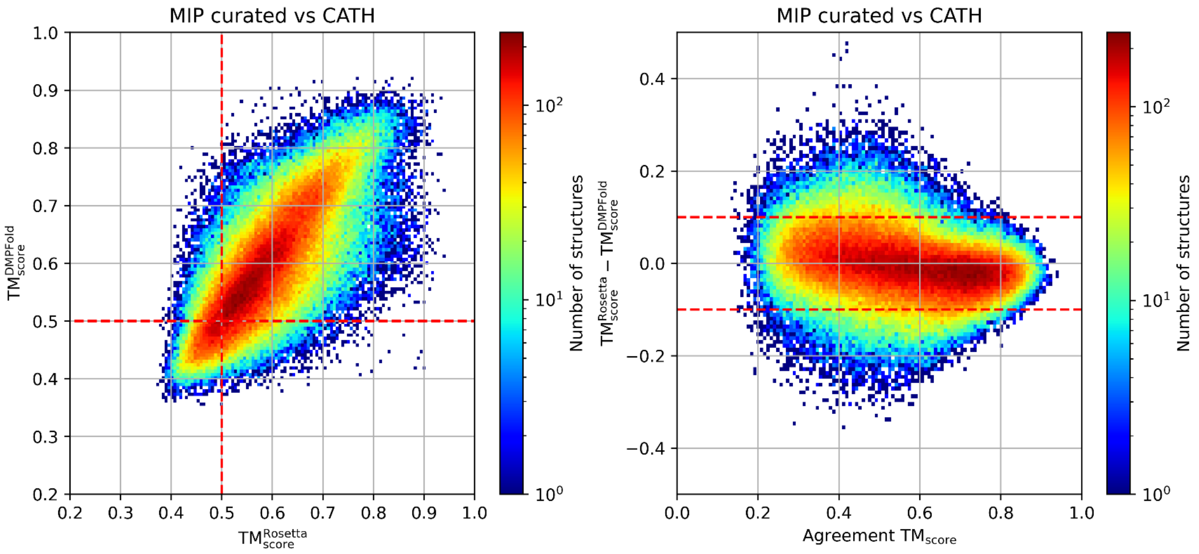

*Fig. S37: CATH filtering step. Left: number of structures in MIP_curated with a given maximum TM-score between Rosetta/DMPFold (shown on axes) and CATH. Only the models in the lower left quadrant (dotted red lines) were compared against the PDB90 in a second step. Right: agreement TM-score between Rosetta and DMPFold vs the difference between TM-score values depicted in the left panel.*

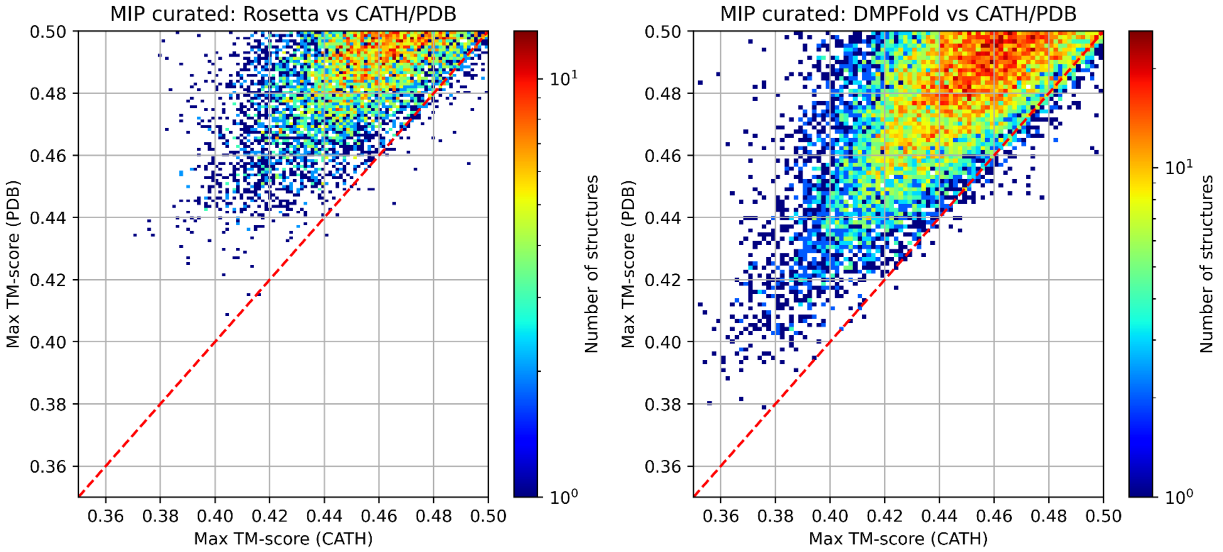

*Fig. S38: PDB vs CATH filtering step. Number of structures in MIP_curated (left: Rosetta, right: DMPFold) with a given maximum TM-score against CATH (horizontal axis) and PDB (vertical axis). Note that the TM-scores are <0.5 indicating low agreement between the predicted models and CATH and PDB.*

**Table S4: Number of *MIP_curated* models that survived after CATH and PDB filtering steps***

| | All | # of models after CATH filtering | # of models after PDB filtering |
|---|---|---|---|
| **Rosetta curated** | 211,069 | 29,624 | 7,434 |
| **DMFPold curated** | 203,877 | 40,079 | 14,294 |
| **Common** | 184,642 | 16,175 | 2,751 |
| **Common (TM-score >= 0.5)** | | | **452** |

**Note that about twice as many DMPFold models survived after both filtering steps, than Rosetta models, which likely originates in the facts that DMPFold models have a higher quality for larger proteins as compared to Rosetta and that larger proteins are less likely to have a match in CATH or the PDB due to the increased conformational space it occupies. Further, DMPFold models have a higher coil content than Rosetta models (see Fig. S4), thereby increasing the conformational space and decreasing the likelihood of a match to CATH or the PDB. The number of putative novel folds is in the last row and last column.*

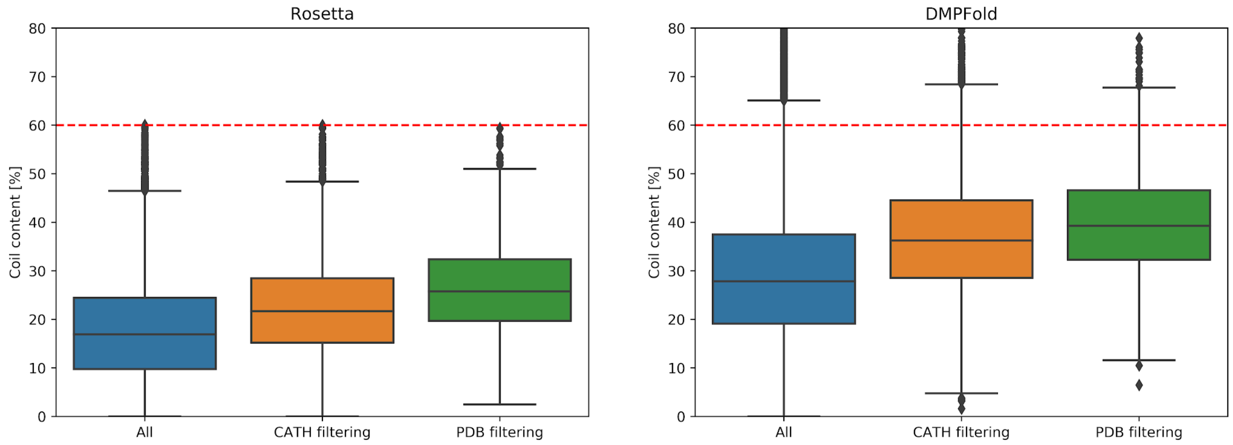

*Fig. S39: Coil content in MIP_curated after CATH and PDB filtering steps. DMPFold models generally have a higher coil content than Rosetta models.*

#### 8.4. Clustering procedure to identify novel fold clusters

We identified novel fold clusters within the set of 452 putative novel folds as follows - the Rosetta and DMPFold datasets were handled separately unless otherwise noted:

1. The 452 novel fold predictions from Rosetta (or DMPFold) were superimposed against each other to compute pairwise TM-scores, resulting in `N*(N-1)/2` comparisons. In each dataset (Rosetta or DMPFold) we found similar models with a TM-score >= 0.5 (normalization by query or target sequence length is not relevant since the comparisons are symmetric).
2. We used the TM-score comparisons as a list of edges and created graphs using the Python package NetworkX. Node positions were computed using the Fruchterman-

Reingold force-directed algorithm whereas connected components (clusters) were computed using the BFS (Breadth-first search) algorithm.

3. From the two datasets (Rosetta/DMPFold) we took the intersection between the two sets as the final set of novel folds.

Step 1 is summarized in Algorithm S1 - note that edge values i.e. node distances are computed as `1 - tmscore(model_1, model_2).`

- As a result, the 452 putative novel folds were clustered into 161 novel folds
  - 7 are large clusters (with size 10 to 87)
  - 40 are medium-size clusters (with size 2-9)
  - 114 are singletons
- After discarding 6 hard and 8 soft false positives through AlphaFold2 verification (see section below) we are left with 438 putative novel folds and 148 common clusters where
  - 7 are large clusters (with size 10 to 87)
  - 39 are medium-size clusters (with size 2-9)
  - 102 are singletons

---

**Algorithm S1: Construction of Rosetta and DMPFold edges used as input in NetworkX**

---

```
for dataset in [Rosetta, DMPFold]
    edges[dataset] = []
    # run through all models
    for model_1 in dataset
        similar = []
        # filter out similar models
        for model_2 in dataset
            if tmscore(model_1, model_2) >= 0.5
                similar.add(model_2)
            end if
        end for
        # fill in the `edges` lists
        for model_2 in similar
            distance = 1 - tmscore(model_1, model_2)
            edges[dataset].add([model_1, model_2, distance])
        end for
    end for
end for
```

---

**Table S5: MIP IDs for the novel fold clusters. Note that these include the false positives.**

| cluster # | MIP ID |
|---|---|
| 1 | MIP_00164068 |
| 2 | MIP_00296181 |
| 3 | MIP_00055264 |

| | |
|---|---|
| 4 | MIP_00222906 |
| 5 | MIP_00253881 |
| 6 | MIP_00221703 |
| 7 | MIP_00306873 |
| 8 | MIP_00154417 |
| 9 | MIP_00146970 |
| 10 | MIP_00216375 |
| 11 | MIP_00251287 |
| 12 | MIP_00230597 |
| 13 | MIP_00144099 |
| 14 | MIP_00304704 |
| 15 | MIP_00248745 |
| 16 | MIP_00258736 |
| 17 | MIP_00202629 |
| 18 | MIP_00278955 |
| 19 | MIP_00297128 |
| 20 | MIP_00081741 |
| 21 | MIP_00251063 |
| 22 | MIP_00206161 |
| 23 | MIP_00146543 |
| 24 | MIP_00241727 |
| 25 | MIP_00201366 |
| 26 | MIP_00225957 |
| 27 | MIP_00214825 |
| 28 | MIP_00191193 |
| 29 | MIP_00297536 |
| 30 | MIP_00311347 |
| 31 | MIP_00197281 |
| 32 | MIP_00280029 |
| 33 | MIP_00262260 |
| 34 | MIP_00081369 |
| 35 | MIP_00259617 |
| 36 | MIP_00240118 |
| 37 | MIP_00308388 |
| 38 | MIP_00152462 |
| 39 | MIP_00260795 |
| 40 | MIP_00225862 |
| 41 | MIP_00094786 |
| 42 | MIP_00084537 |
| 43 | MIP_00055961 |
| 44 | MIP_00309458 |
| 45 | MIP_00204672 |
| 46 | MIP_00309641 |
| 47 | MIP_00260903 |
| 48 | MIP_00161949 |
| 49 | MIP_00156786 |
| 50 | MIP_00267839 |
| 51 | MIP_00195930 |
| 52 | MIP_00198179 |
| 53 | MIP_00254475 |

| | |
|---|---|
| 54 | MIP_00153147 |
| 55 | MIP_00157706 |
| 56 | MIP_00302280 |
| 57 | MIP_00220756 |
| 58 | MIP_00054198 |
| 59 | MIP_00295365 |
| 60 | MIP_00296597 |
| 61 | MIP_00245964 |
| 62 | MIP_00196377 |
| 63 | MIP_00279819 |
| 64 | MIP_00226049 |
| 65 | MIP_00159413 |
| 66 | MIP_00254274 |
| 67 | MIP_00157006 |
| 68 | MIP_00049171 |
| 69 | MIP_00307315 |
| 70 | MIP_00217963 |
| 71 | MIP_00253334 |
| 72 | MIP_00240690 |
| 73 | MIP_00059601 |
| 74 | MIP_00303678 |
| 75 | MIP_00310815 |
| 76 | MIP_00292565 |
| 77 | MIP_00266643 |
| 78 | MIP_00180717 |
| 79 | MIP_00194449 |
| 80 | MIP_00202088 |
| 81 | MIP_00204905 |
| 82 | MIP_00055068 |
| 83 | MIP_00145352 |
| 84 | MIP_00309094 |
| 85 | MIP_00102285 |
| 86 | MIP_00294478 |
| 87 | MIP_00074380 |
| 88 | MIP_00257721 |
| 89 | MIP_00157952 |
| 90 | MIP_00271154 |
| 91 | MIP_00158688 |
| 92 | MIP_00229184 |
| 93 | MIP_00199421 |
| 94 | MIP_00250694 |
| 95 | MIP_00295500 |
| 96 | MIP_00201772 |
| 97 | MIP_00215628 |
| 98 | MIP_00241978 |
| 99 | MIP_00158052 |
| 100 | MIP_00220620 |
| 101 | MIP_00227741 |
| 102 | MIP_00189616 |
| 103 | MIP_00278709 |

| | |
|---|---|
| 104 | MIP_00213258 |
| 105 | MIP_00214517 |
| 106 | MIP_00217686 |
| 107 | MIP_00278848 |
| 108 | MIP_00230606 |
| 109 | MIP_00252634 |
| 110 | MIP_00308970 |
| 111 | MIP_00050646, MIP_00054709 |
| 112 | MIP_00185383, MIP_00307639 |
| 113 | MIP_00197470, MIP_00197989 |
| 114 | MIP_00146881, MIP_00157428 |
| 115 | MIP_00197321, MIP_00214392 |
| 116 | MIP_00147428, MIP_00155394 |
| 117 | MIP_00227385, MIP_00246175 |
| 118 | MIP_00244693, MIP_00257234 |
| 119 | MIP_00087284, MIP_00092279 |
| 120 | MIP_00268380, MIP_00326517 |
| 121 | MIP_00196533, MIP_00213475 |
| 122 | MIP_00095928, MIP_00105602 |
| 123 | MIP_00309498 |
| 124 | MIP_00296892 |
| 125 | MIP_00279120, MIP_00325018 |
| 126 | MIP_00295467, MIP_00324908 |
| 127 | MIP_00274364, MIP_00325219 |
| 128 | MIP_00193161, MIP_00229097, MIP_00233299 |
| 129 | MIP_00227792, MIP_00260396, MIP_00307513 |
| 130 | MIP_00245301, MIP_00271607, MIP_00303146 |
| 131 | MIP_00156248, MIP_00180064, MIP_00185265 |
| 132 | MIP_00052143, MIP_00057946, MIP_00063157 |
| 133 | MIP_00197711, MIP_00208158, MIP_00252474 |
| 134 | MIP_00247788, MIP_00251626, MIP_00264403 |
| 135 | MIP_00297599, MIP_00310617, MIP_00326283 |
| 136 | MIP_00226272, MIP_00240663, MIP_00298054 |
| 137 | MIP_00078622, MIP_00295276, MIP_00325113 |
| 138 | MIP_00278553, MIP_00297349, MIP_00299462, MIP_00311387 |
| 139 | MIP_00179258, MIP_00187054, MIP_00199795, MIP_00208243 |
| 140 | MIP_00178014, MIP_00213767, MIP_00230966, MIP_00245922 |
| 141 | MIP_00197039, MIP_00222838, MIP_00240930, MIP_00265151 |
| 142 | MIP_00246539, MIP_00248499, MIP_00255736, MIP_00257828, MIP_00273178 |
| 143 | MIP_00297499, MIP_00297626, MIP_00299300, MIP_00306653, MIP_00308130 |
| 144 | MIP_00068041, MIP_00085074, MIP_00085909, MIP_00144174, MIP_00159046, MIP_00180365 |
| 145 | MIP_00260750, MIP_00271812, MIP_00280561, MIP_00294938, MIP_00295429, MIP_00308577 |
| 146 | MIP_00247823, MIP_00249069, MIP_00278640, MIP_00295849, MIP_00302861, MIP_00325914 |
| 147 | MIP_00176786, MIP_00199585, MIP_00206875, MIP_00212347, MIP_00221839, MIP_00231479, MIP_00232949 |
| 148 | MIP_00217796, MIP_00217874, MIP_00225385, MIP_00232829, MIP_00240077, MIP_00258985, MIP_00269643, MIP_00299431 |
| 149 | MIP_00260429, MIP_00296222, MIP_00299925, MIP_00304110, MIP_00305968, MIP_00307861, MIP_00309352, MIP_00326591 |
| 150 | MIP_00254040 |

| | |
|---|---|
| 151 | MIP_00147377 |
| 152 | MIP_00249344, MIP_00261138, MIP_00262362, MIP_00264146, MIP_00304501, MIP_00326528 |
| 153 | MIP_00210273, MIP_00212106, MIP_00222133, MIP_00223995, MIP_00248863, MIP_00251996, MIP_00263890, MIP_00273556, MIP_00299160 |
| 154 | MIP_00088406, MIP_00165525, MIP_00211356, MIP_00212714, MIP_00214515, MIP_00215955, MIP_00224102, MIP_00232465, MIP_00241193 |
| 155 | MIP_00091051, MIP_00154142, MIP_00158467, MIP_00183571, MIP_00185281, MIP_00220809, MIP_00224343, MIP_00228269, MIP_00232818, MIP_00234843, MIP_00254538 |
| 156 | MIP_00166287, MIP_00177071, MIP_00178352, MIP_00182849, MIP_00183599, MIP_00184460, MIP_00196992, MIP_00198685, MIP_00199027, MIP_00234745, MIP_00240645, MIP_00296972 |
| 157 | MIP_00193894, MIP_00207383, MIP_00209661, MIP_00211255, MIP_00212034, MIP_00216727, MIP_00217454, MIP_00241860, MIP_00253458, MIP_00254609, MIP_00259051, MIP_00272986 |
| 158 | MIP_00212877, MIP_00226974, MIP_00240585, MIP_00241642, MIP_00244666, MIP_00246167, MIP_00248174, MIP_00253352, MIP_00258163, MIP_00258887, MIP_00259789, MIP_00261729, MIP_00265735, MIP_00266464, MIP_00267641, MIP_00279236, MIP_00292101, MIP_00294595, MIP_00324660 |
| 159 | MIP_00154164, MIP_00154467, MIP_00154943, MIP_00157617, MIP_00157981, MIP_00158076, MIP_00158664, MIP_00160017, MIP_00162356, MIP_00163233, MIP_00164924, MIP_00165656, MIP_00176391, MIP_00177659, MIP_00182860, MIP_00183204, MIP_00185488, MIP_00185557, MIP_00190387, MIP_00190667, MIP_00191346, MIP_00198257, MIP_00216029 |
| 160 | MIP_00261424, MIP_00263752, MIP_00264541, MIP_00264596, MIP_00267406, MIP_00269355, MIP_00274524, MIP_00278918, MIP_00290826, MIP_00292559, MIP_00292648, MIP_00298985, MIP_00299754, MIP_00302700, MIP_00303803, MIP_00305120, MIP_00306932, MIP_00307360, MIP_00308485, MIP_00309593, MIP_00310169, MIP_00324816, MIP_00325274 |
| 161 | MIP_00206574, MIP_00210880, MIP_00212896, MIP_00220471, MIP_00221200, MIP_00221345, MIP_00221370, MIP_00221394, MIP_00223115, MIP_00223708, MIP_00228994, MIP_00229177, MIP_00230672, MIP_00233022, MIP_00234467, MIP_00234709, MIP_00242090, MIP_00243548, MIP_00243551, MIP_00244111, MIP_00244990, MIP_00246103, MIP_00247485, MIP_00247996, MIP_00248322, MIP_00249061, MIP_00249438, MIP_00249931, MIP_00250573, MIP_00251953, MIP_00252387, MIP_00253275, MIP_00254698, MIP_00254824, MIP_00259384, MIP_00259718, MIP_00259989, MIP_00260973, MIP_00261827, MIP_00261864, MIP_00262086, MIP_00262092, MIP_00263644, MIP_00266479, MIP_00266497, MIP_00267415, MIP_00268211, MIP_00268218, MIP_00268903, MIP_00269480, MIP_00270210, MIP_00270267, MIP_00273381, MIP_00274428, MIP_00278570, MIP_00278861, MIP_00279211, MIP_00279647, MIP_00292796, MIP_00293153, MIP_00293157, MIP_00294176, MIP_00294930, MIP_00294968, MIP_00296715, MIP_00297556, MIP_00298218, MIP_00298310, MIP_00300516, MIP_00302964, MIP_00305558, MIP_00305639, MIP_00306385, MIP_00307596, MIP_00308279, MIP_00308625, MIP_00309604, MIP_00310801, MIP_00313352, MIP_00313433, MIP_00324442, MIP_00324963, MIP_00325368, MIP_00325983, MIP_00326003, MIP_00326623, MIP_00326689 |

#### 8.5. AlphaFold2 verification of putative novel folds

We ran AlphaFold2 on all of 452 putative novel fold sequences. Fig. S40 compares Rosetta, DMPFold and AlphaFold2 predictions. The majority of our predictions agree well with AlphaFold2. Interestingly, the mean agreement TM-scores between Rosetta/AlphaFold2 and DMPFold/AlphaFold2 are higher than the one between Rosetta and DMPFold which may indicate that AlphaFold2 predictions possess some features of both Rosetta and DMPFold models.

Moreover, the agreement TM-score between Rosetta/AlphaFold2 and DMPFold/AlphaFold2 models correlates well with Rosetta/DMPFold MQA scores (see Fig. S42). We also see a slightly weaker correlation between the mean pLDDT score (i.e. AlphaFold2 MQA score) and Rosetta/DMPFold MQA scores (see Fig. S43).

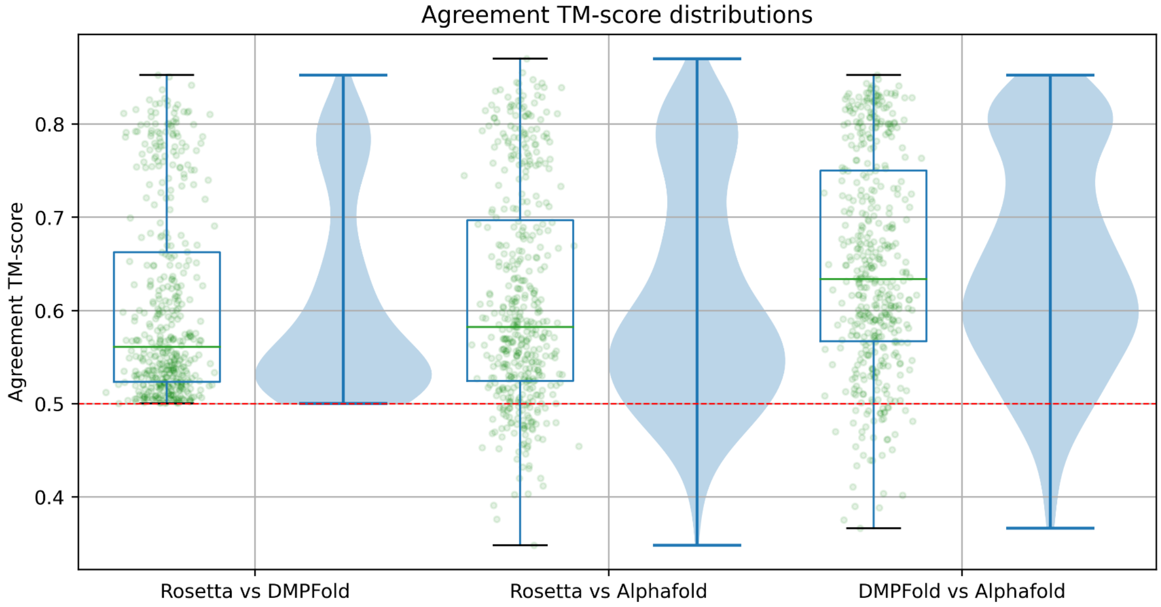

*Fig.*
*Fig. S40: Comparison between Rosetta, DMPFold and AlphaFold2 putative novel fold predictions. Points correspond to agreement TM-score between two given methods. Note the scale on the y-axis doesn't cover the full range of TM-scores from 0 to 1.*

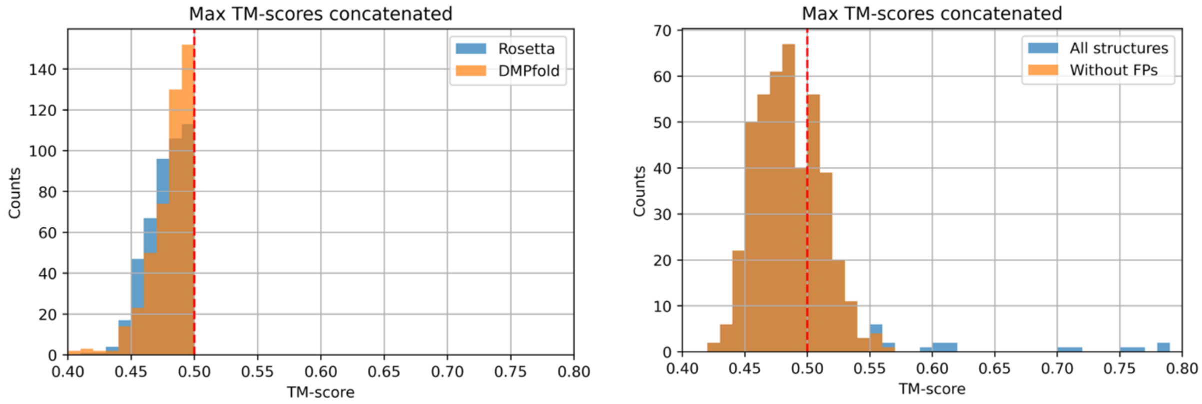

*Fig. S41: Distribution of maximum TM-scores against PDB from putative novel folds dataset (see Table S5) for Rosetta and DMPFold (left) and AlphaFold predictions (right).*

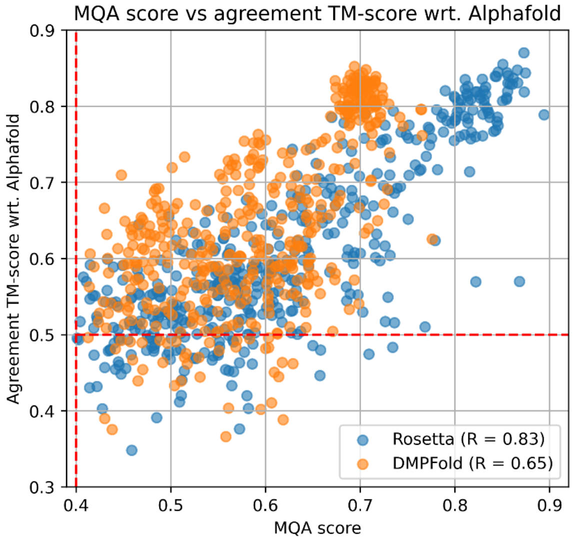

*Fig. S42: MQA score vs agreement TM-score between Rosetta/DMPFold and AlphaFold2. All structures come from the putative novel fold dataset. Pearson R correlation coefficients are shown in the legends.*

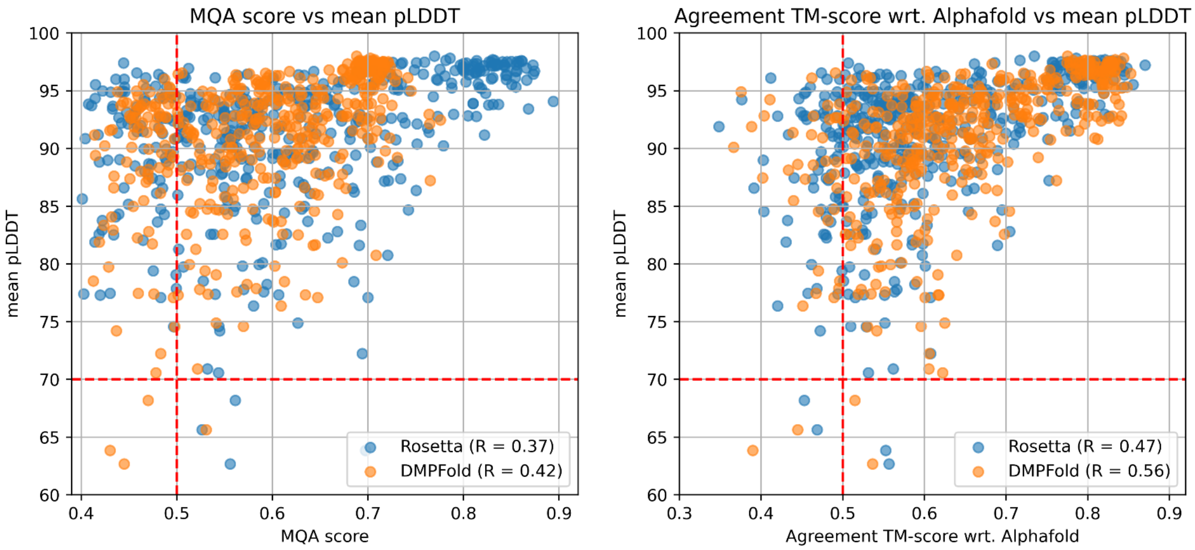

*Fig. S43: Left: MQA score for Rosetta and DMPFold vs mean pLDDT (local accuracy of AlphaFold2 predictions). Right: agreement TM-score between Rosetta/AlphaFold2 and DMPFold/AlphaFold2 vs mean pLDDT. All models come from the putative novel fold dataset. Pearson R correlation coefficients are shown in the legends.*

#### *Potential false positives*

For the majority of AlphaFold2 novel fold models the maximum TM-score against CATH and PDB is below the threshold of 0.5 (see Fig. S44). However, there are some models for which similar structures are already deposited in the PDB. We decided not to discard models just above the threshold (~0.5-0.55) since such predictions should be verified experimentally. For the remainder, we distinguished two types of potential false positives (see Fig. S44):

- Hard false positives - total of 6 models:

    ○ Maximum TM-score between AlphaFold2 model and PDB >= 0.7
- Soft false positives (SFP) - total of 8 models:
    ○ maximum TM-score between AlphaFold2 model and PDB >= 0.58 (slightly below 0.6 - see Fig. S44)
    ○ **or** maximum TM-score between AlphaFold2 model and CATH >= 0.55
    ○ **minus** Hard False Positive above

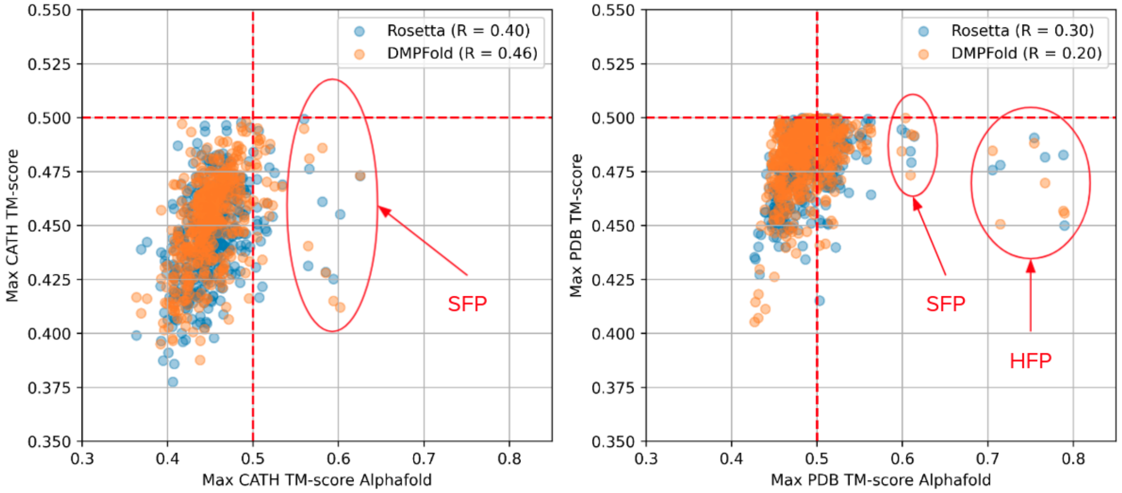

*Fig. S44: Left: maximum TM-score between AlphaFold2 and CATH (horizontal axis) and between Rosetta/DMPFold and CATH (vertical axis). Right: maximum TM-score between AlphaFold2 and PDB (horizontal axis) and between Rosetta/DMPFold and PDB (vertical axis). All structures come from the putative novel fold dataset. Pearson R correlation coefficients are shown in the legends.*

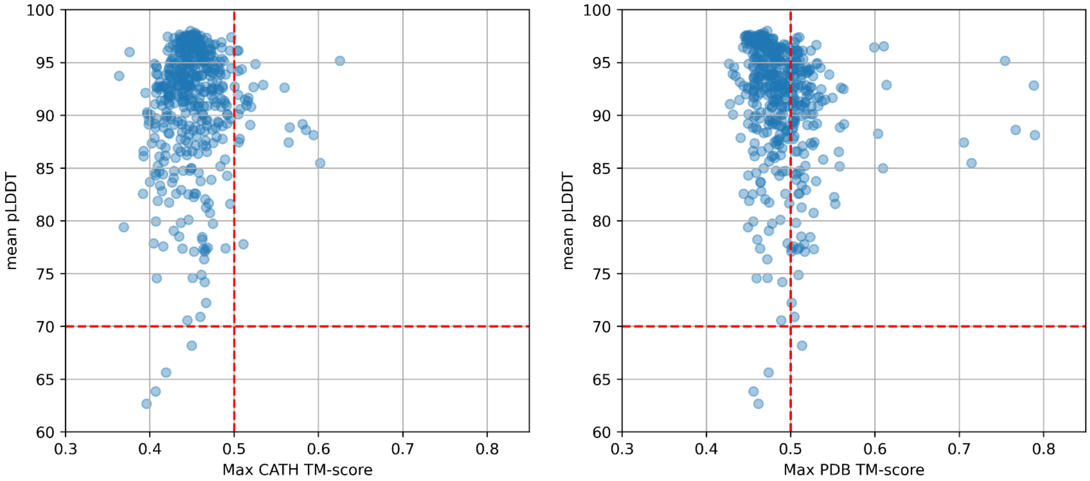

*Fig. S45: maximum TM-score between AlphaFold2 and CATH (left)/PDB (right) vs mean pLDDT.*

#### Table S6: Hard false positives

| | Agreement TM-score vs Rosetta | Agreement TM-score vs DMPFold | CATH homologous superfamily number | CATH TM-score (MIP-normed) | PDB90 domain name | PDB90 TM-score (MIP-normed) | DISOPRED 3 percentage | mean pLDDT | Cluster ID |
|---|---|---|---|---|---|---|---|---|---|

| | | | | | | | | | |
|---|---|---|---|---|---|---|---|---|---|
| MIP_00152462 | 0.525 | 0.440 | 2x8kC01 | 0.602 | 5w5fA | 0.714 | 3.4 | 85.5 | 38 |
| MIP_00201772 | 0.470 | 0.433 | 4hkhA00 | 0.594 | 6j0nn | 0.789 | 0.0 | 88.1 | 96 |
| MIP_00074380 | 0.454 | 0.404 | 3vkiA02 | 0.425 | 3teeA | 0.788 | 7.6 | 92.8 | 87 |
| MIP_00220620 | 0.528 | 0.445 | 2x8kC01 | 0.585 | 6j0nb | 0.767 | 0.0 | 88.6 | 100 |
| MIP_00297536 | 0.480 | 0.483 | 4hkhA00 | 0.565 | 6xgrA | 0.705 | 4.2 | 87.4 | 29 |
| MIP_00196377 | 0.517 | 0.537 | 2qzbA00 | 0.625 | 4ifaA | 0.754 | 0.0 | 95.2 | 62 |

**Table S7: Soft false positives**

| | Agreement TM-score vs Rosetta | Agreement TM-score vs DMPFold | CATH homologous superfamily number | CATH TM-score (MIP-normed) | PDB90 domain name | PDB90 TM-score (MIP-normed) | DISOPRED 3 percentage | mean pLDDT | Cluster ID |
|---|---|---|---|---|---|---|---|---|---|
| MIP_00280029 | 0.554 | 0.585 | 3h0nA00 | 0.560 | 3h0nA | 0.560 | 0.7 | 92.6 | 32 |
| MIP_00227385 | 0.501 | 0.532 | 3ar4A02 | 0.449 | 6a2uA | 0.609 | 18.6 | 85.0 | 117 |
| MIP_00253881 | 0.558 | 0.586 | 4fczA00 | 0.566 | 5uwbB | 0.558 | 10.1 | 88.9 | 5 |
| MIP_00266643 | 0.551 | 0.690 | 2z0rC00 | 0.581 | 2z0rA | 0.563 | 0.0 | 89.2 | 77 |
| MIP_00251287 | 0.545 | 0.645 | 5gzkA00 | 0.423 | 6y56A | 0.599 | 0.0 | 96.5 | 11 |
| MIP_00309094 | 0.467 | 0.502 | 5o6hA00 | 0.534 | 5kf2A | 0.614 | 1.8 | 92.9 | 84 |
| MIP_00054198 | 0.474 | 0.491 | 1c5eA00 | 0.462 | 5wk1X | 0.610 | 0.0 | 96.5 | 58 |
| MIP_00246175 | 0.521 | 0.596 | 5b1aA00 | 0.423 | 6a2uA | 0.603 | 17.4 | 88.3 | 117 |

All hard and soft false positives are singletons in our novel fold clusters. Two SFPs (MIP_00227385, MIP_00246175) belong to the same cluster (cluster 117). The mean pLDDT for those AlphaFold2 predictions is generally >= 0.85, so the confidence in these predictions is pretty high (see Fig. S45). Therefore, we are more convinced that they are indeed potential false positives and we decided to remove them from the final putative novel fold dataset.

##### *A note on fragments, templates and what the prediction methods "know"*

Rosetta does not directly use templates but it uses fragments for model building. The fragments come from the *VALL* database which was last updated around 2011. Contacts for Rosetta are predicted by GREMLIN which uses multiple sequence alignments generated from public databases, which should have been trained around 2016/7.

DMPFold uses neural networks trained on public databases before 2019, so it would not have seen anything after that time.

AlphaFold2, on the other hand, downloads the entire PDB during installation or setup, to use as templates. We downloaded the PDB on July 31, 2021, so AlphaFold2 would have seen every one of the hard false positive example structures in the PDB.

##### *Hard false positive examples*

Below we show and briefly describe all hard false positive examples.

###### Hard False Positive #1

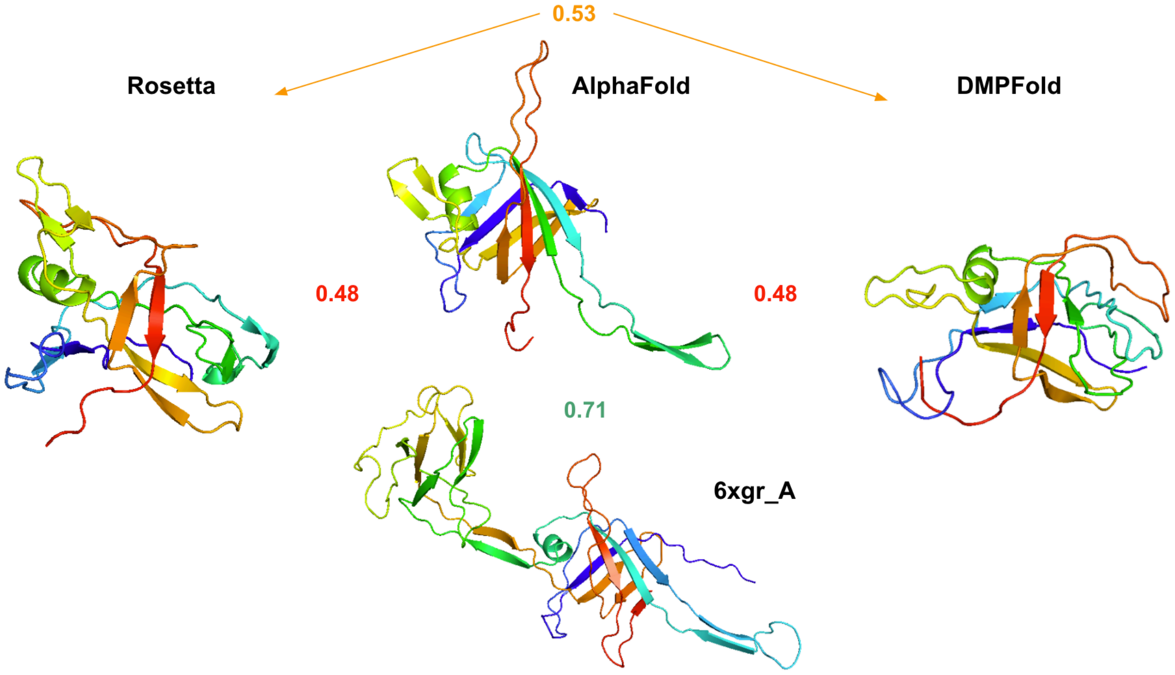

| | | | |
|---|---|---|---|
| **Max CATH TM-score:** | 0.43 | 0.56 | 0.44 |
| **Max PDB TM-score:** | 0.48 | 0.71 | 0.48 |
| **MQA score / Coil content:** | 0.47 / 28% | 0.87 / 26% | 0.6 / 27% |

*Fig. S46: MIP_00297536 structure comparison between Rosetta, AlphaFold2 and DMPFold (upper models) and the most similar PDB to the AlphaFold2 prediction (lower structure). Mutual TM-scores are shown in between. Additional parameters are listed in the table.*

- Most similar PDB90 structure for all models is 6xgr_A (**Deposited:** 2020-06-17 **Released:** 2020-07-01)
- very high agreement of PDB structure against AlphaFold2 model is due to the tail position (similar tail might be found in two templates used by AlphaFold2 for model prediction: 6nbx_N Deposited: 2018-12-10 Released: 2019-02-27; 6nby_N Deposited: 2018-12-10 Released: 2019-02-27)
- Tail fragment predicted by AlphaFold2 with mean pLDDT around 0.77 (confident)
- pTM (global score) for AlphaFold2: 0.77 (confident)

#### Hard False Positive #2

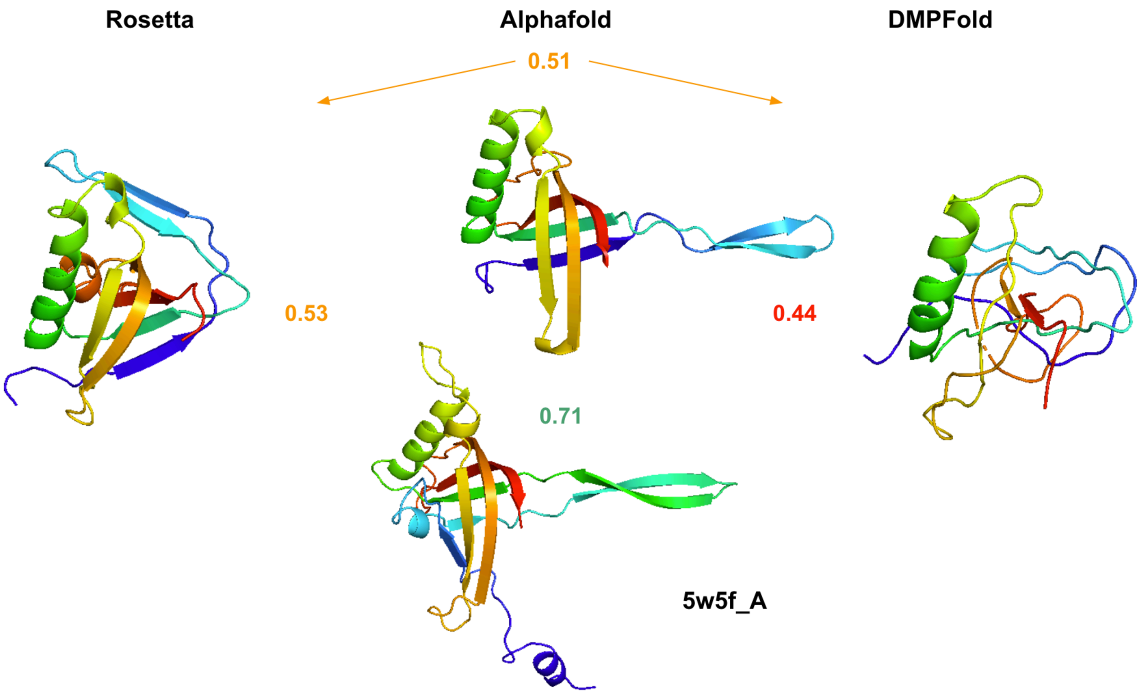

| | Rosetta | Alphafold | DMPFold |
|---|---|---|---|
| **Max CATH TM-score:** | 0.46 | 0.6 | 0.41 |
| **Max PDB TM-score:** | 0.48 | 0.71 | 0.45 |
| **MQA score / Coil content:** | 0.55 / 25% | 0.85 / 20% | 0.47 / 32% |

*Fig. S47: MIP_00152462 structure comparison between Rosetta, AlphaFold2 and DMPFold (upper models) and the most similar PDB to the AlphaFold2 prediction (lower structure). Mutual TM-scores are shown in between. Additional parameters are listed in the table.*

- All AlphaFold2 templates released in PDB database before 2019
- Most similar PDB structure for AlphaFold2 model: 5w5f_A (**Deposited:** 2017-06-15 **Released:** 2017-08-16)
- very good agreement between Rosetta and AlphaFold2 predictions except for the tail fragment
- tail fragment predicted by AlphaFold2 with mean pLDDT around 0.82 (confident)
- pTM (global score) for AlphaFold2: 0.76

#### Hard False Positive #3

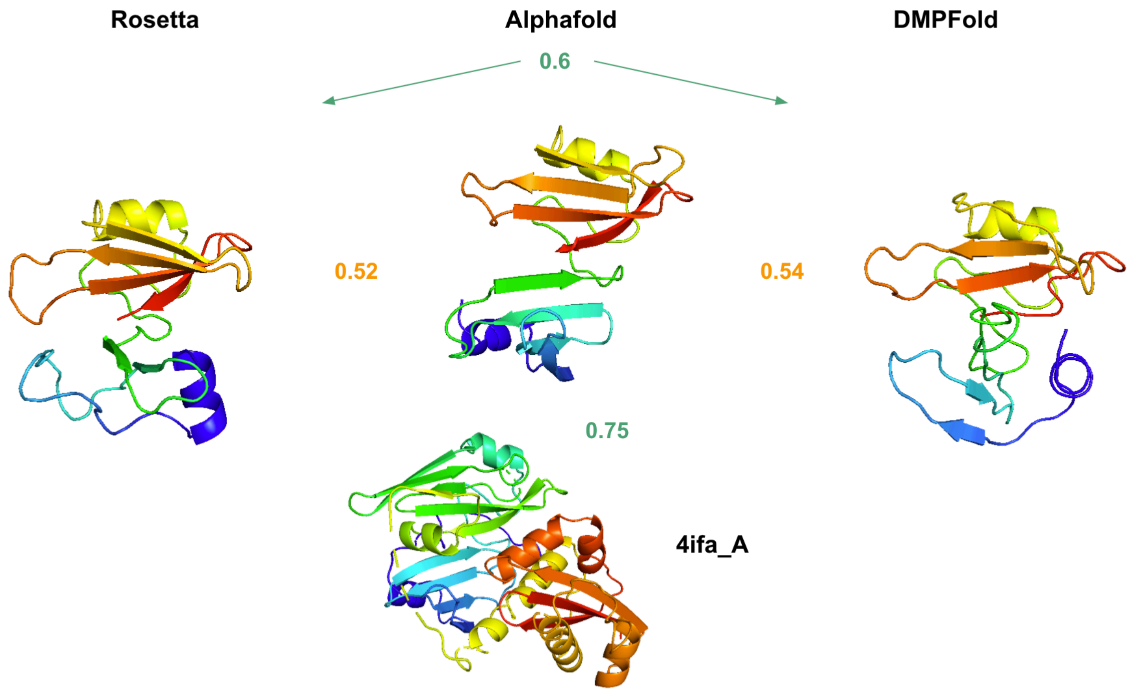

| | Rosetta | Alphafold | DMPFold |
|---|---|---|---|
| **Max CATH TM-score:** | 0.47 | 0.63 | 0.47 |
| **Max PDB TM-score:** | 0.49 | 0.75 | 0.49 |
| **MQA score / Coil content:** | 0.62 / 32% | 0.95 / 34% | 0.49 / 44% |

*Fig. S48: MIP_00196377 structure comparison between Rosetta, AlphaFold2 and DMPFold (upper models) and the most similar PDB to the AlphaFold2 prediction (lower structure). Mutual TM-scores are shown in between. Additional parameters are listed in the table.*

- All AlphaFold2 templates released in PDB database before 2019
- Most similar PDB structure for AlphaFold2 model: 4ifa_A (**Deposited:** 2012-12-14 **Released:** 2012-12-26)
- AlphaFold2 prediction has high symmetry which is the reason for high agreement to 4ifa_A
- No disorder, very confident AlphaFold2 predictions
- pTM (global score) for AlphaFold2: 0.86

#### Hard False Positive #4

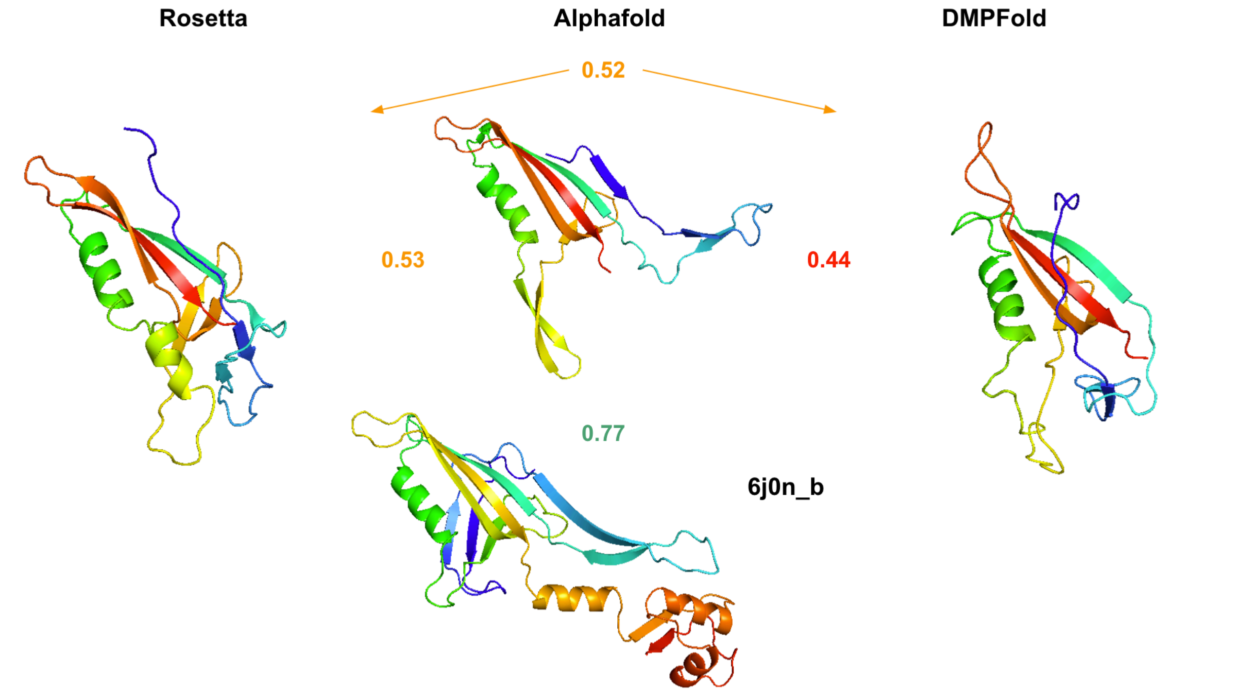

| | Rosetta | Alphafold | DMPFold |
|---|---|---|---|
| **Max CATH TM-score:** | 0.43 | 0.58 | 0.43 |
| **Max PDB TM-score:** | 0.48 | 0.77 | 0.47 |
| **MQA score / Coil content:** | 0.47 / 23% | 0.89 / 19% | 0.61 / 35% |

*Fig. S49: MIP_00220620 structure comparison between Rosetta, AlphaFold2 and DMPFold (upper models) and the most similar PDB to the AlphaFold2 prediction (lower structure). Mutual TM-scores are shown in between. Additional parameters are listed in the table.*

- Most similar PDB structure for AlphaFold2 model: 6j0n_b (**Deposited:** 2018-12-25 **Released:** 2019-04-10 )
- However, among templates used by AlphaFold2 we can find: 6rao_G Deposited: 2019-04-06 Released: 2019-04-17; 6j0n_d Deposited: 2018-12-25 Released: 2019-04-10
- Similarly to *Hard False Positive #1* and *Hard False Positive #2*, high agreement with PDB is due to matching tail fragments
- DISOPRED (sequence-based) predicts that blue/yellow tail fragments weakly disordered up to 40%/20% (17%/13% on average)
- However, they are predicted by AlphaFold2 with mean pLDDT around 0.77 (confident)
- pTM (global score) for AlphaFold2: 0.76

#### Hard False Positive #5

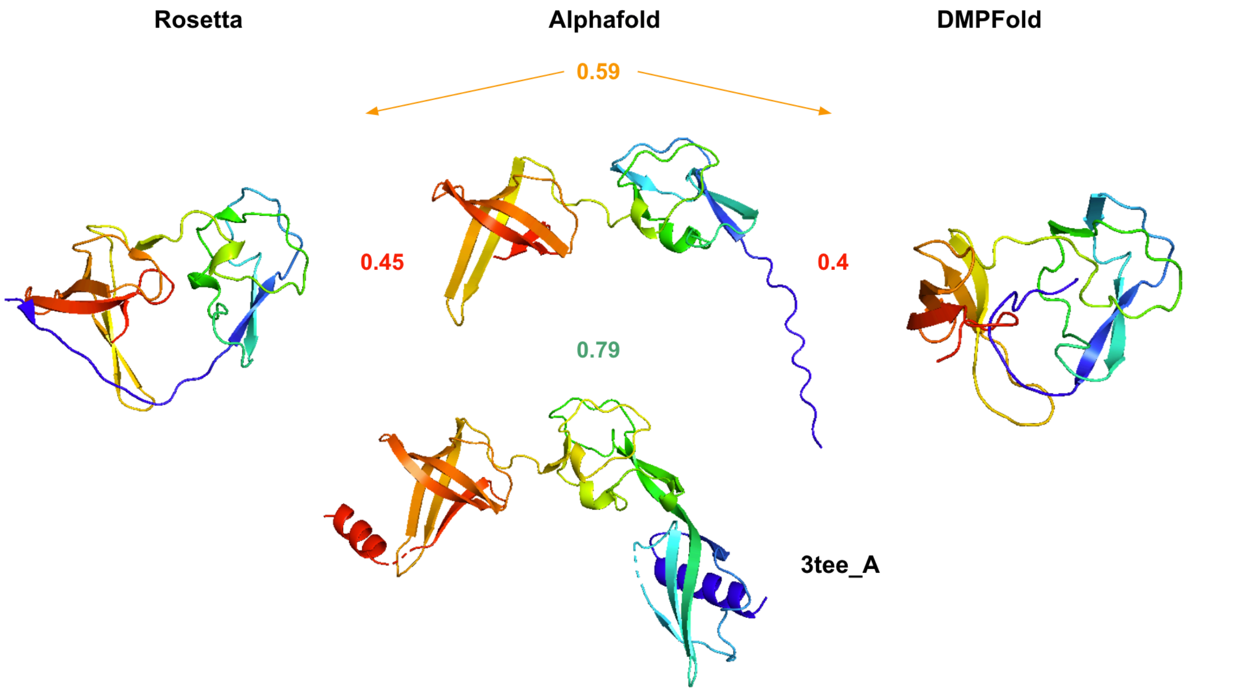

| | Rosetta | Alphafold | DMPFold |
|---|---|---|---|
| **Max CATH TM-score:** | 0.41 | 0.43 | 0.42 |
| **Max PDB TM-score:** | 0.48 | **0.79** | 0.46 |
| **MQA score / Coil content:** | 0.56 / 32% | 0.93 / 39% | 0.56 / 32% |

*Fig. S50: MIP_00074380 structure comparison between Rosetta, AlphaFold2 and DMPFold (upper models) and the most similar PDB to the AlphaFold2 prediction (lower structure). Mutual TM-scores are shown in between. Additional parameters are listed in the table.*

- the most similar PDB structure (3tee_A) was also used by AlphaFold2 as a template:
  - 3tee_A - Deposited: 2011-08-12 Released: 2012-08-15
- However, the alignment is not perfect due to disordered N-terminus (65% on average) which doesn't match well with the template (pLDDT for that fragment is around 0.63)
- pTM (global score) for AlphaFold2: 0.71

#### Hard False Positive #6

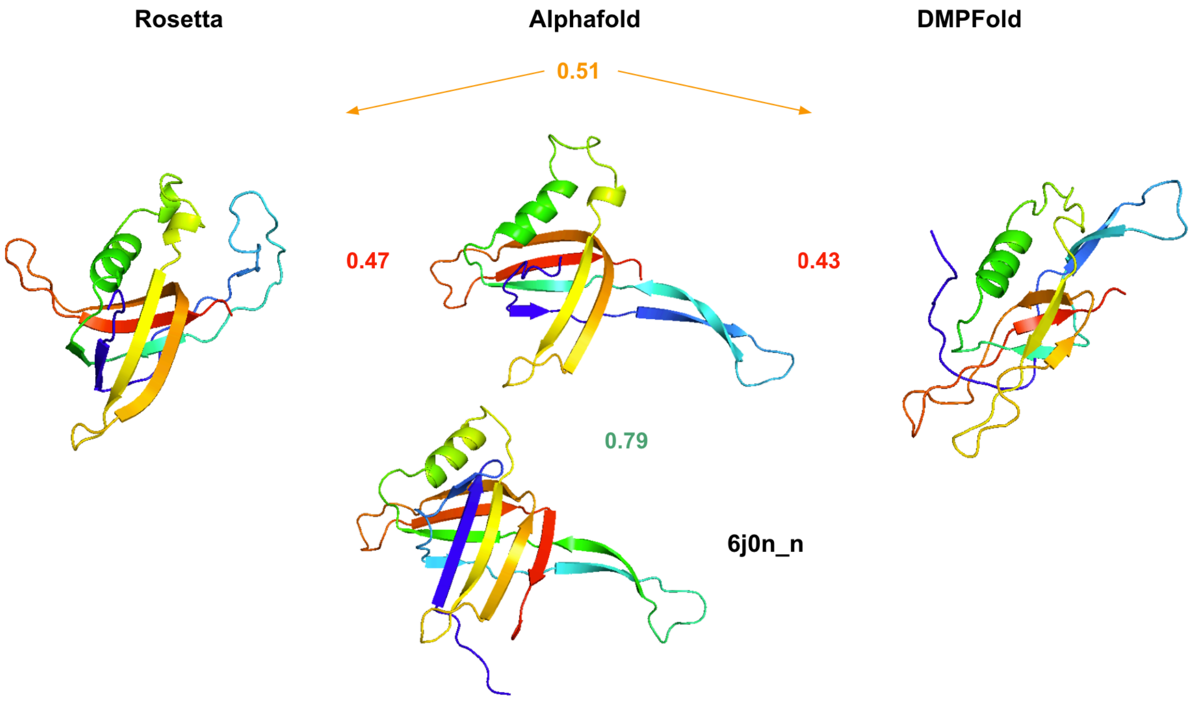

| | | | |
|---|---|---|---|
| **Max CATH TM-score:** | 0.43 | 0.59 | 0.42 |
| **Max PDB TM-score:** | 0.45 | 0.79 | 0.46 |
| **MQA score / Coil content:** | 0.52 / 26% | 0.88 / 22% | 0.5 / 26% |

*Fig. S51: MIP_00201772 structure comparison between Rosetta, AlphaFold2 and DMPFold (upper models) and the most similar PDB to the AlphaFold2 prediction (lower structure). Mutual TM-scores are shown in between. Additional parameters are listed in the table.*

- Most similar PDB structure for AlphaFold2: 6j0n_n (**Deposited:** 2018-12-25 **Released:** 2019-04-10)
- this structure (different chain name) is also present among templates used by AlphaFold2 (not all templates listed below):
  - 6rao_F Deposited: 2019-04-06 Released: 2019-04-17
  - 6j0n_r Deposited: 2018-12-25 Released: 2019-04-10
  - 6rbn_B Deposited: 2019-04-11 Released: 2019-04-24
  - 6rap_A Deposited: 2019-04-07 Released: 2019-04-17
- The prediction of the tail fragment (mean pLDDT around 94%) agrees perfectly with the template; there are however some subtle differences in the middle part (yellow color)
- No disordered fragments
- pTM (global score) for AlphaFold2: 0.78

A similar trend is observed for Soft False Positives (SFPs) i.e. AlphaFold2 predictions are generally more similar to structures in the PDB, likely because it used them as templates. However, the TM-scores between all three predictions is higher (with mean around 0.55 as compared to 0.5 for Hard False Positives). Moreover, mean pLDDT and pTM for AlphaFold2 predictions are larger: 0.8 and 0.7 respectively.

#### 8.6. Novel fold examples

Below we describe a few examples of our predicted novel folds verified by AlphaFold2.

##### *Novel fold example #1*

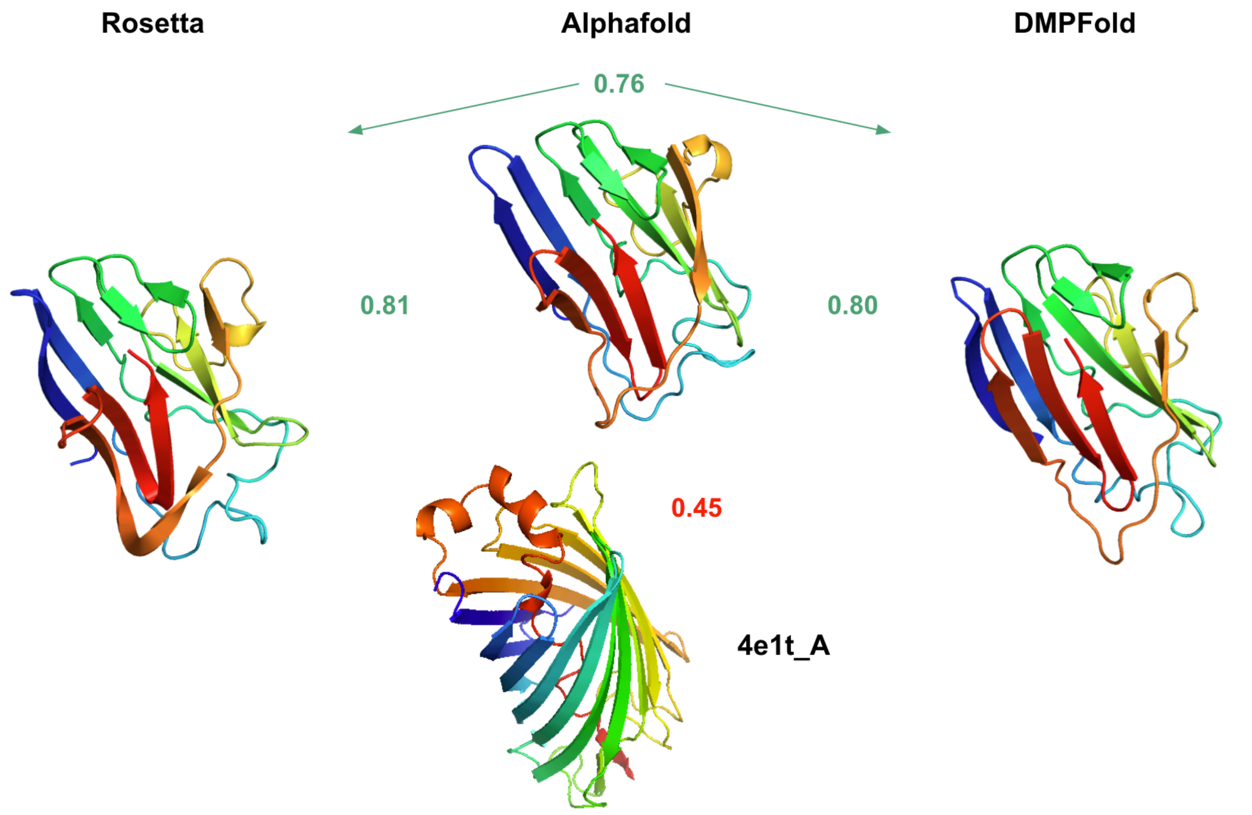

| | | | |
|---|---|---|---|
| **Max CATH TM-score:** | 0.44 | 0.44 | 0.45 |
| **Max PDB TM-score:** | 0.46 | 0.45 | 0.49 |
| **MQA score / Coil content:** | 0.80 / 30% | 0.97 / 24% | 0.71 / 31% |

*Fig. S52: MIP_00261827 (cluster 161) structure comparison between Rosetta, AlphaFold2 and DMPFold (upper models) and the most similar PDB to the AlphaFold2 prediction (lower structure). Mutual TM-scores are shown in between. Additional parameters are listed in the table.*

- this example shows the models in the largest cluster with 87 representatives
- Excellent agreement between all methods
- All max TM-scores against CATH and PDB (including AlphaFold2) below 0.5
- No disorder
- pTM (global score) for AlphaFold2: 0.90

#### *Novel fold example #2*

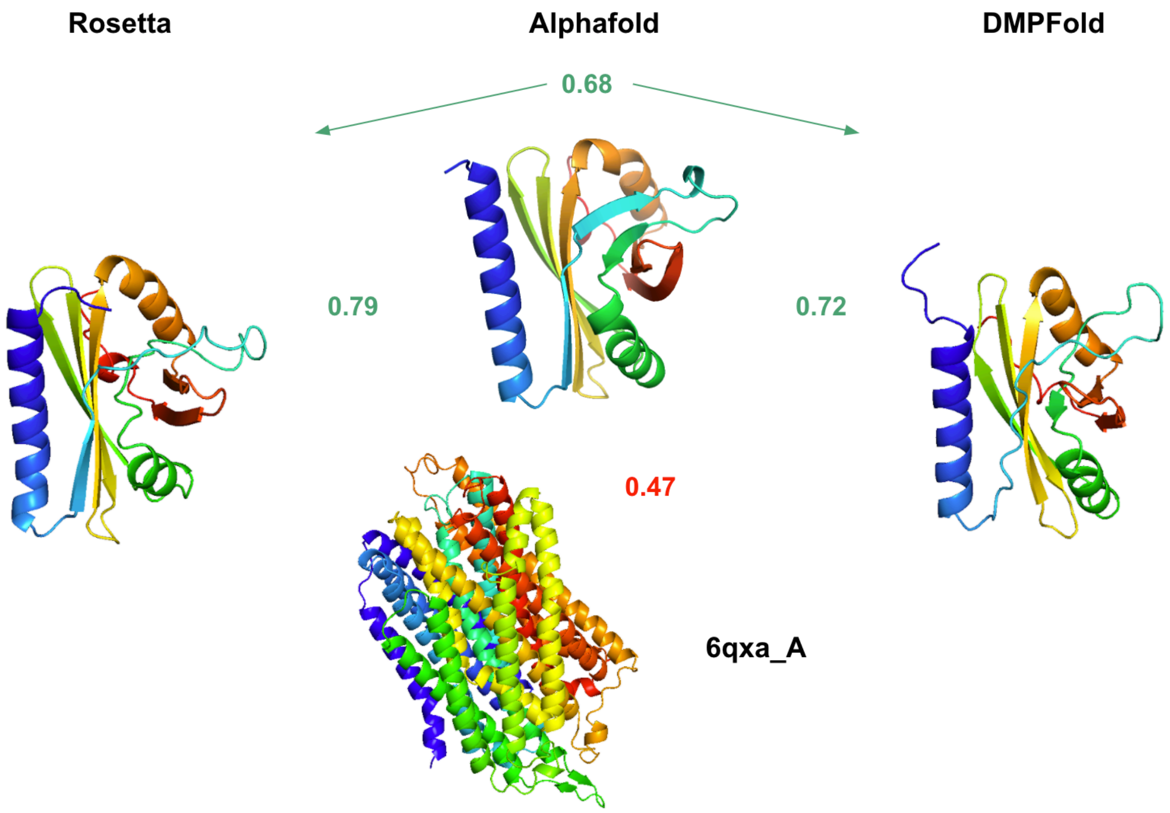

| | | | |
|---|---|---|---|
| **Max CATH TM-score:** | 0.44 | 0.43 | 0.42 |
| **Max PDB TM-score:** | 0.46 | 0.47 | 0.47 |
| **MQA score / Coil content:** | 0.67 / 19% | 0.93 / 14% | 0.68 / 19% |

*Fig. S53: MIP_00189616 (cluster 102 singleton) structure comparison between Rosetta, AlphaFold2 and DMPFold (upper models) and the most similar PDB to the AlphaFold2 prediction (lower structure). Mutual TM-scores are shown in between. Additional parameters are listed in the table.*

- singleton cluster with high agreement between predicted models and low agreement to PDB structure
- pTM (global score) for AlphaFold2: 0.88

#### *Novel fold example #3*

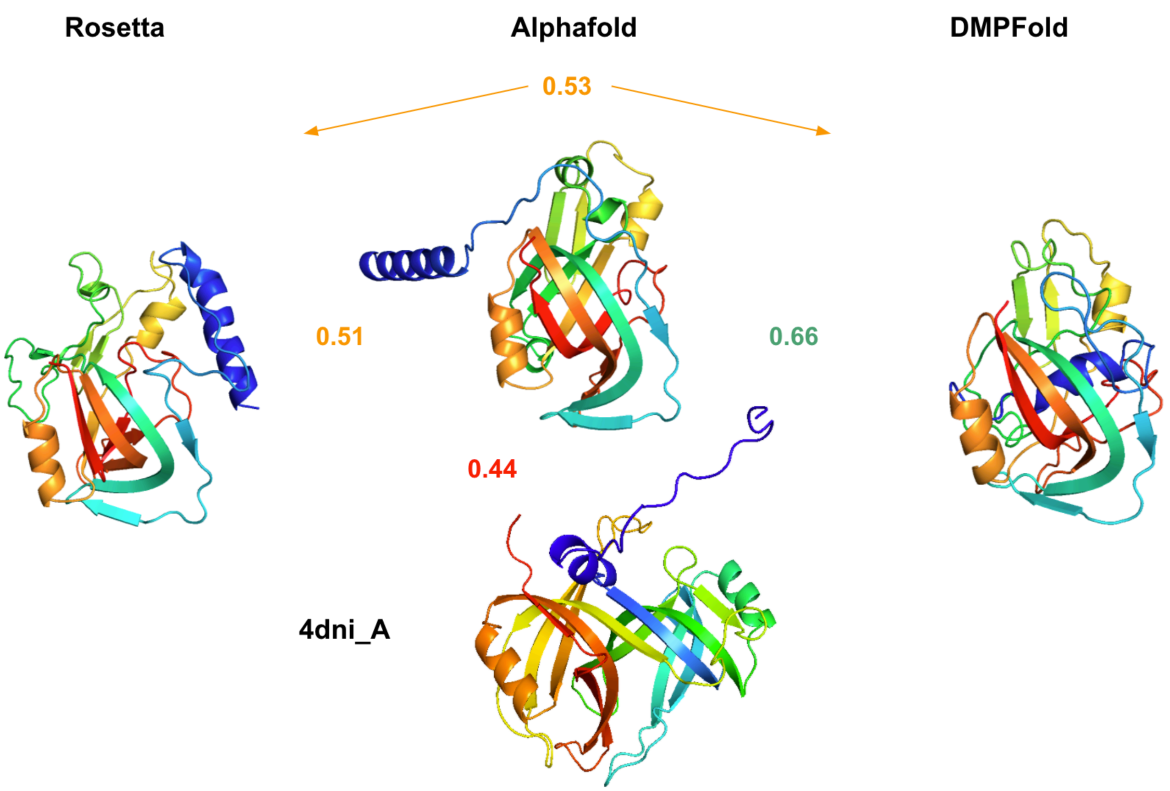

| | Rosetta | Alphafold | DMPFold |
|---|---|---|---|
| **Max CATH TM-score:** | 0.42 | 0.41 | 0.41 |
| **Max PDB TM-score:** | 0.46 | 0.44 | 0.41 |
| **MQA score / Coil content:** | 0.46 / 30% | 0.93 / 18% | 0.57 / 36% |

*Fig. S54: MIP_00307861 (cluster 149 representative) structure comparison between Rosetta, AlphaFold2 and DMPFold (upper models) and the most similar PDB to the AlphaFold2 prediction (lower structure). Mutual TM-scores are shown in between. Additional parameters are listed in the table.*

- Lower agreement due to disordered (83% on average) N-terminus alpha helix which is predicted differently by all methods
- Mean pLDDT for AlphaFold2 for that fragment is equal 72% (confident)
- pTM (global score) for AlphaFold2: 0.85

#### *Novel fold example #4*

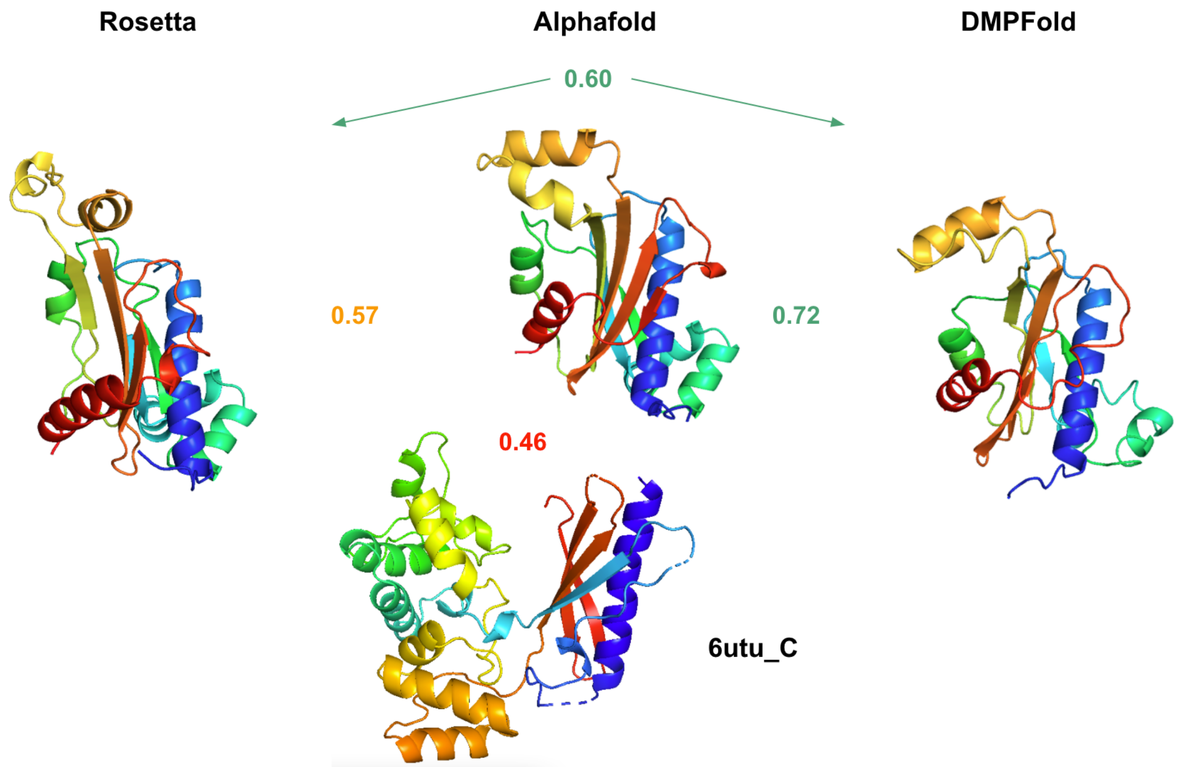

| | Rosetta | Alphafold | DMPFold |
|---|---|---|---|
| **Max CATH TM-score:** | 0.43 | 0.42 | 0.42 |
| **Max PDB TM-score:** | 0.46 | 0.46 | 0.47 |
| **MQA score / Coil content:** | 0.57 / 30% | 0.93 / 22% | 0.66 / 37% |

*Fig. S55: MIP_00261729 (cluster 158 representative) structure comparison between Rosetta, AlphaFold2 and DMPFold (upper models) and the most similar PDB to the AlphaFold2 prediction (lower structure). Mutual TM-scores are shown in between. Additional parameters are listed in the table.*

- As in *Novel fold example* #2, the first 6 residues in N-terminus are disordered and therefore predicted differently by all methods but it doesn't influence the results
- pTM (global score) for AlphaFold2: 0.88

##### 8.7. Largest novel fold clusters

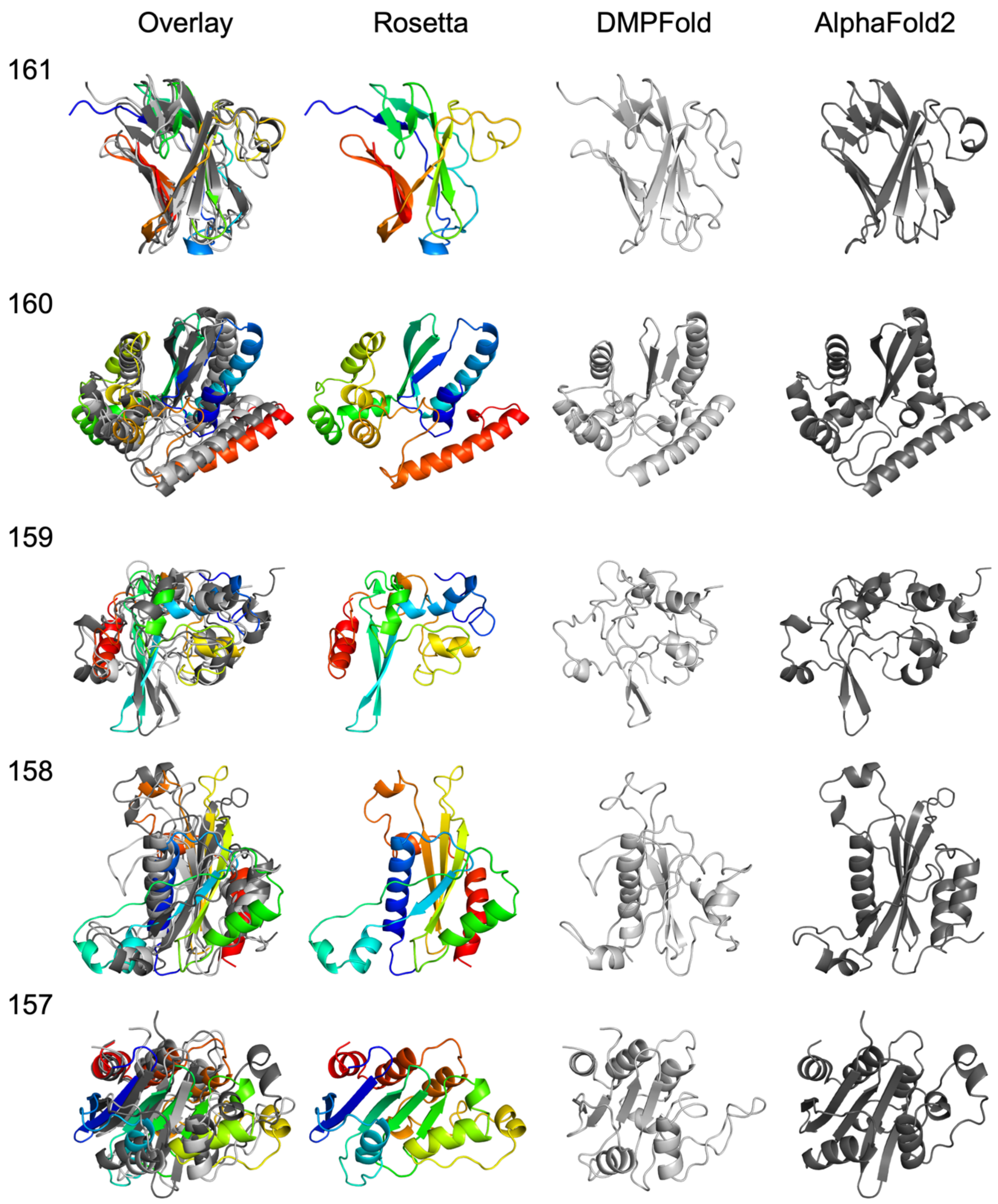

*Fig. S56: Comparison of modeling methods for the largest novel fold clusters.*

#### 8.8. All novel fold clusters

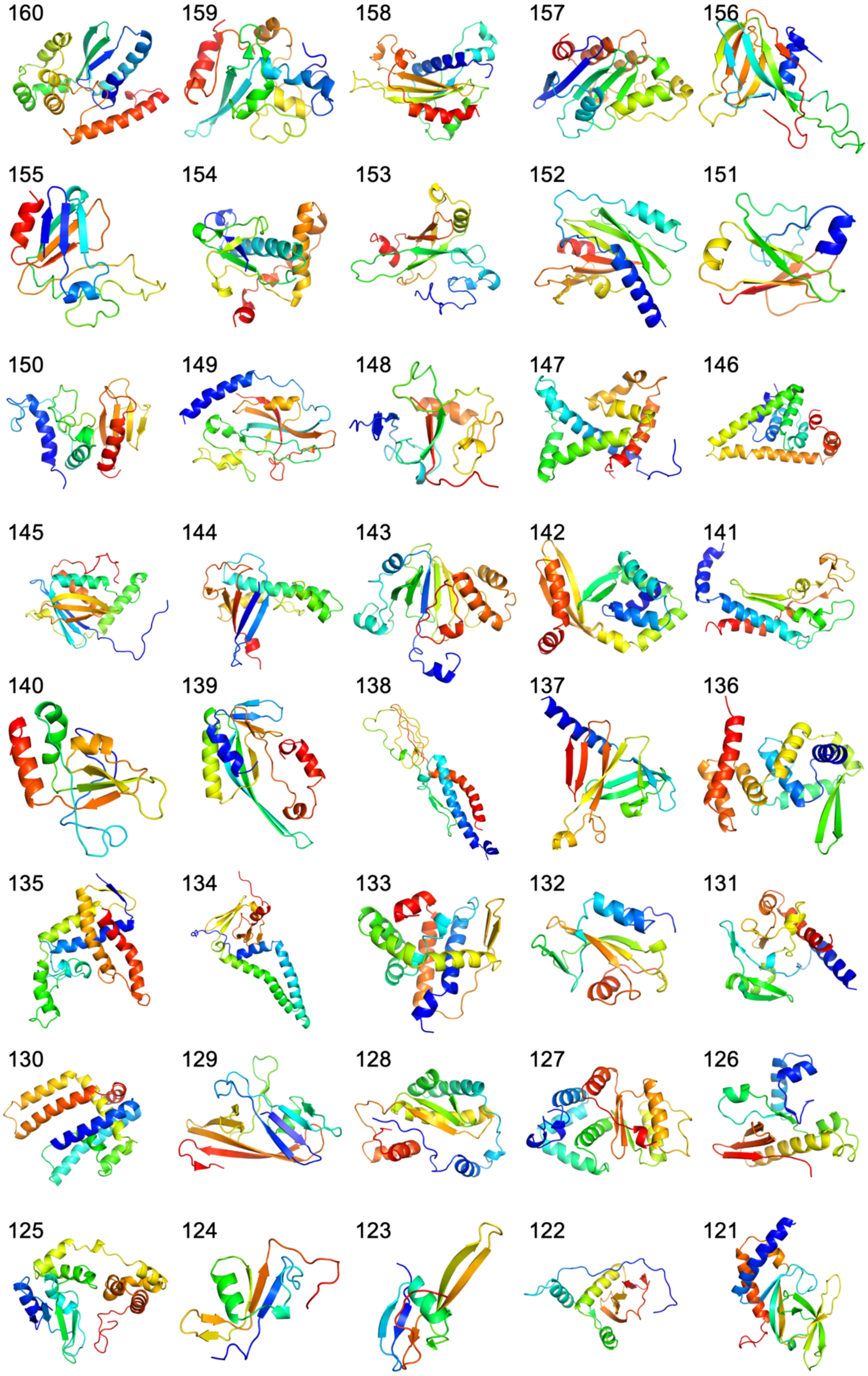

*Fig. S57: Representative Rosetta models for all novel fold clusters (numbered).Figures with red background are hard fals positives and with yellow background are soft false positives.*

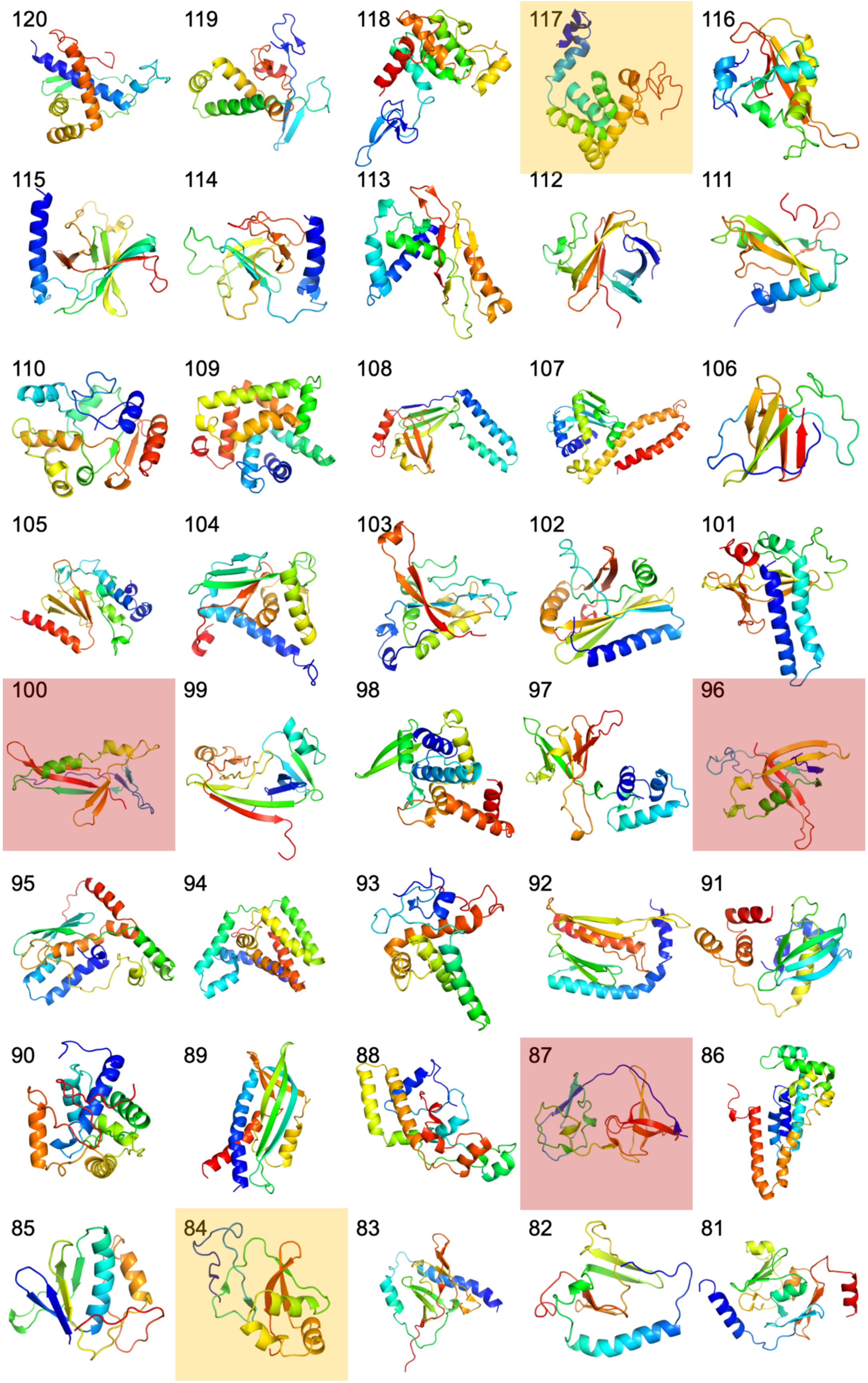

*Fig. S57 continued*

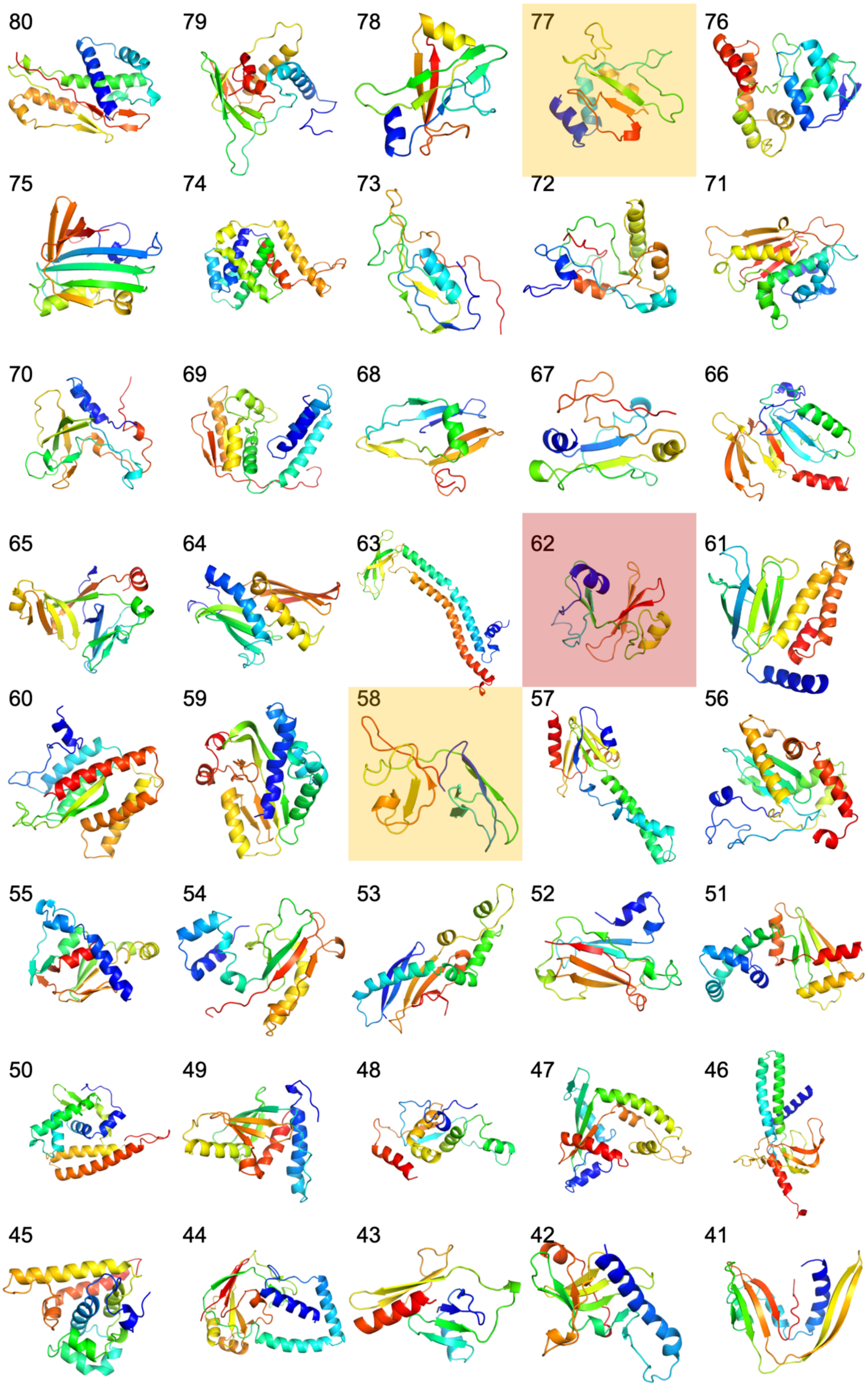

*Fig. S57 continued*

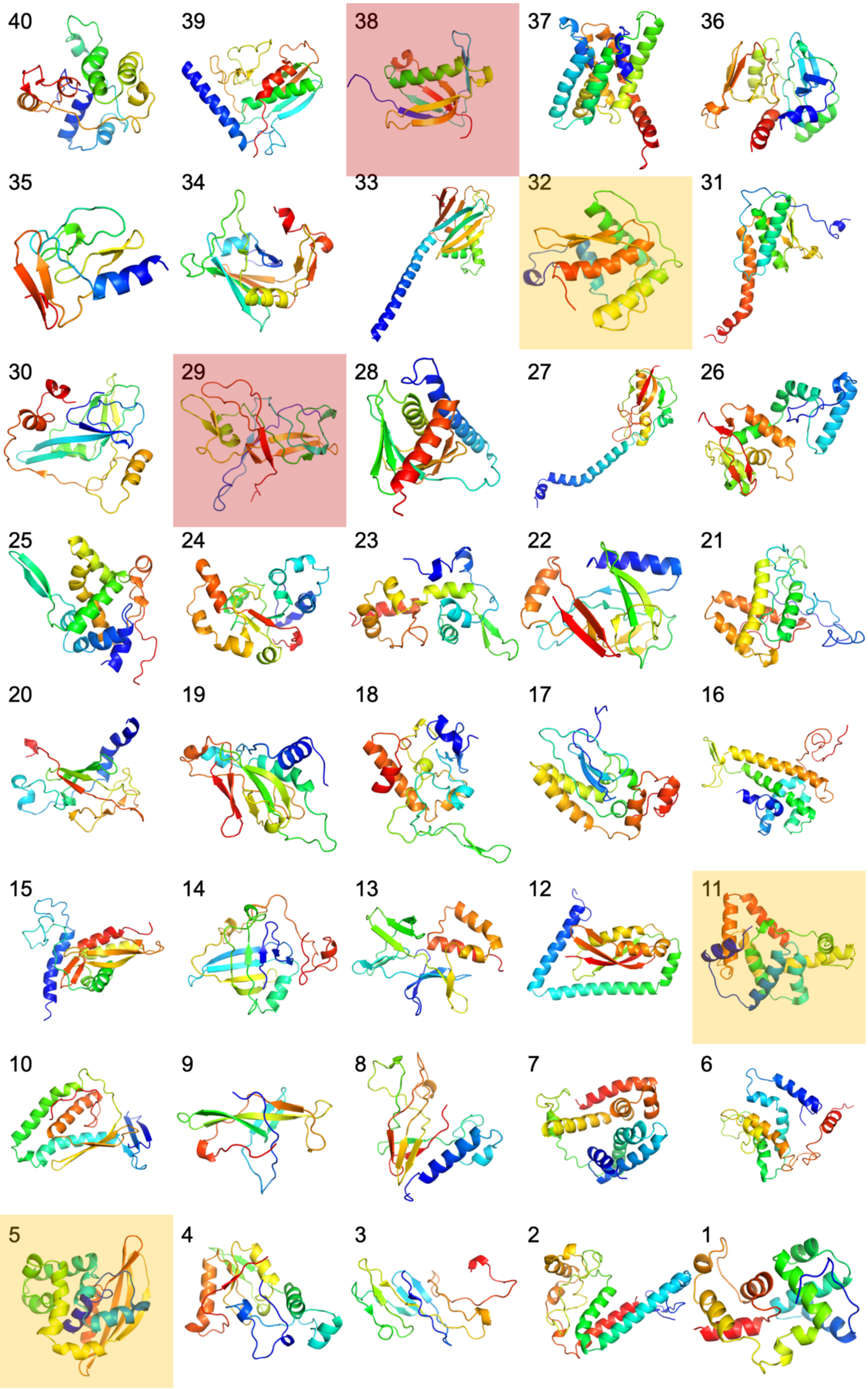

*Fig. S57 continued*

#### 8.9. Functions of the novel fold proteins

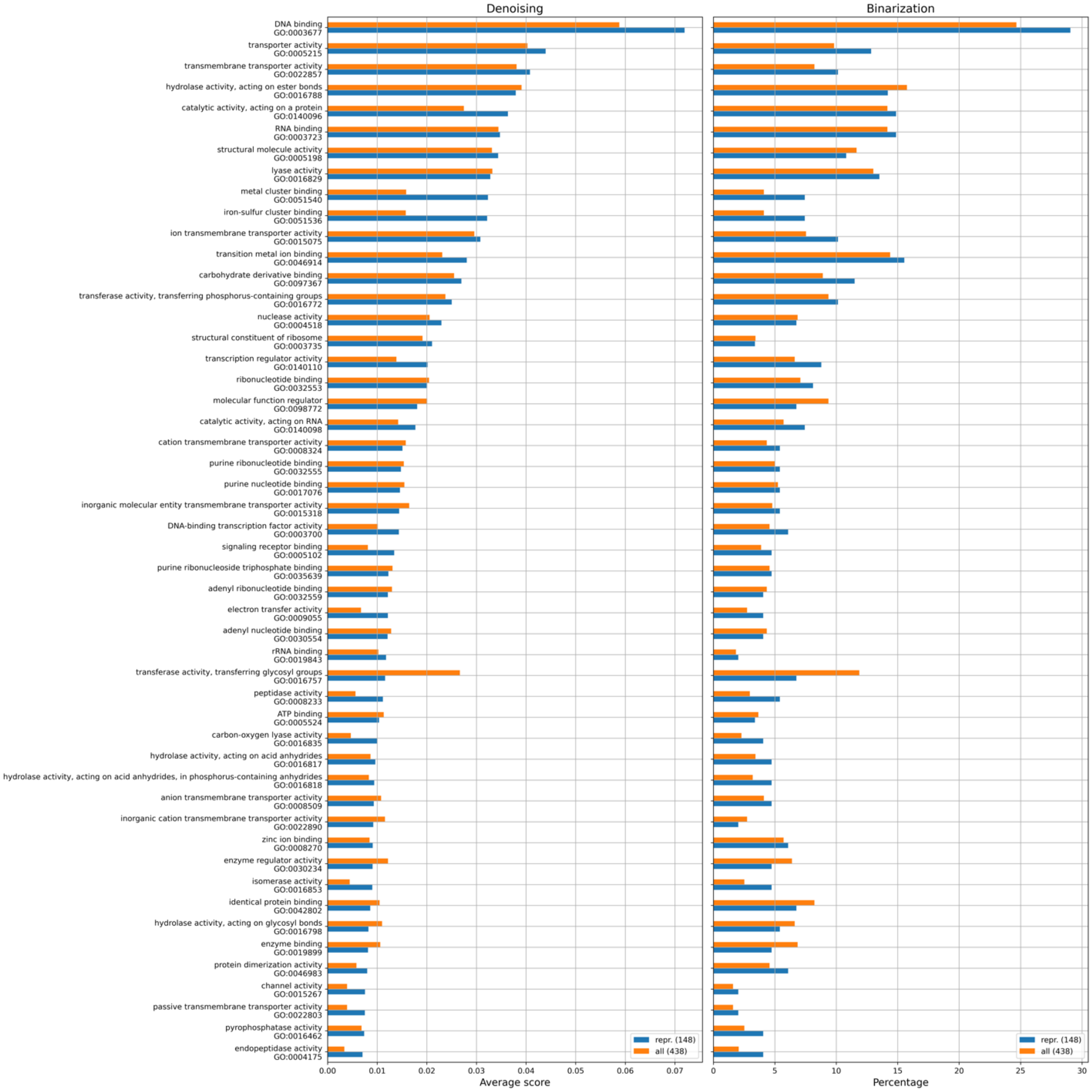

*Fig. S58: The 50 most prevalent functions for the novel fold dataset, covering either the 148 representatives for all novel fold clusters (blue) or all 438 models (orange). For the histogram in the left panel, the function vectors were denoised with a score cutoff of 0.1, then summed over the models in the novel fold dataset and then sorted from the largest to the smallest scores. This means that the left histogram takes into account the magnitude of the scores. For the histogram on the right, the score vectors were binarized: scores $\leq 0.1$ were set to 0 and scores $> 0.1$ were set to 1. Score vectors were summed over the models in the novel fold dataset and then sorted from the largest to the smallest scores. This means that the histogram on the right does not consider the magnitude of the scores but rather the prevalence of these particular functions.*

### 9. Protein universe visualization

The details about how we embedded the models from the MIP visualization dataset into the 3D structure space are described in the Methods section of the main paper - an overview is shown in Fig. S59. In the following sections we show visualizations of the protein universe using both UMAP and Principal Component Analysis (PCA), color coded by various protein features. Each dot in the protein universe represents one model from the MIP visualization dataset.

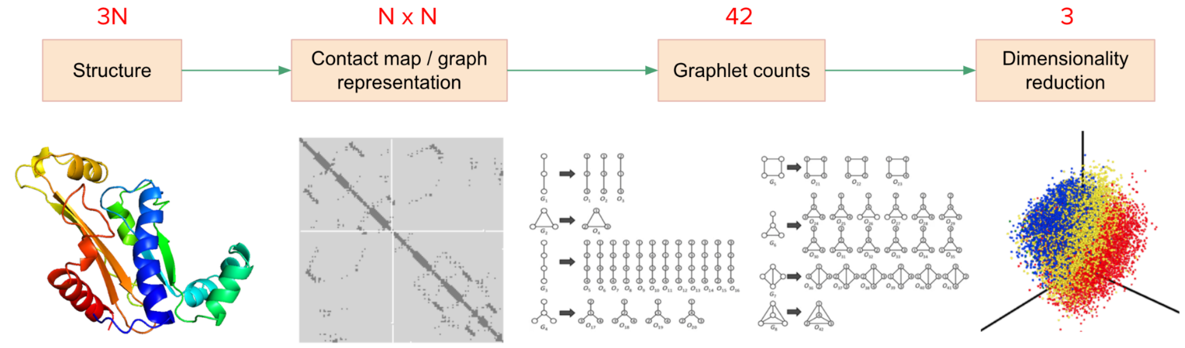

*Fig. S59: An overview of the protein visualization pipeline. N denotes the number of residues in the structure. The graphlet count image was reprinted without changes and with permission from reference[31], which was published under a Creative Commons license (https://creativecommons.org/licenses/by/4.0/).*

#### 9.1. Protein features mapped onto the protein universe visualization

##### *Sequence length*

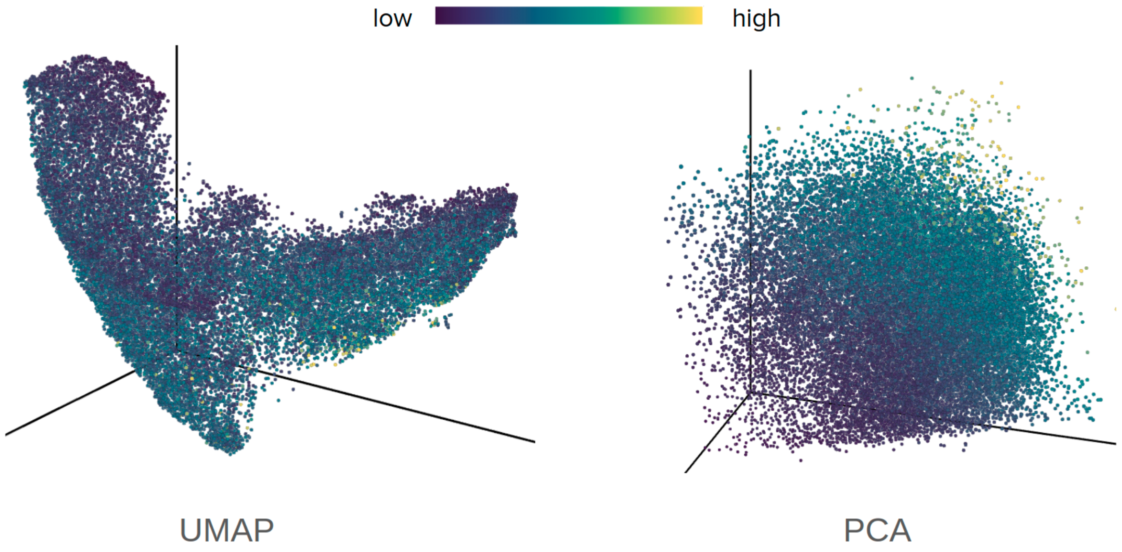

*Fig. S60: 3D structure space visualization for the sequence length.*

- Strong gradient in sequence length might be observed (especially for PCA).
- The largest proteins are placed in the bottom right in the UMAP plot and in the top right in the PCA plot (they correspond to CATH proteins with > 1000 residues).

##### *CATH classes*

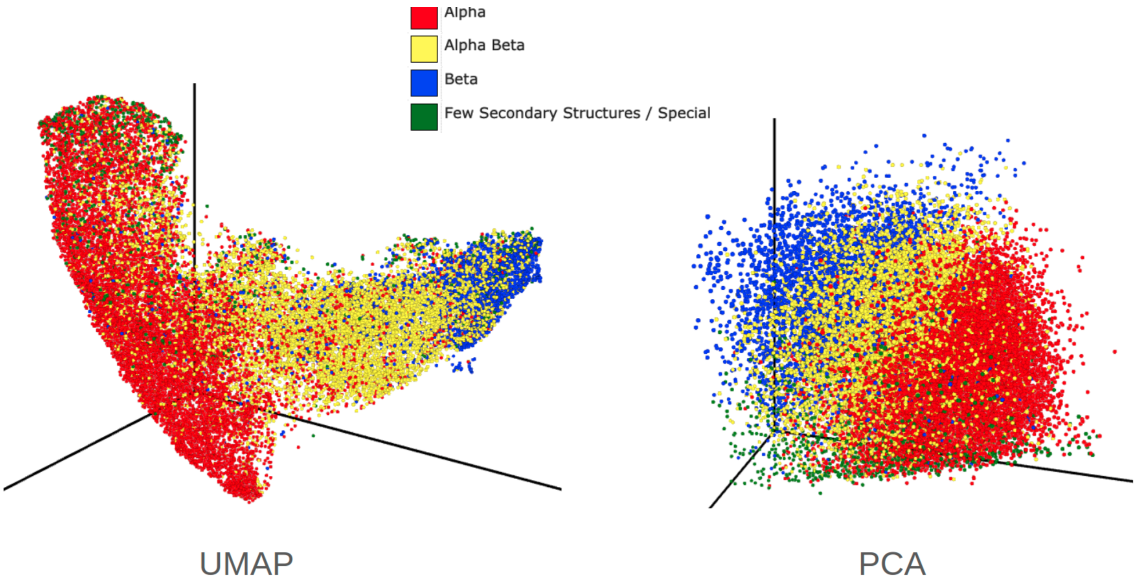

*Fig. S61: 3D structure space visualization for the CATH classes.*

- Structure space is continuous (general conclusion to all plots).
- Strong separation between CATH classes.
- Majority of green points (corresponding to 4th and 6th CATH classes) come from CATH structures - few Rosetta and DMPFold models were annotated this way.
- Majority of points on the outskirts in the PCA plot (in green) are also mainly from CATH.

##### *Relative contact order*

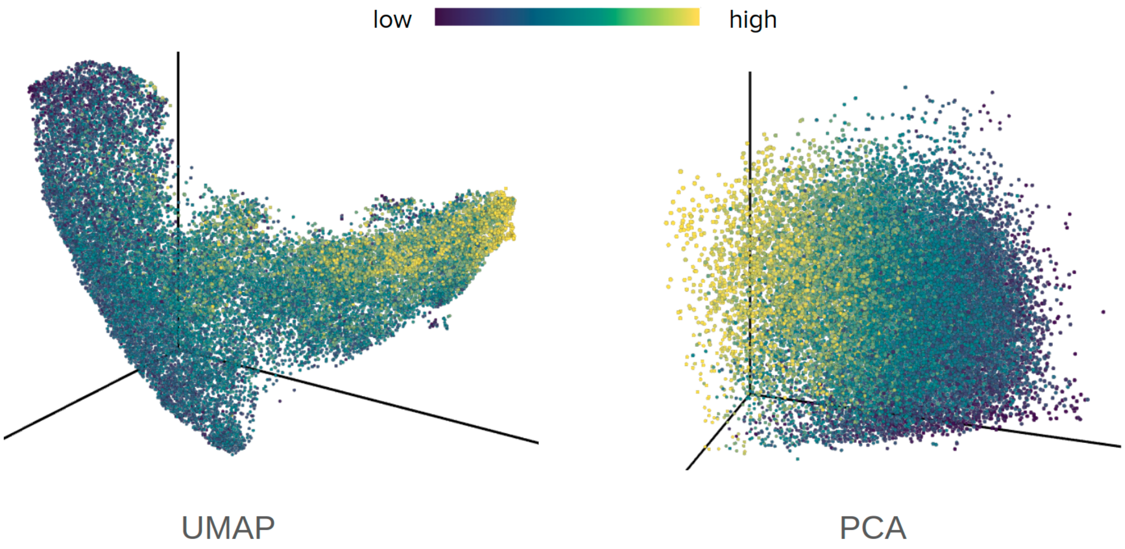

*Fig. S62: 3D structure space visualization for relative contact order.*

- Clear correlation between CATH class and relative contact order. Proteins with higher strand content generally have a higher relative contact order due to long-range contacts, which are fewer in alpha-helical proteins.

##### *Alpha-helical transmembrane spans*

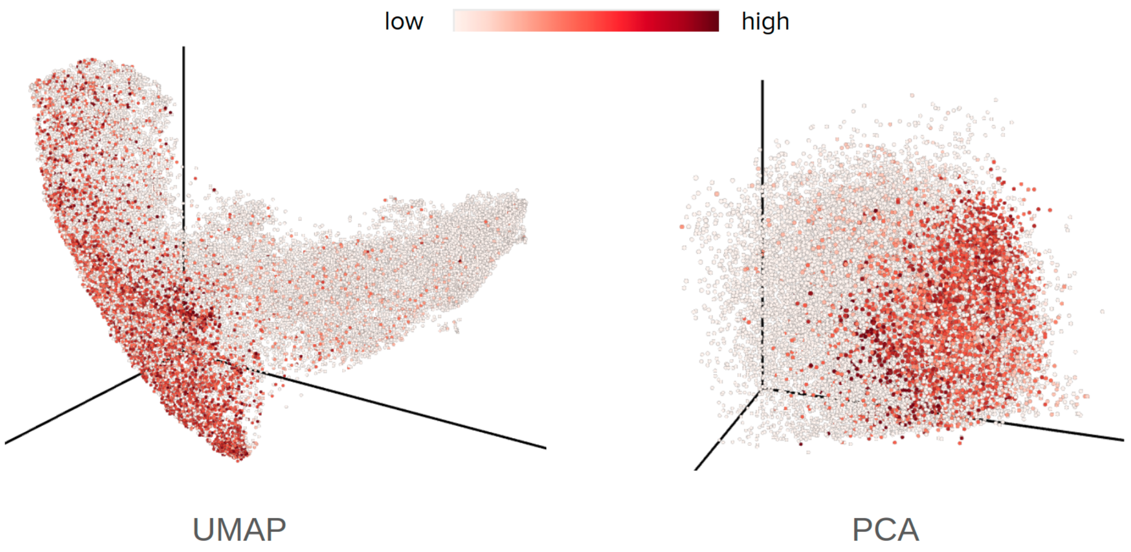

*Fig. S63: 3D structure space visualization for alpha-helical transmembrane spans.*

##### *Beta-strand transmembrane spans*

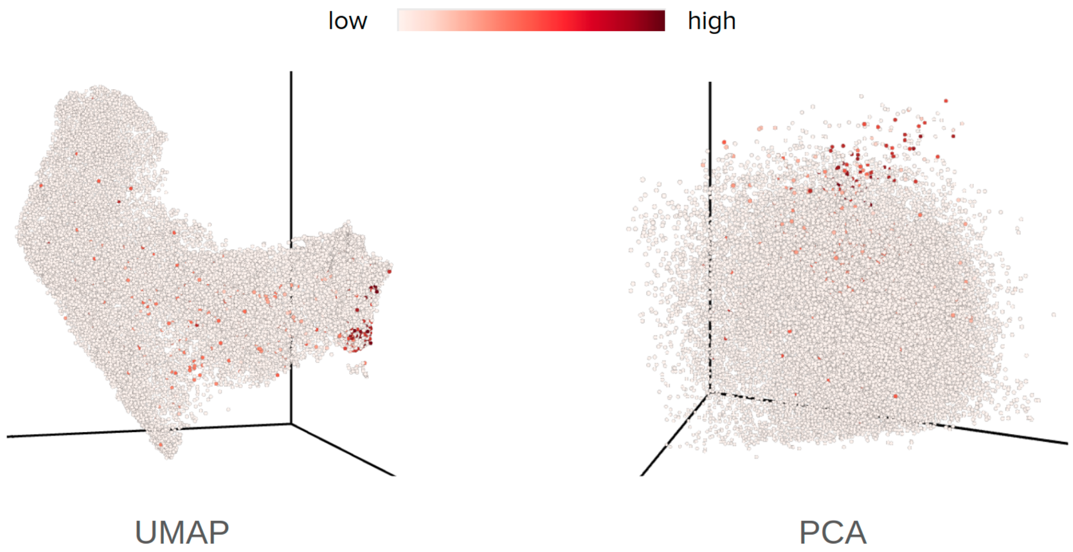

*Fig. S64: 3D structure space visualization: beta-strand transmembrane spans.*

- Alpha/beta transmembrane spans are grouped mainly in the alpha/beta region. Note that alpha/beta transmembrane spans were predicted using OCTOPUS/BOCTOPUS and were predicted using the protein sequence only.

##### *Putative novel folds*

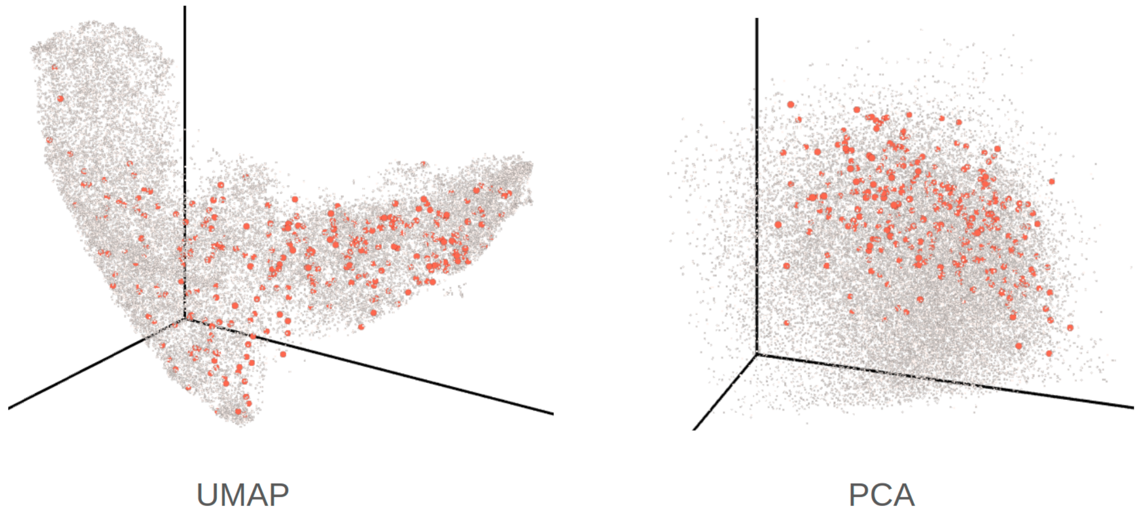

*Fig. S65: 3D structure space visualization for 161 putative novel folds.*

- Putative novel folds are mainly in alpha/beta proteins and cover the entire space with fewer novel folds in purely alpha or purely beta proteins.

#### 9.2. Sample structures

In order to have a better intuition of what structures are located in specific parts of our UMAP and PCA projections, in Fig. S66 we identified positions of our novel fold examples and denoted a few representative structures on the edges (presented in Fig. S67 and Fig. S68).

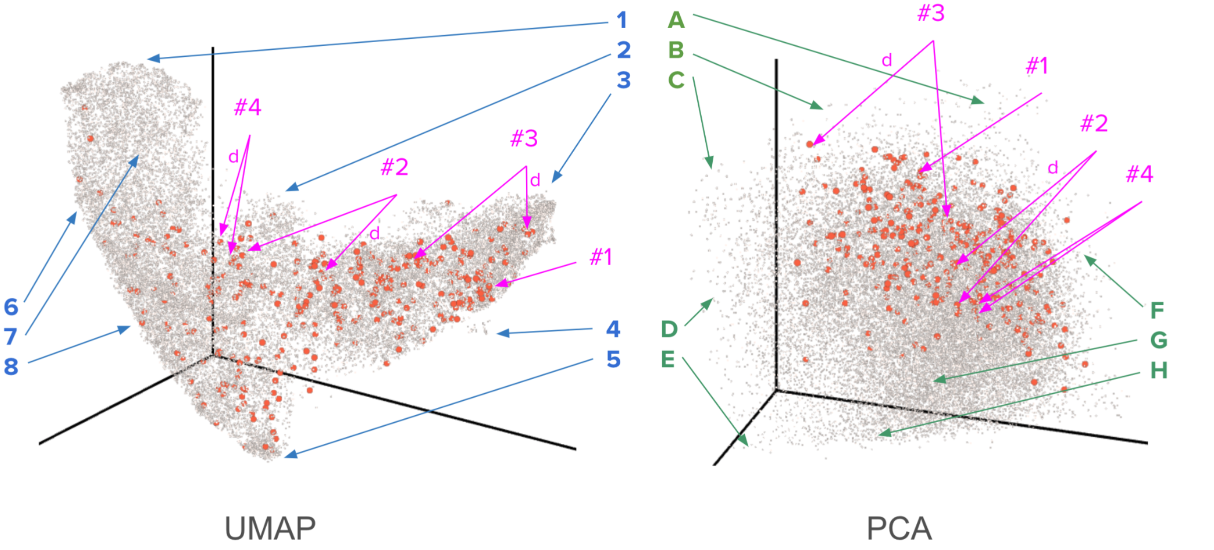

*Fig. S66: 3D structure space visualization: sample structures (shown in Fig. S67 and Fig. S68) among putative novel folds. Blue and green arrows correspond to sample structures placed on the edges in UMAP and PCA projections respectively. Pink arrows point to putative novel fold examples in Fig. S52 – S55 (the arrow labelled with 'd' points to the DMPFold model, the other arrow points to the Rosetta model).*

The Rosetta and DMPFold models are close in 3D space for the novel fold examples #1 and #4. Larger distances are observed for the novel fold examples #2 and #3 using both methods UMAP and PCA. For example #3 the larger distance is likely due to the N-terminal alpha helix position which significantly changes the graphlets counts. Importantly, examples shown in Fig. S67 and Fig. S68 agree with our expectations with respect to alpha/beta-like structure positions, sequence length and transmembrane protein locations.

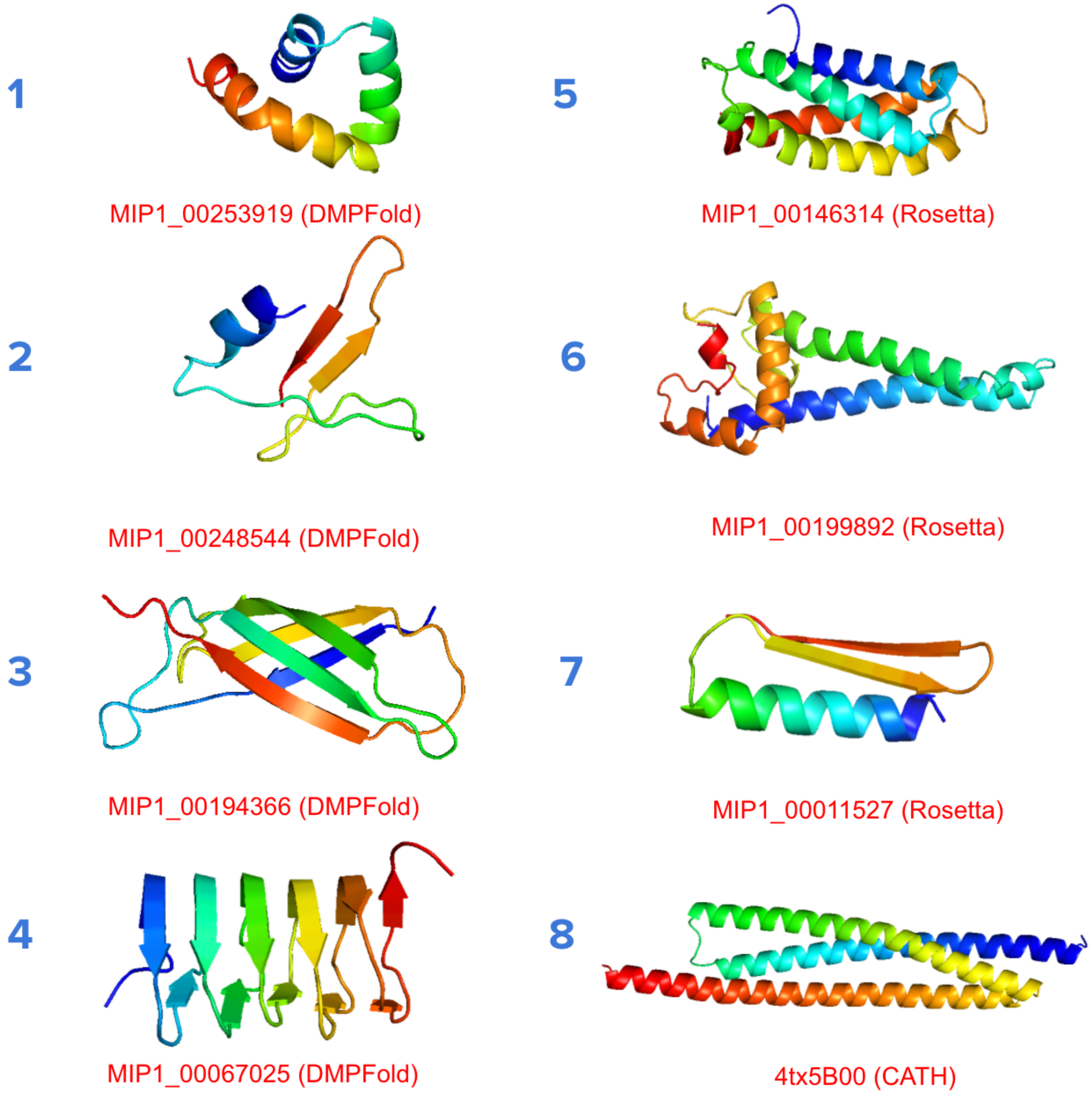

*Fig. S67: Sample structures presented in Fig. S66 (UMAP projection).*

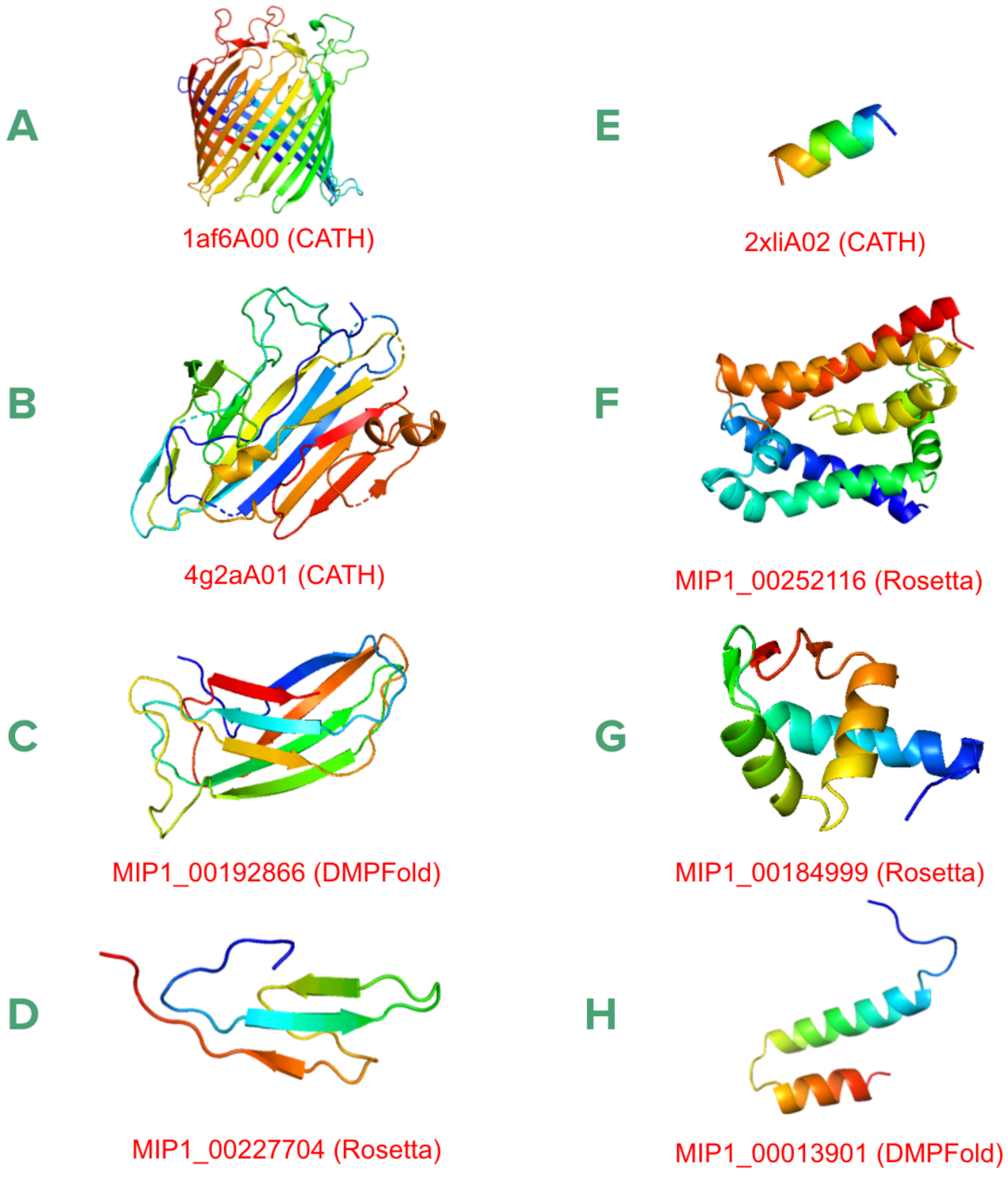

*Fig. S68: Sample structure presented in Fig. S66 (PCA projection).*

#### 9.3. Caveat

Graphlet counts are simple and in some cases perform better than the other, more sophisticated methods[31] but they reflect local rather than global protein geometry. Therefore, when looking at specific examples in UMAP/PCA projections we can find different folds close to each other, which creates potential room for improvements. Nevertheless, this approach is sufficient for our needs and in general agreement TM-score increases if Rosetta and DMPFold models are closer to each other (see Fig. S69). The overall topology created by UMAP visualization should be interpreted with caution, as its shape will depend on starting parameters and stochastic effects. Still, the overall features of this projection do not change with the choice of parameters.

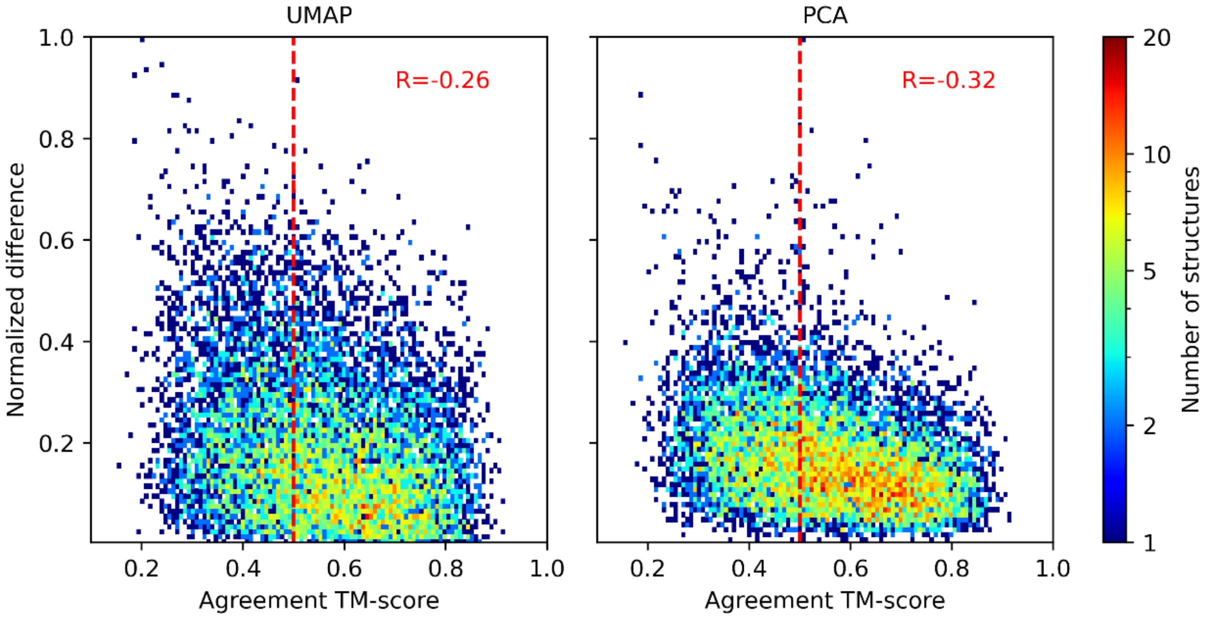

*Fig. S69: Euclidean distance between Rosetta and DMPFold predictions for UMAP (left) and PCA (right) visualizations. Pearson R correlation coefficient is depicted in red in the upper right corners. For easy comparison between PCA and UMAP, the y-axes are normalised by the largest value.*

#### 10. Sanity checks for sequence-structure-function relationships

For initial sanity checks, we compared our *Random5000* data to subsets of the PDB. For this, we constructed two additional datasets by clustering the PDB based on sequence identity:

1. 1000 chains with pairwise sequence identities distributed between 0 and 100% (with 10% step size - 10perc BINS)
2. sequence identity threshold ≤ 20% (20perc THR)

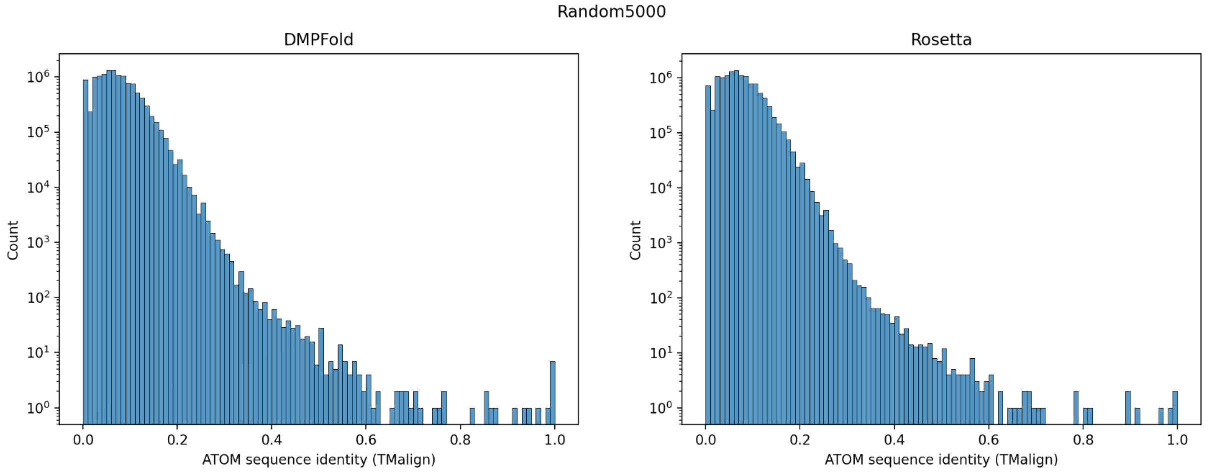

*Fig. S70: By design, our MIP datasets (here from Random5000) exhibit very low pairwise sequence identity. The distribution has a maximum around 10% due to the pairwise nature of the comparison, even though some sequences have substantially higher sequence identity due to domain splitting, which allows few domains to be more similar.*

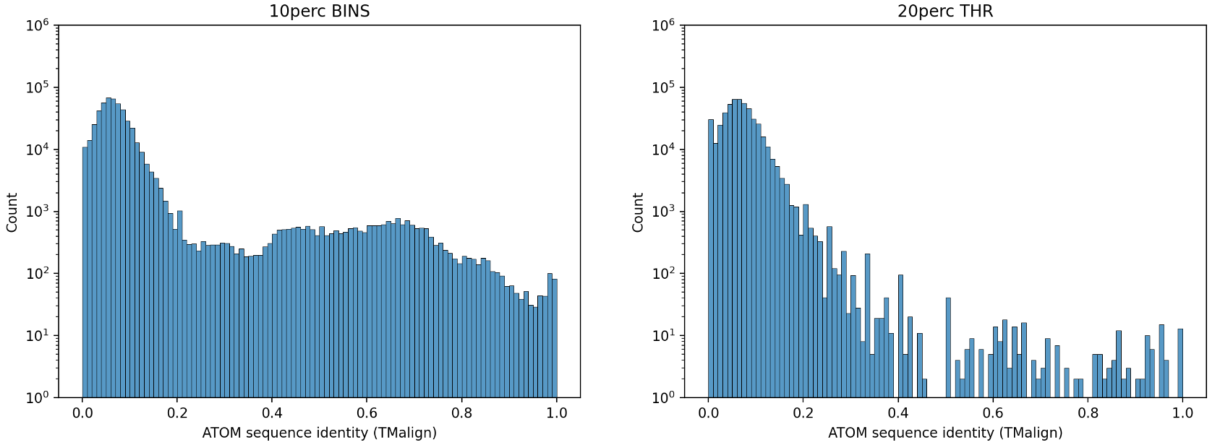

*Fig. S71: Pairwise sequence identities on the subsets of the PDB (Left) that covers the range of sequence identities (10perc bins) and (Right) with a sequence identity cutoff of 20% (20perc thr). The binned subset on the left has higher baseline counts because the pairwise sequence identities cover the whole range.*

#### 10.1. Sequence-structure comparison

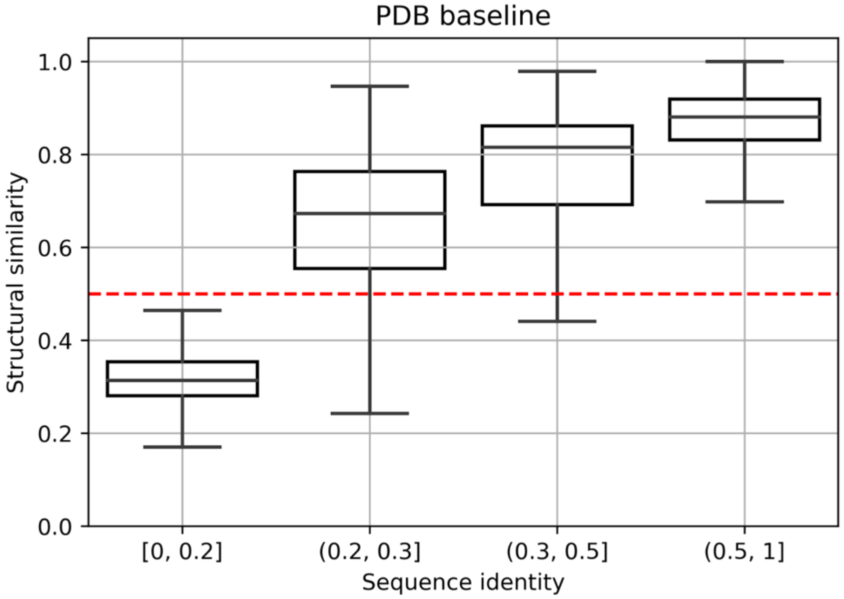

*Fig. S72: Trend between sequence identity and structural similarity (using the TMscore) for the PDB baseline in Fig. 2a.*

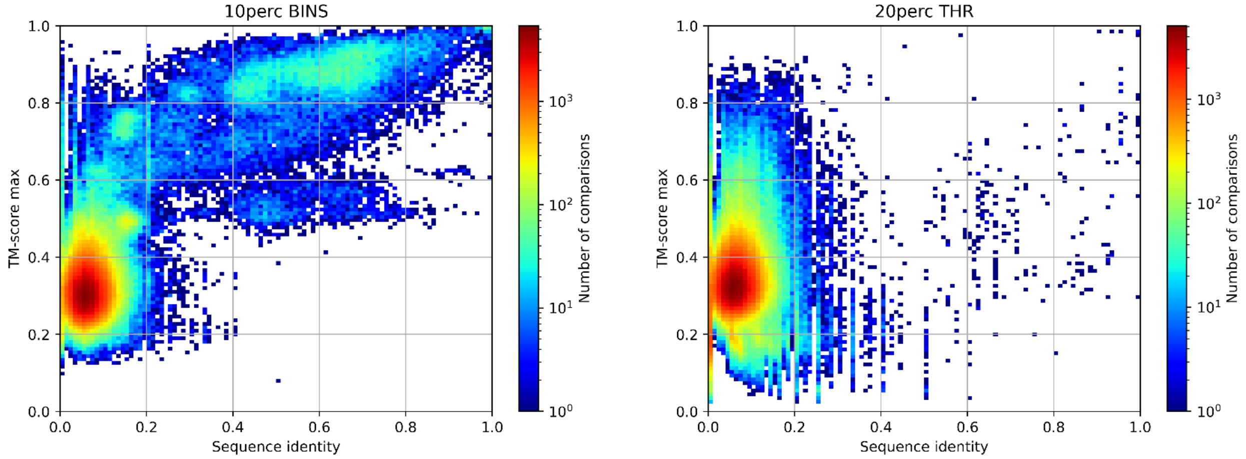

*Fig. S73: Pairwise TM-score plotted over the sequence identity of the two PDB subsets described at the beginning of this section. These plots imply that sequence identity ≤ 0.2 implies low structural similarity and sequence identity > 0.2 implies high structural similarity. The binned subset of the PDB on the left is a more accurate representation of the entire PDB, which covers all pairwise sequence identities. The subset with the threshold is a more accurate representation of our MIP dataset because we picked the sequences specifically to have a low pairwise sequence identity. This is confirmed with comparing the right panel with panel (b) in manuscript Fig. 2.*

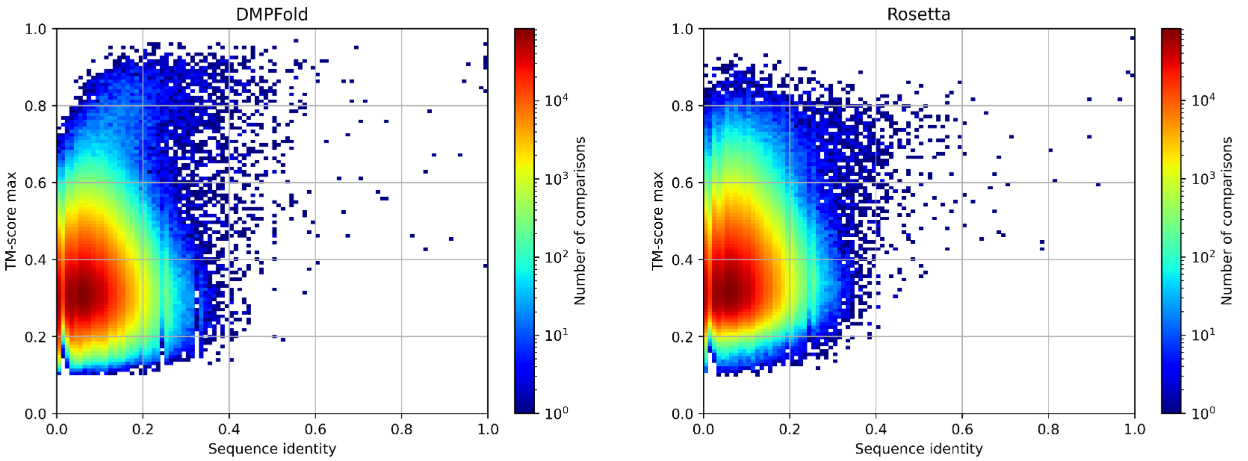

*Fig. S74: Pairwise TM-score plotted over the sequence identity for the DMPFold (left) and Rosetta (right) models in the Random5000 dataset.*

#### 10.2. Sequence-function comparison

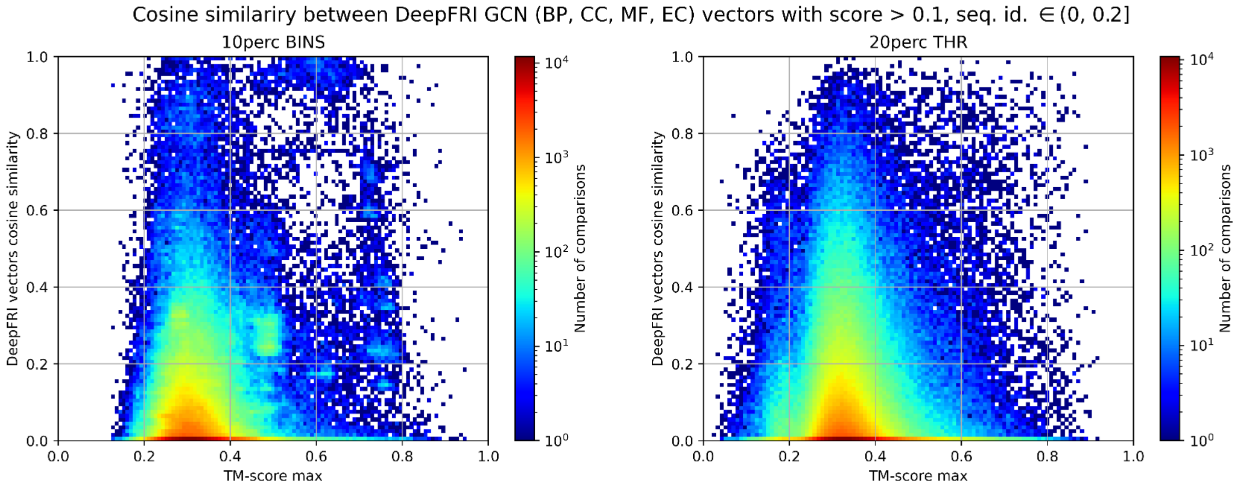

*Fig. S75: Functional similarity (cosine similarity of DeepFRI vectors) plotted over the structural similarity (TM-score) for dissimilar sequences (sequence identities between 0 and 20%) for the binned PDB subset (left) and the PDB subset with a sequence identity cutoff of 20%. These plots show low structural similarity and therefore low functional similarity for the majority of the datasets.*

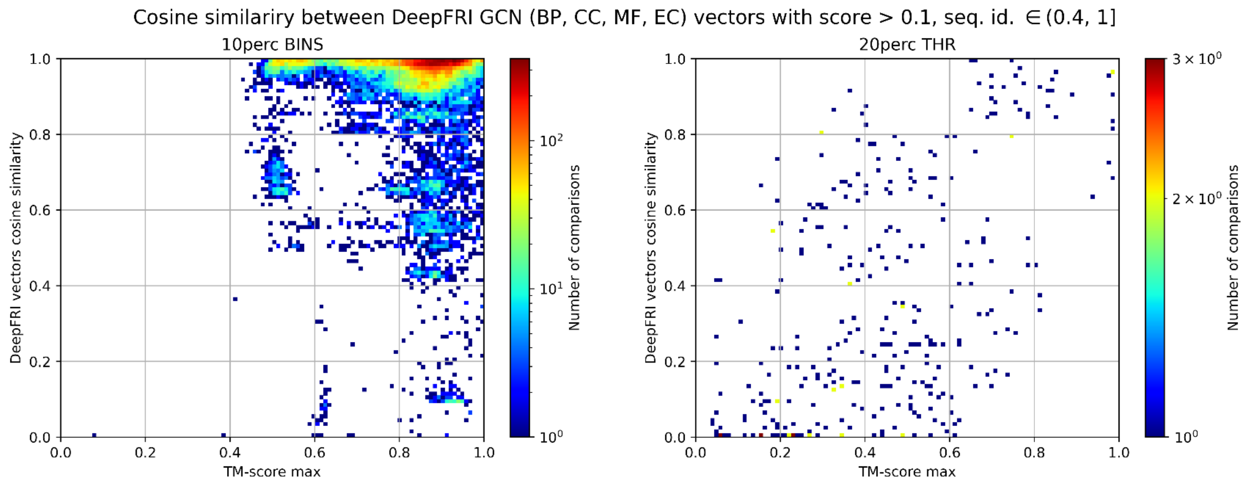

*Fig. S76: Functional similarity (cosine similarity of DeepFRI vectors) plotted over the structural similarity (TM-score) for similar sequences (sequence identities between 40 and 100%) for the binned PDB subset (left) and the PDB subset with a sequence identity cutoff of 20%. These plots show low structural similarity and therefore low functional similarity for the majority of the datasets.*

#### 10.3. Structure-function comparison

We computed structural similarities within clusters of models that have the same functional prediction, i.e. the same GO term. To achieve this, we first grouped MIP models with DeepFRI predictions (score >= 0.2) were by GO-term. Only GO-terms that have 50 - 1000 models within *MIP_curated* were selected. Pairwise structural similarity was computed separately for each GO-term as maximum TM-score of two superimposed MIP structures. We find that structural similarity increases with increasing information content in each GO-group (BP, MF, CC) both for Rosetta and DMPFold predictions (Fig. S77). This is expected because higher information content corresponds to more specific functions which can be represented by a more limited number of folds - therefore their structural similarity is higher.

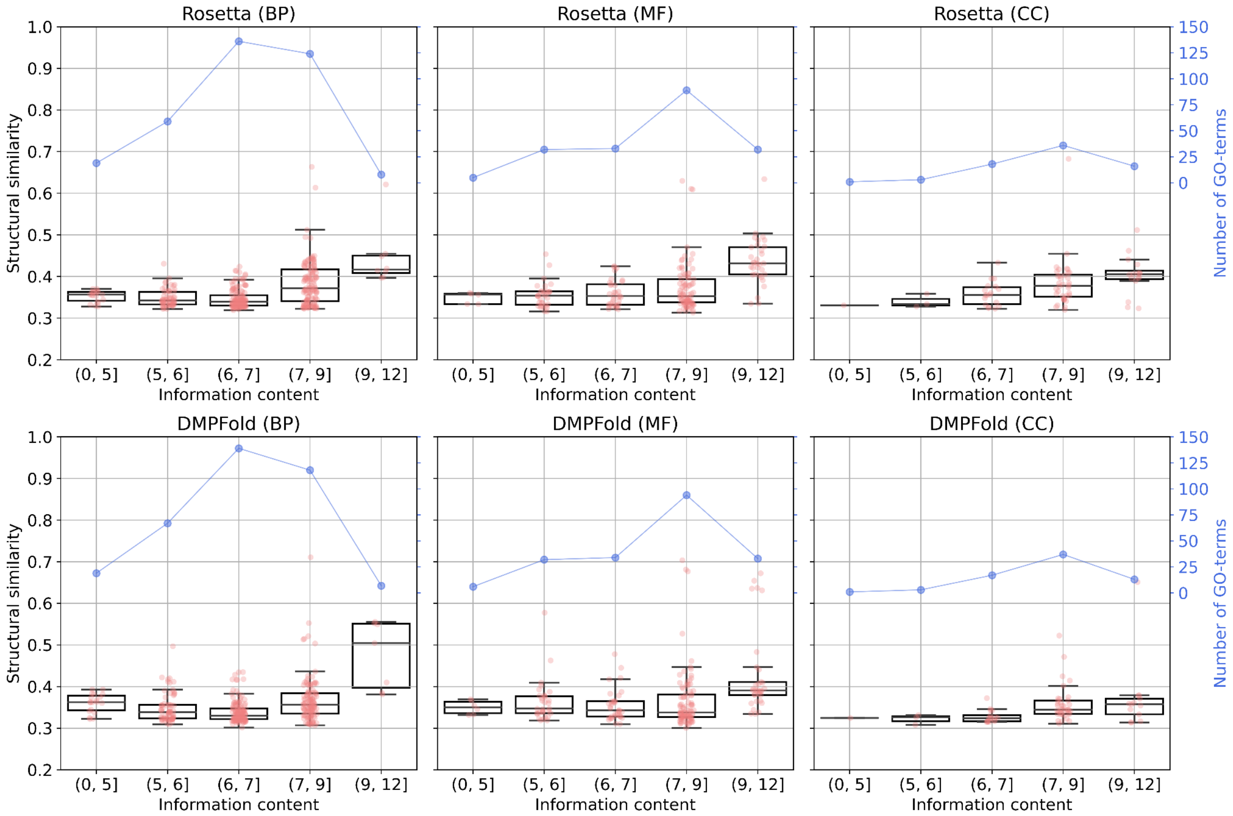

*Fig. S77: Box plots showing structural similarity across GO groups for MIP_curated Rosetta (upper panels) and DMPFold (lower panels) models sorted by increasing information content. Each point on the plot corresponds to the mean of such TM-scores (one point for each GO-term). The blue curve depicts the number of compared GO-terms.*

### 11. Structure Function Motifs

#### 11.1. Functional diversity of proteins with the same structure

Fig. 3 shows examples where we look at the functional diversity of proteins with the same structure. For the proteins in Fig. 3a the TM-scores are 0.55 and 0.70 for Rosetta / DMPFold models, respectively. For the proteins in Fig. 3b the TM-scores are 0.58 / 0.82 for Rosetta / DMPFold models, respectively. High similarities of the contact maps between the two proteins in each example indicate that it is the embedding of the structural motif (white rectangle) in the tertiary structure that is responsible for the function, instead of just the secondary structure. We also show the class activation maps with the functional predictions mapped onto the sequence to highlight the similarities there.

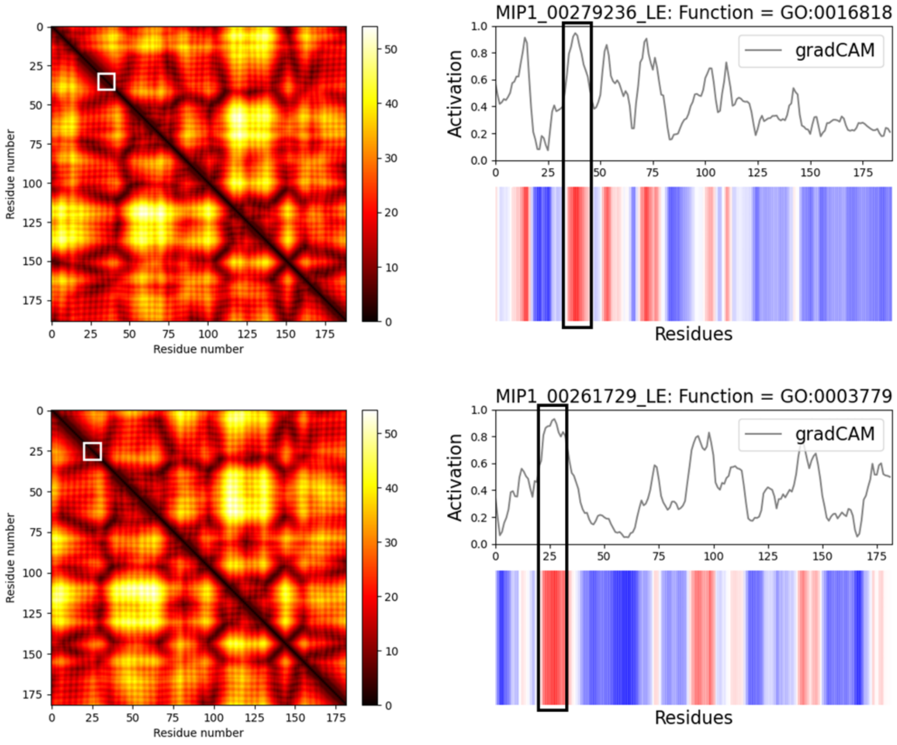

*Fig. S78: Contact maps (left) and class activation maps (right) for both proteins in Fig. 3a. The boxes in white and black highlight the functional motif with the highest function prediction score.*

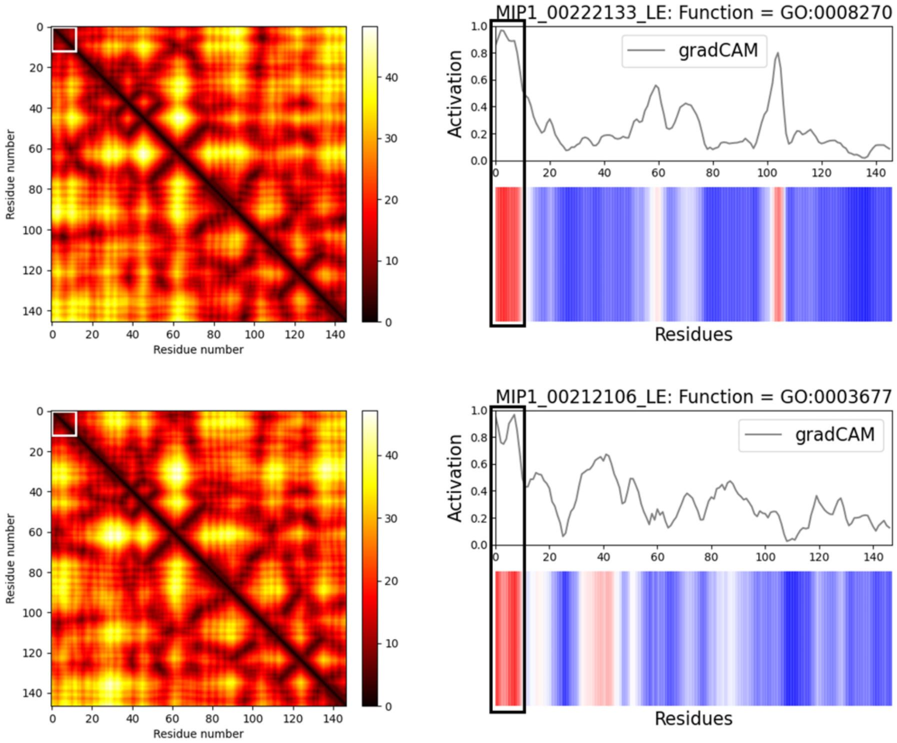

*Fig. S79: Contact maps (left) and class activation maps (right) for both proteins in Fig. 3b. The boxes in white and black highlight the functional motif with the highest function prediction score.*

#### 11.2. Structural diversity of proteins with the same function

In Fig. 4 we highlight several cases where we explored the structural diversity of proteins with the same predicted function. Here we take a closer look at some of the highlighted structures and their functional relationships. For each of the three functions (in panels a, b, and c), Fig. 4 shows structural relationships of the proteins in each functional cluster via a heatmap (through pairwise TM-scores). Specific models are then shown on the right.

#### 11.3. Carbohydrate binding (MF GO:0030246) - Fig. 4a

This functional cluster for carbohydrate binding covers many folds but includes one major fold (cluster shown in (a) as indicated by the yellow square in top-left corner of the heatmap in Fig. 4a. Additionally, this cluster represents the largest cluster of proposed novels folds from this study (example #1 in the *New Fold examples* section above). This molecular function is two levels below the top of the GO hierarchy and has more than 40 child terms. There are ~5,100 entries in the PDB annotated with this label. Carbohydrate binding is an important protein function for all microbes, particularly in gut microbiomes where carbohydrate metabolism benefits both the microbe and its host, as recently reviewed[32].

Carbohydrate binding modules (CBMs) direct carbohydrate-active enzymes (CAZymes) to specific carbohydrate motifs. CAZymes have been cataloged and classified in the CAZy Database[33]. CBMs have been classified based on their type and fold family[34]; the CAZy database currently describes 89 CBM families, which we compared our MIP models to (Fig. S80).

The novel fold cluster has features frequently found in carbohydrate binding modules such as a beta-sandwich topology and clusters of large-hydrophobic residues being indicated as salient for carbohydrate binding functionality (Fig. S81A), but doesn't have high structural similarity to any experimentally determined structures of CBM listed in the CAZy Database (Fig. S80) or any structures annotated as having carbohydrate binding functionality (GO:0030246). It could potentially be a new class of CBM but further analysis and experiments would be required to verify this possibility.

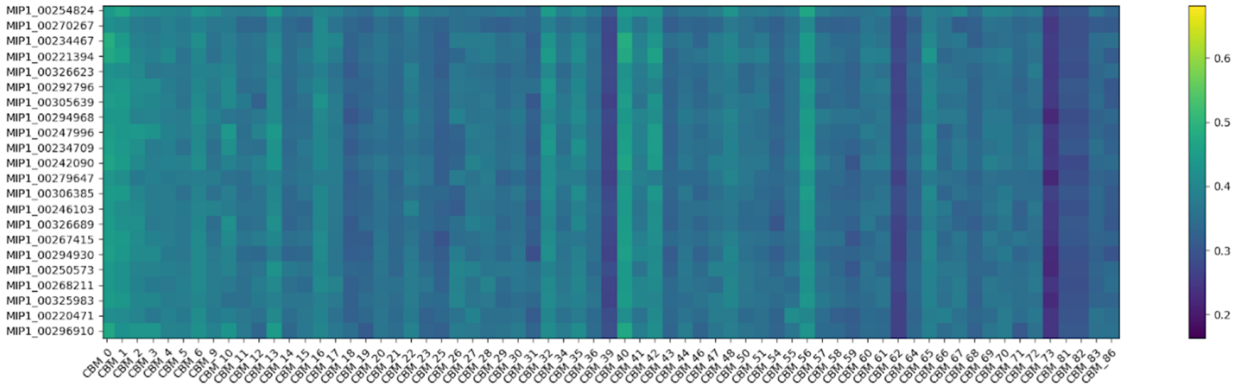

*Fig. S80: Comparing MIP models with known Carbohydrate Binding Modules. Heatmap showing the highest TM-score between Rosetta models of MIP sequences belonging to the novel fold cluster predicted to have carbohydrate binding*

*functionality and a representative of experimentally determined structures of all 89 classes of carbohydrate binding modules listed in the CAZy database.*

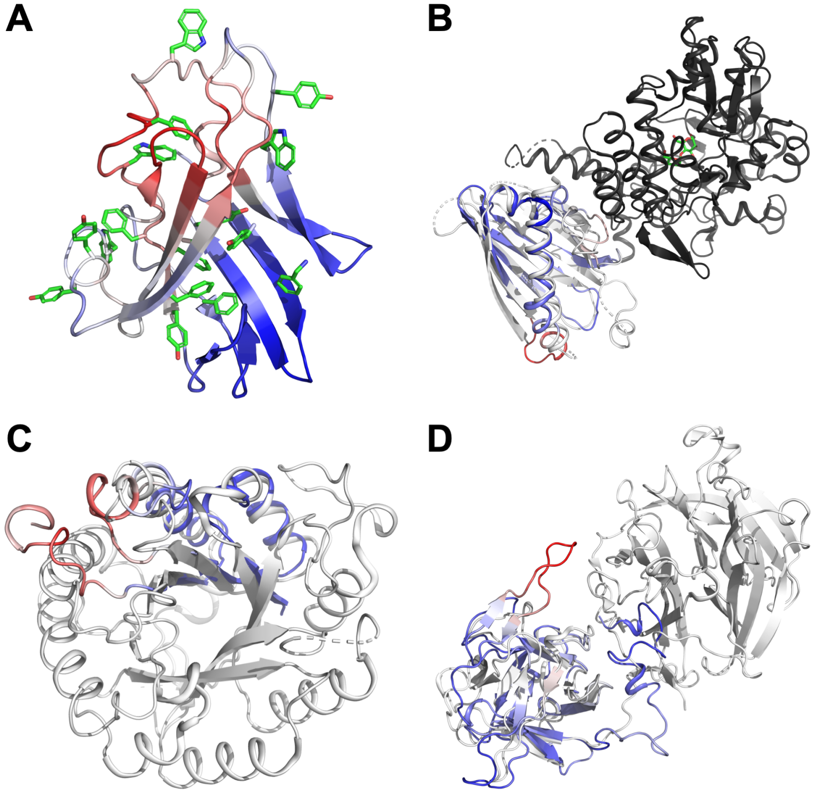

*Fig. S81: (A) Representative structure of the novel fold cluster with PHE, TRP, TYR residues shown as green sticks. As in other Figures, MIP models are colored by DeepFRI score to indicate predicted saliency for the given function. (B - D) Rosetta predicted models of several MIP sequences super imposed on structurally similar experimentally determined structures. Experimentally determined structures used in superpositions are shown white or black. (B) MIP_00224264 superimposed on PDB ID 4XZV chain A (TM-score 0.63) of the SLMO1-TRIAP1 complex[35]. SLOM1 does not bind carbohydrates, but is associated with maltose-binding periplasmic protein TRIAP1. (C) MIP_00094272 superimposed on PDB ID 1LF1 chain A[36] (TM-score 0.64), the catalytic core of a glucoside hydrolyase. (D) MIP_00326772 superimposed on PDB ID 3VT1 chain D[37] (TM-score 0.63). The MIP sequence superimposes with the carbohydrate binding module (type 13) of a glycoside hydrolase.*

#### 11.4. Maintenance of CRISPR repeat elements (BP GO:0043571) - Fig. 4b

This functional cluster covers several folds but includes one major fold (cluster shown in (A) as indicated by the yellow square in the heatmap in Fig. 4b). This biological process has no child terms in the GO hierarchy and there are ~200 PDB entries with this label. The CRISPR sequences are DNA sequences found in the genomes of prokaryotic organisms like bacteria and archaea and they derive from DNA fragments of bacteriophages that previously infected those organisms. Hence, the CRISPR-Cas system functions as a microbial immune system and it is not surprising to find CRISPR sequences in our MIP database. There are few gene products associated with this function and they are associated with Cas1,2,5,6,9,D,and A. Cas1 and Cas2 identify the site in the bacterial genome where they insert viral DNA for later identification and ultimately cleavage by Cas9. As an example, the structure with PDB ID 5XVP[38] has four copies of Cas1 (chains

ABCD) and two copies of Cas2 (chains EF); Cas1 and Cas2 have a different fold. The structural cluster in (A) overlays with a large part of Cas2 and the salient residues highlighted by the function prediction DeepFRI bind DNA in the structure. Clusters marked (D) and (E) in Fig. 4b are similar to parts of Cas1 but don't overlay perfectly. Fig. S82 below shows the Rosetta predicted models of MIP_00052663 superimposed on 5XVP chain F and MIP_00011079 superimposed on 5XVP chain C.

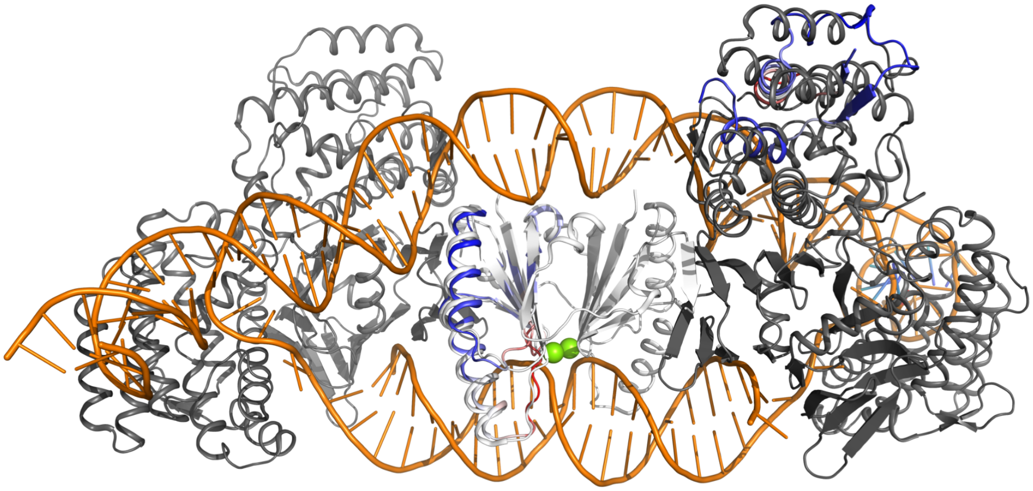

*Fig. S82: MIP_00052663 aligned to 5XVP chain F (TM-score 0.87) and MIP_00011079 aligned to 5XVP chain C (TM-score 0.47).*

##### 11.5. Sole sub-class for lyases that do not belong in the other subclasses (EC 4.99.1.-) - Fig. 4c

As shown in Fig. 4c, all models in this functional cluster have the same fold and the salient residues overlap nicely across all structures. Lyases are enzymes that catalyze the breaking of chemical bonds by means other than hydrolysis or oxidation. Because they break bonds, lyases only require a single substrate, unlike for the reverse reaction, which requires two substrates. The E.C.4. class of enzymes are all lyases that cleave carbon-carbon bonds (E.C.4.1.), carbon-oxygen bonds (E.C.4.2.), carbon-nitrogen bonds (E.C.4.3.), carbon-sulfur bonds (E.C.4.4.), carbon-halide bonds (E.C.4.5.), and phosphorus-oxygen bonds (E.C.4.6.). The E.C.4.99.1.- class of lyases is therefore the "sole subclass for lyases that do not belong in the other subclasses" and covers several chelatases for iron, nickel, cobalt, and magnesium, dehydratases, a lyase and a ligase. None of the lyases in the other classes with known structures has the same fold as our predicted MIP models, even though there are structural similarities. Our models have an (αβ)x3 fold with sequential strand connections the other lyases have various (αβ)xN folds but their strand connections are non-sequential, with some strands inserting between existing ones. Fig. S83 below shows the Rosetta prediction of MIP_00077627 superimposed with PDB ID 5Z7T chain B[39]. The experimentally determined structure is of SirB, a ferrochelatase from *Bacillus subtilis*. The salient esidues for the function predicted by DeepFRI superimpose to residues in the experimentally determined structure that coordinate a Cobalt ion with which it was co-crystalized.

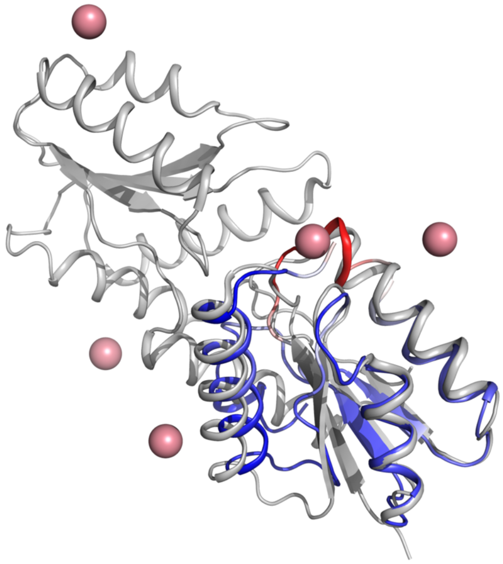

*Fig. S83: MIP_00077627 aligned to 5Z7T chain B (TM-score 0.77).*

#### 12. Structure-to-function examples: comparing functions for novel fold structural clusters

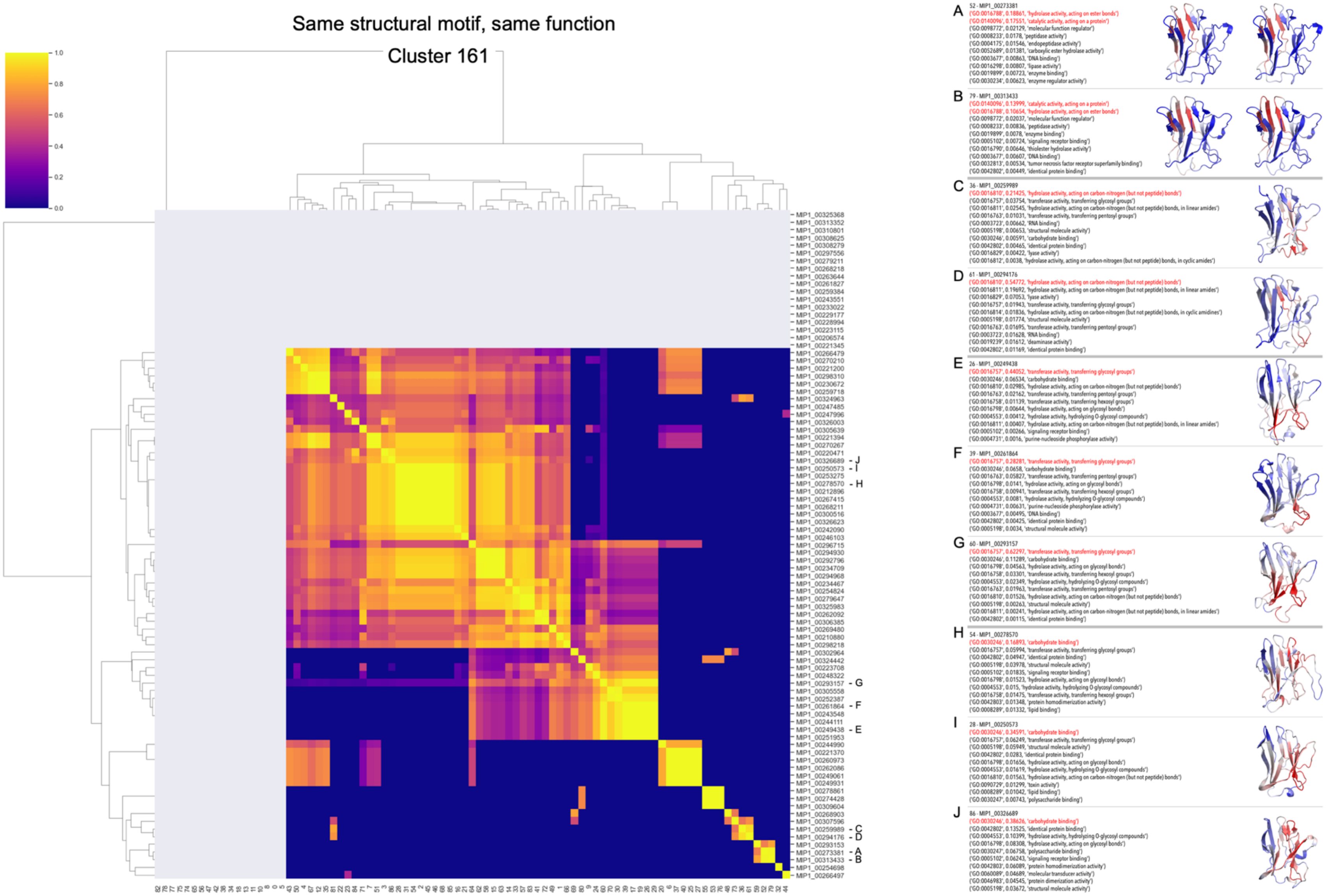

*Fig. S84: Comparison between functions across proteins for the largest novel-fold cluster. The 452 proteins with previously unseen structures were clustered into 161 folds. Here we show the largest of the structural clusters, which has 87 representatives. The heatmap on the left shows the functional similarity (cosine similarity of function vectors) between protein pairs in this cluster. The majority of proteins (in this cluster and in general) follow the well-known observation that similar structures generate similar functions. When mapping residues-specific functions onto the protein structures (residues responsible for the function in red), one can see that different structural motifs are responsible for different functions - panel on the right.*

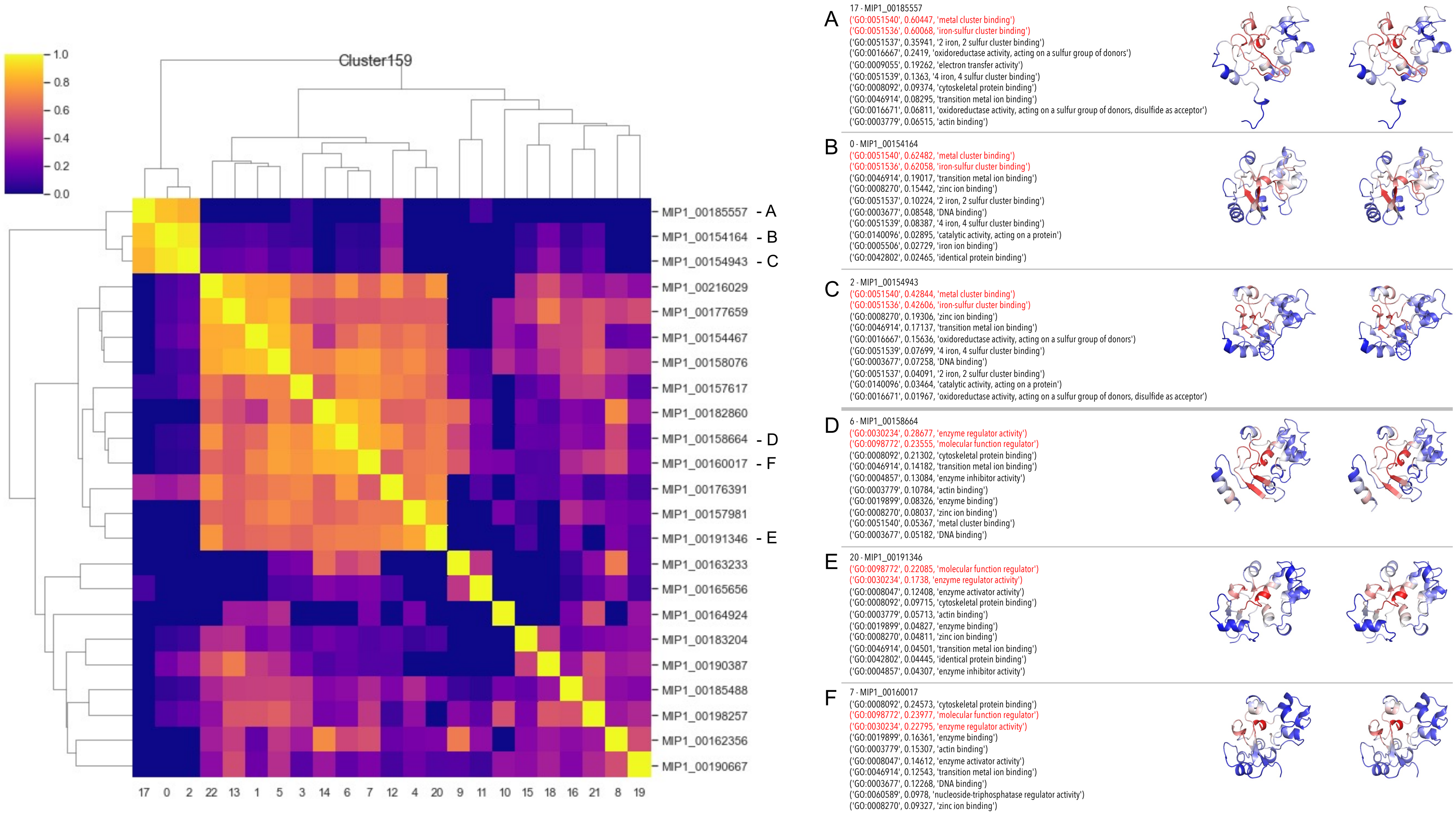

*Fig. S85. Cluster159 with examples of similar structural motifs that produce similar functions. A, B, and C regulate iron-sulfur cluster binding which is a child term of metal cluster binding, and D, E, and F have molecular function regulator activity which includes enzyme regulator activity.*

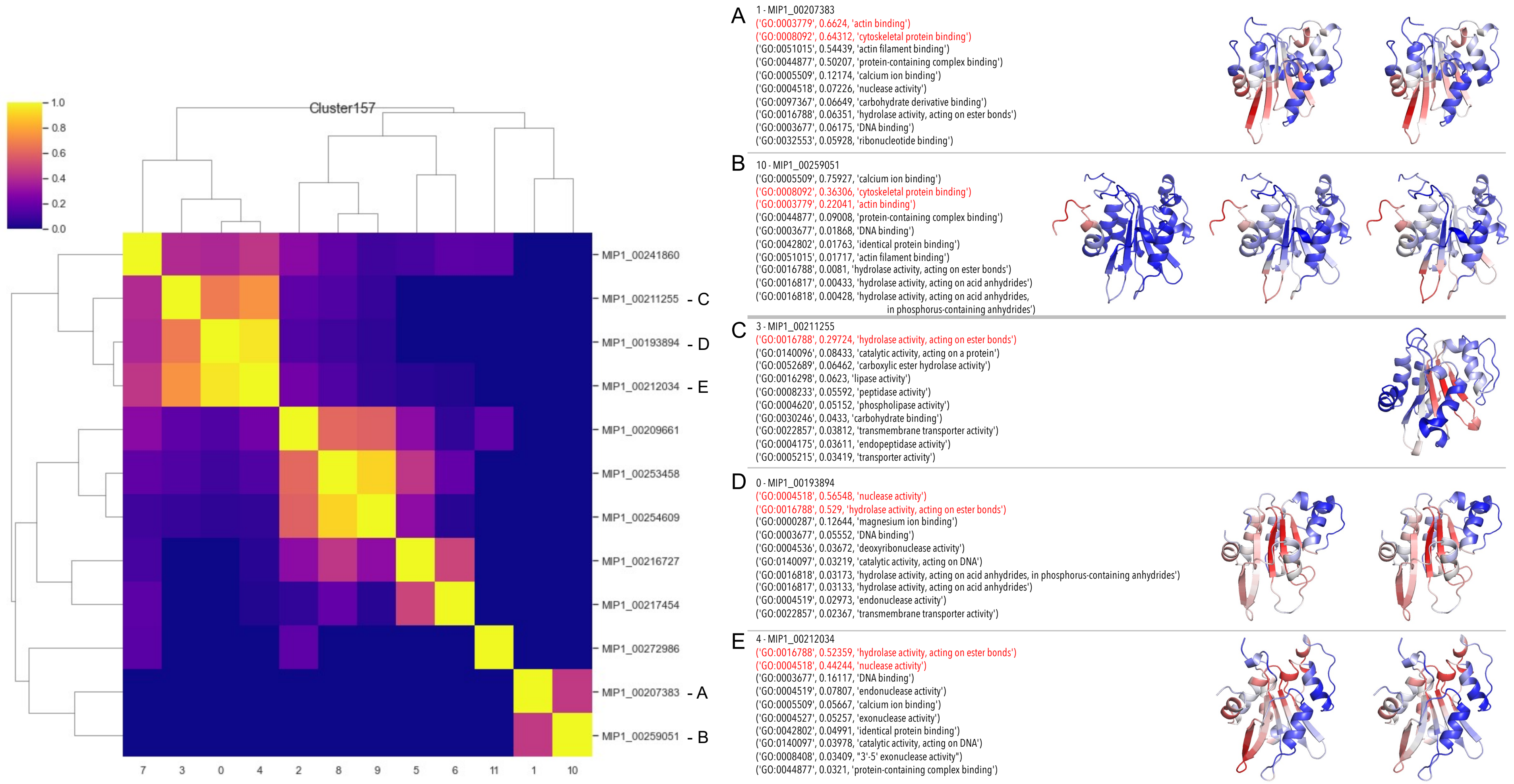

*Fig. S86. Cluster 157 with examples of structure-function disparity. For A and B the same functions (cytoskeletal protein binding, which includes actin binding) can be produced by different structural motifs. For A, D and E the same structural motif produce different functions (cytoskeletal protein binding / actin binding and hydrolase activity, acting on ester bonds / nuclease activity).*

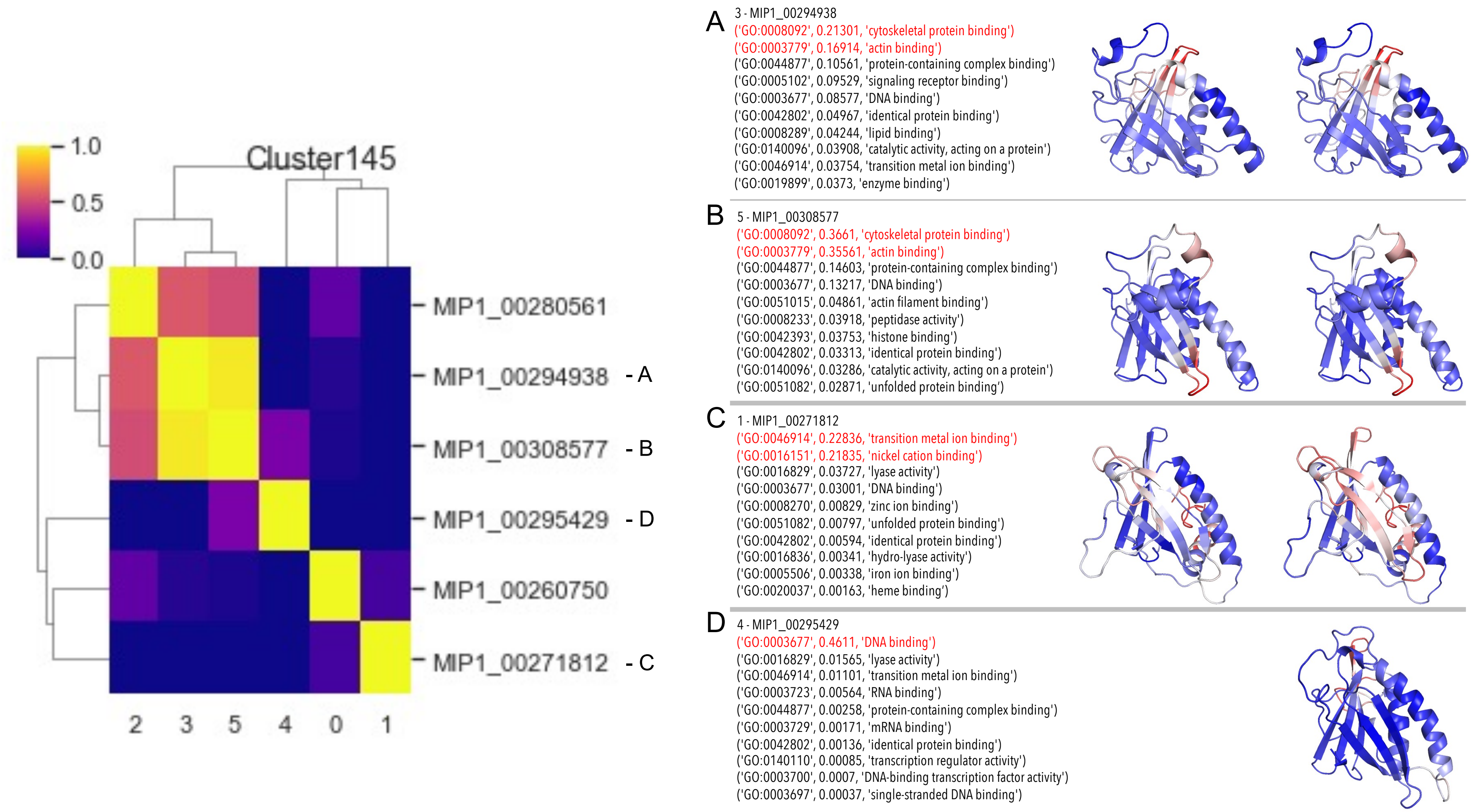

*Fig. S87. Cluster 145 with examples of different structural motifs producing similar functions. In A and B, cytoskeletal protein binding, specifically actin binding, are produced by a different $\beta$-turn in the structure.*

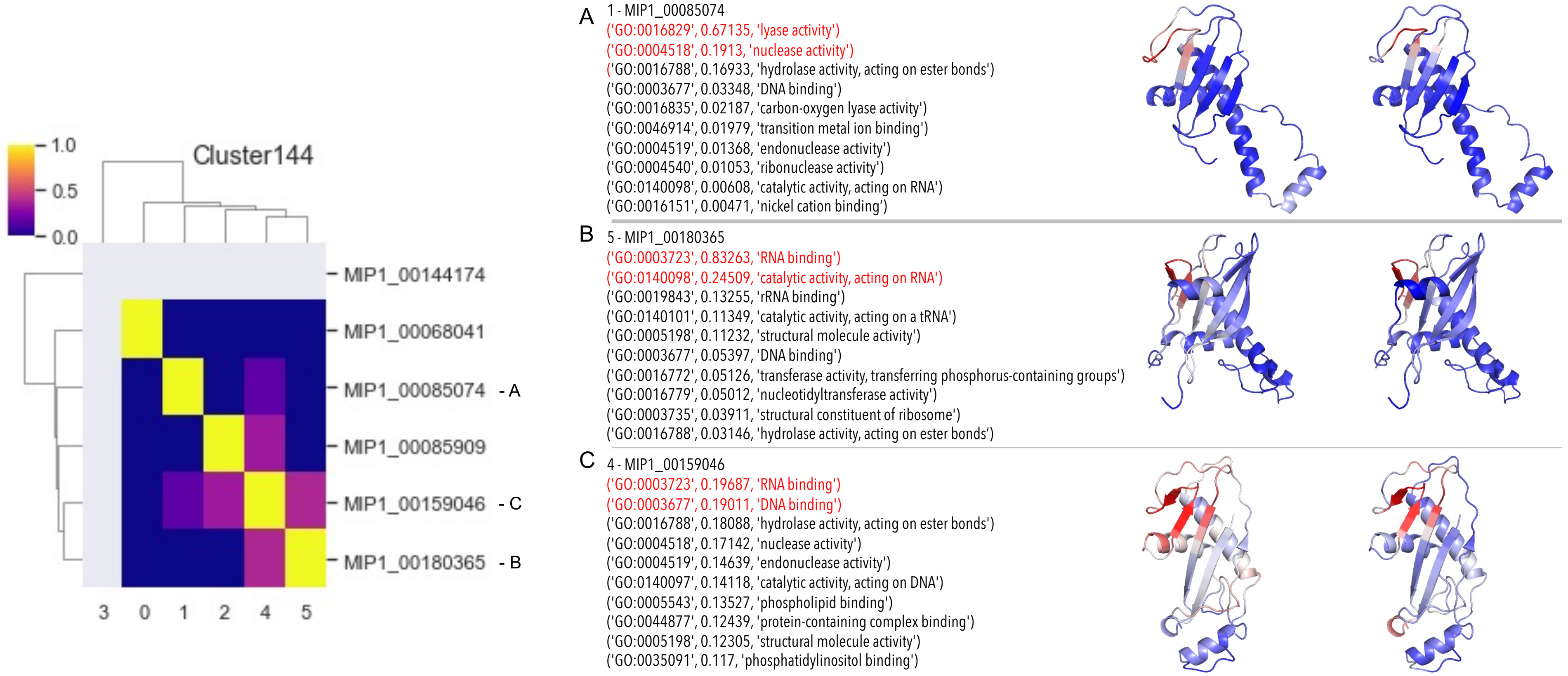

*Fig. S88. Cluster 144 with examples of similar structural motifs producing different functions. In A, the $\beta$-turn produces lyase and nuclease activity, whereas in B and C, the same turn is responsible for RNA and DNA binding.*

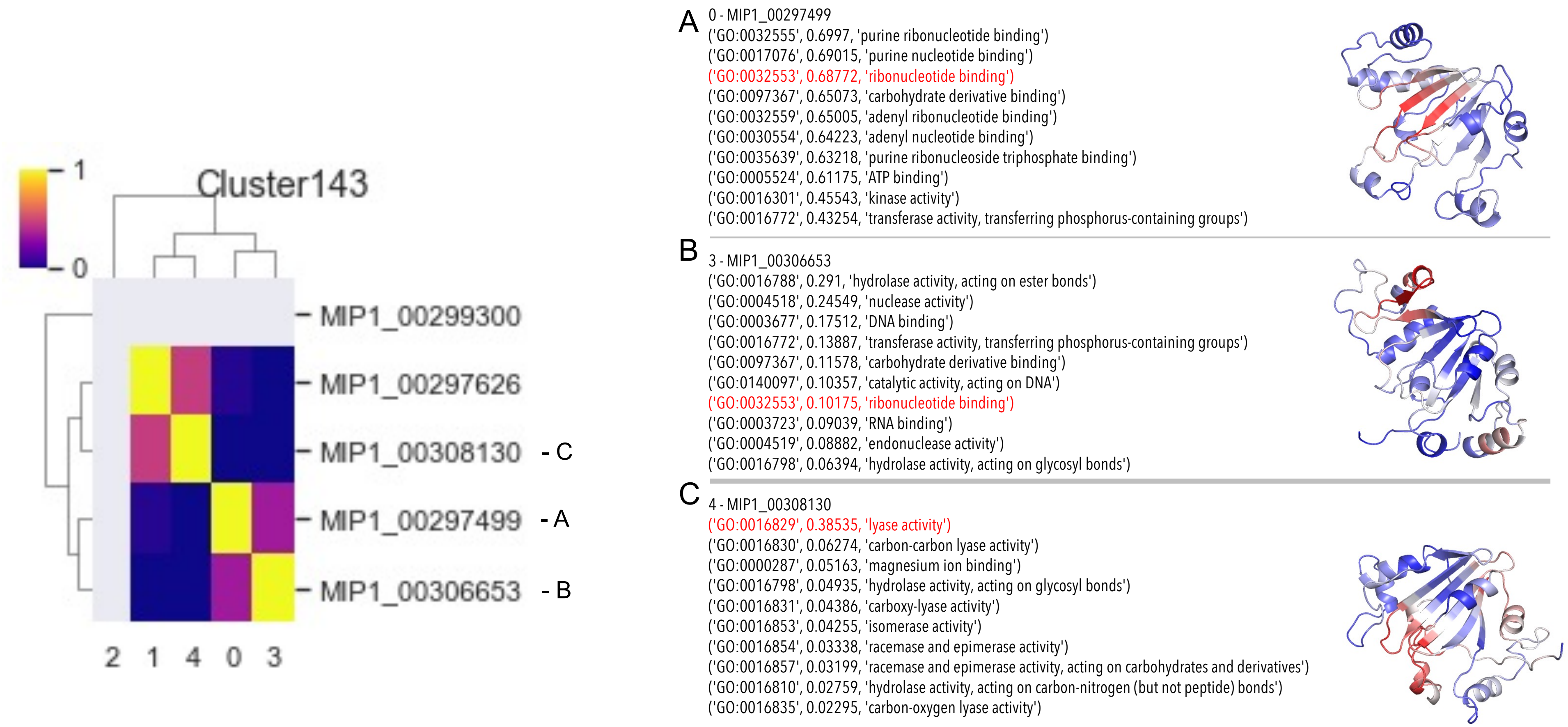

*Fig. S89. Cluster 143. (A) and (B) have overlap in the structural motif that produces the same function (ribonucleotide binding). (C) shows a different function (lyase activity) that is produced by a different structural motif.*

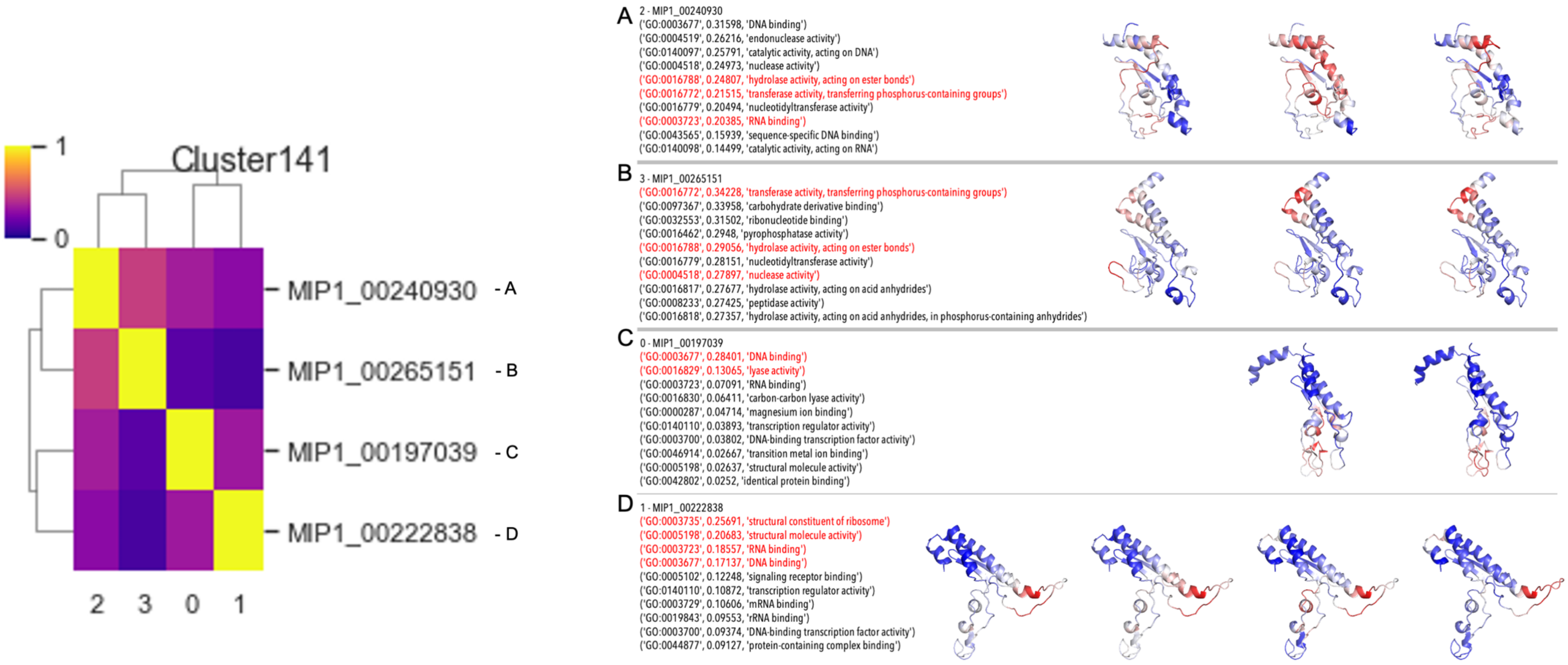

*Fig. S90. Cluster 141 includes examples where overlapping structural motifs having the same function. Transferase activity in (A) and (B) are carried out by some of the same residues in the structure and DNA binding in (C) and (D) are carried out by the same structural motif.*

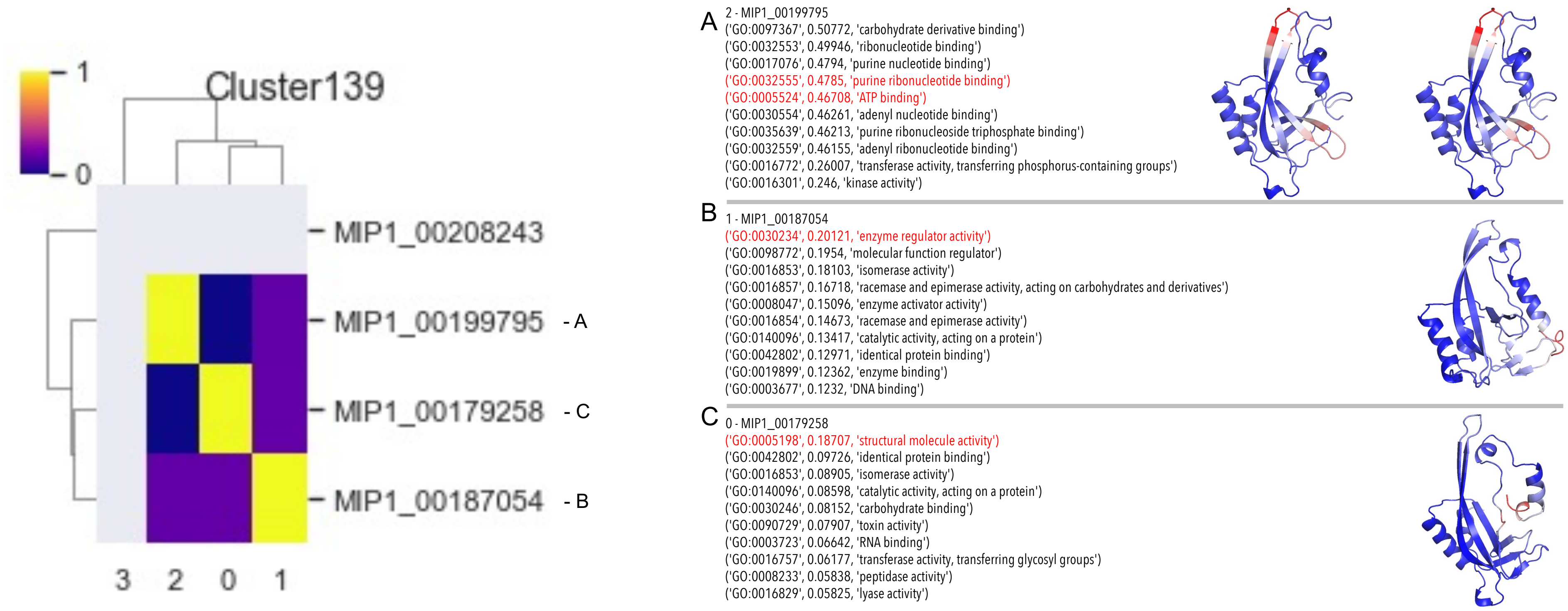

*Fig. S91. Cluster 139 contains examples have low functional similarity. The termini in (B) and (C) have either enzyme regular activity or structural molecule activity.*

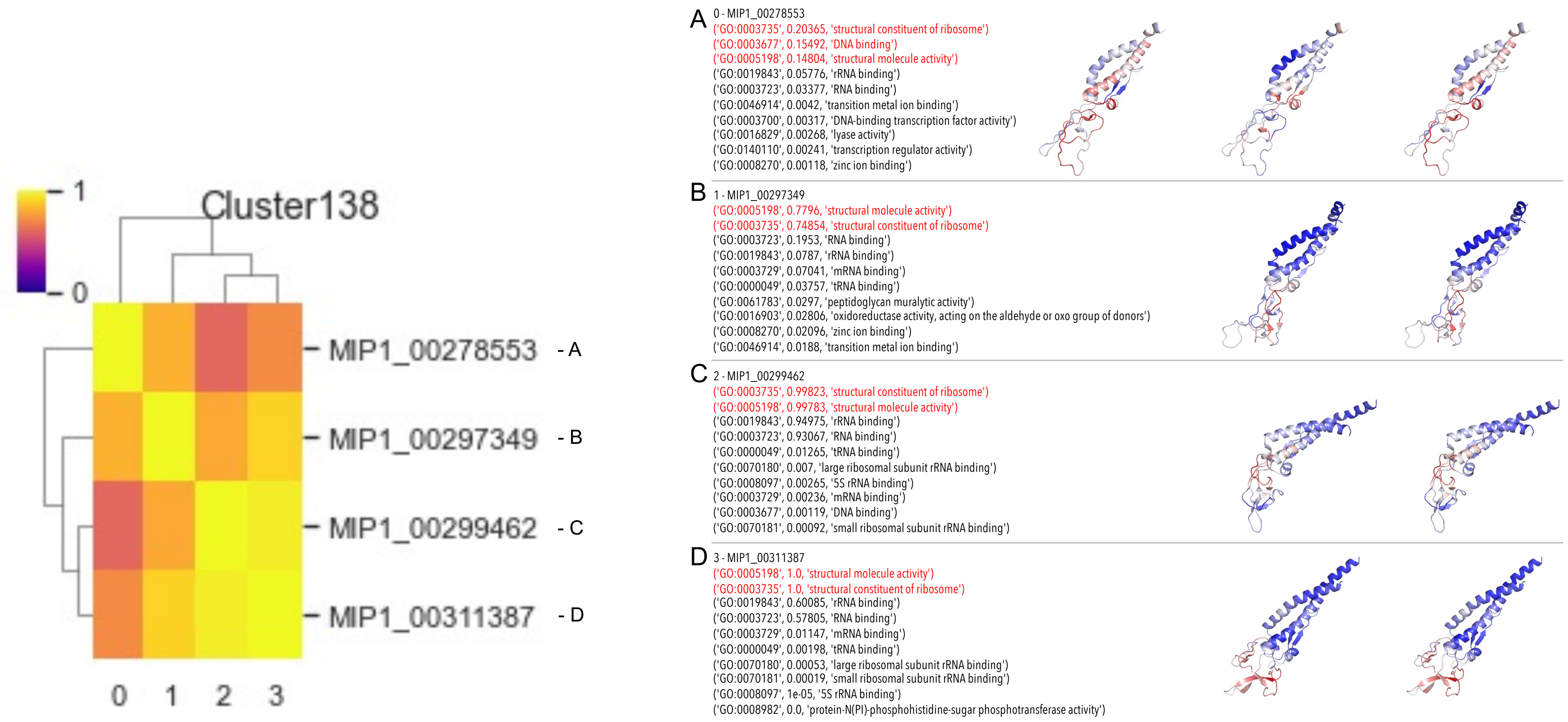

*Fig. S92. Cluster 139 with examples of similar structural motifs producing similar functions. While the two helices show functional activitity in some examples, the loop region consistently has structural molecule activity.*

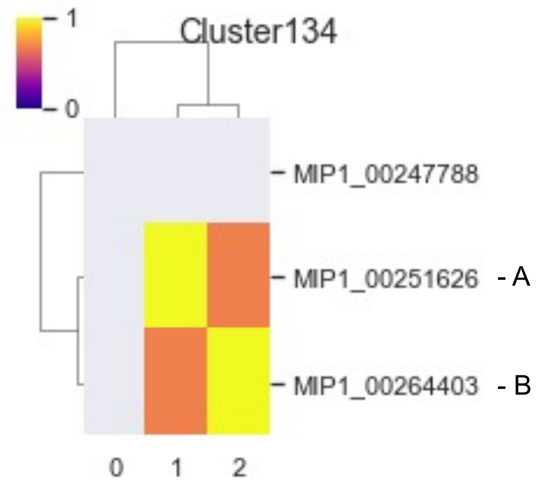

A

1 - MIP1_00251626

('GO:0016757', 0.40122, 'transferase activity, transferring glycosyl groups')
('GO:0016798', 0.26446, 'hydrolase activity, acting on glycosyl bonds')
('GO:0016758', 0.26312, 'transferase activity, transferring hexosyl groups')
('GO:0004553', 0.2209, 'hydrolase activity, hydrolyzing O-glycosyl compounds')
('GO:0016763', 0.15095, 'transferase activity, transferring pentosyl groups')
('GO:0008194', 0.12714, 'UDP-glycosyltransferase activity')
('GO:0046527', 0.06041, 'glucosyltransferase activity')
('GO:0030246', 0.04456, 'carbohydrate binding')
('GO:0016780', 0.02546, 'phosphotransferase activity, for other substituted phosphate groups')
('GO:0060089', 0.02343, 'molecular transducer activity')

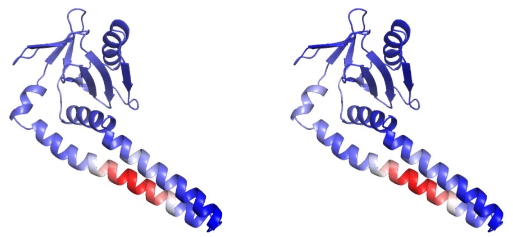

B

2 - MIP1_00264403

('GO:0016757', 0.53021, 'transferase activity, transferring glycosyl groups')
('GO:0016763', 0.25278, 'transferase activity, transferring pentosyl groups')
('GO:0016758', 0.22021, 'transferase activity, transferring hexosyl groups')
('GO:0022857', 0.1915, 'transmembrane transporter activity')
('GO:0005215', 0.17557, 'transporter activity')
('GO:0016772', 0.16742, 'transferase activity, transferring phosphorus-containing groups')
('GO:0008509', 0.16338, 'anion transmembrane transporter activity')
('GO:0015075', 0.15479, 'ion transmembrane transporter activity')
('GO:0140096', 0.12676, 'catalytic activity, acting on a protein')
('GO:0008194', 0.12526, 'UDP-glycosyltransferase activity')

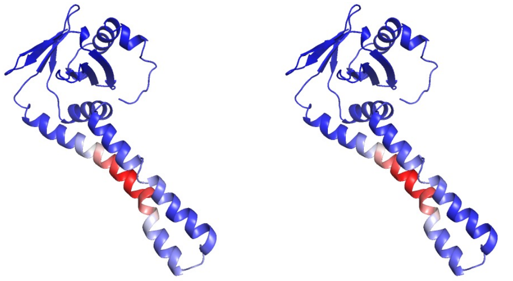

*Fig. S93. Cluster 134. While both proteins have an overlap of functions as shown in the heatmap, the function transferase activity, transferring glycosyl or hexosyl groups pertains to the same structural motif, i.e. the center of one of the helices.*

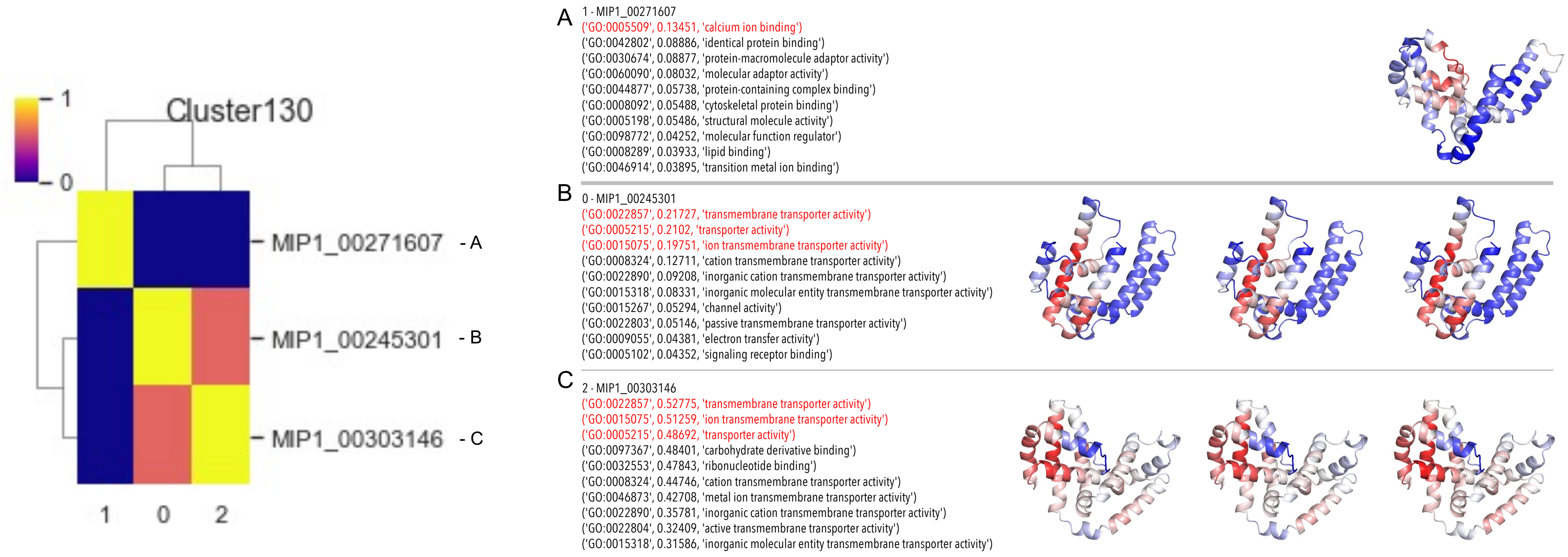

*Fig. S94. Cluster 130. The protein in (A) has mostly different functions than (B) and (C) and the latter two have overlapping structural motifs that generate the functions of transmembrane transporter activity.*

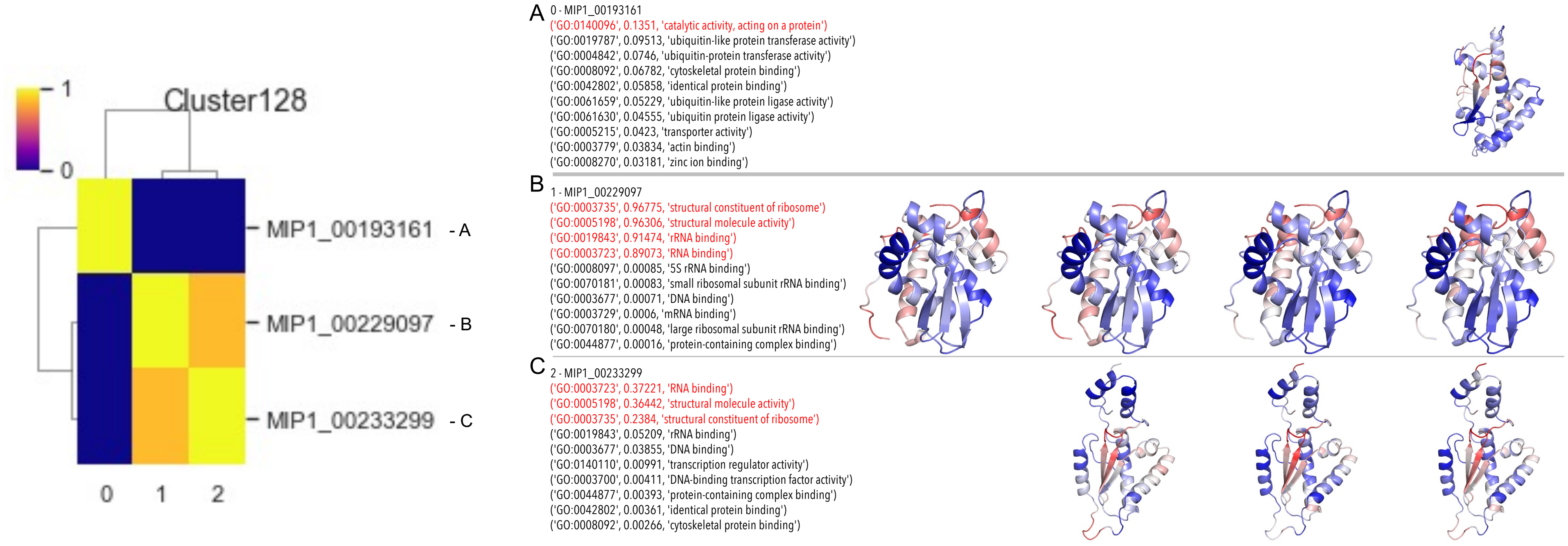

*Fig. S95. Cluster 128. The functions of (A) are very different than those of (B) and (C). For the latter two, the same functions are created by different structural motifs. Further, the superposition is poor because the N- and C-termini have different interfaces to the core of the fold.*

#### 13. Function-to-structure examples: comparing structures for specific functions

### BP GO:0030683 - mitigation of host immune response by virus

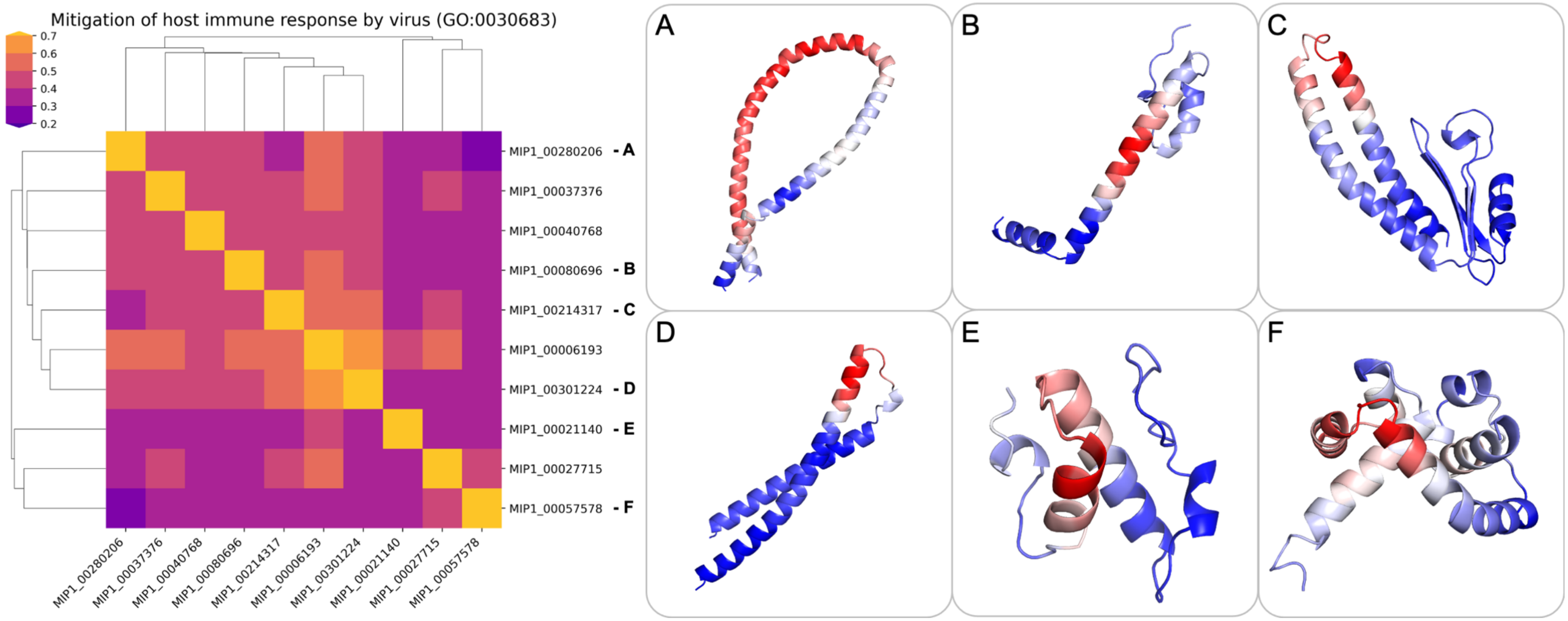

*Fig. S96: The figure shows a functional cluster with a heatmap of pairwise TM-scores of the structures in that cluster. The proteins in that cluster evade the host immune response by a virus; this GO-term has six child terms. The PDB has over 6,000 entries with the GO-term, many of which are virus capsid proteins or proteases. The structures in this cluster have very different folds but most often contain two helices. We suspect that different folds generate this function because the function is very general.*

### EC 2.7.10.- Protein tyrosine kinases

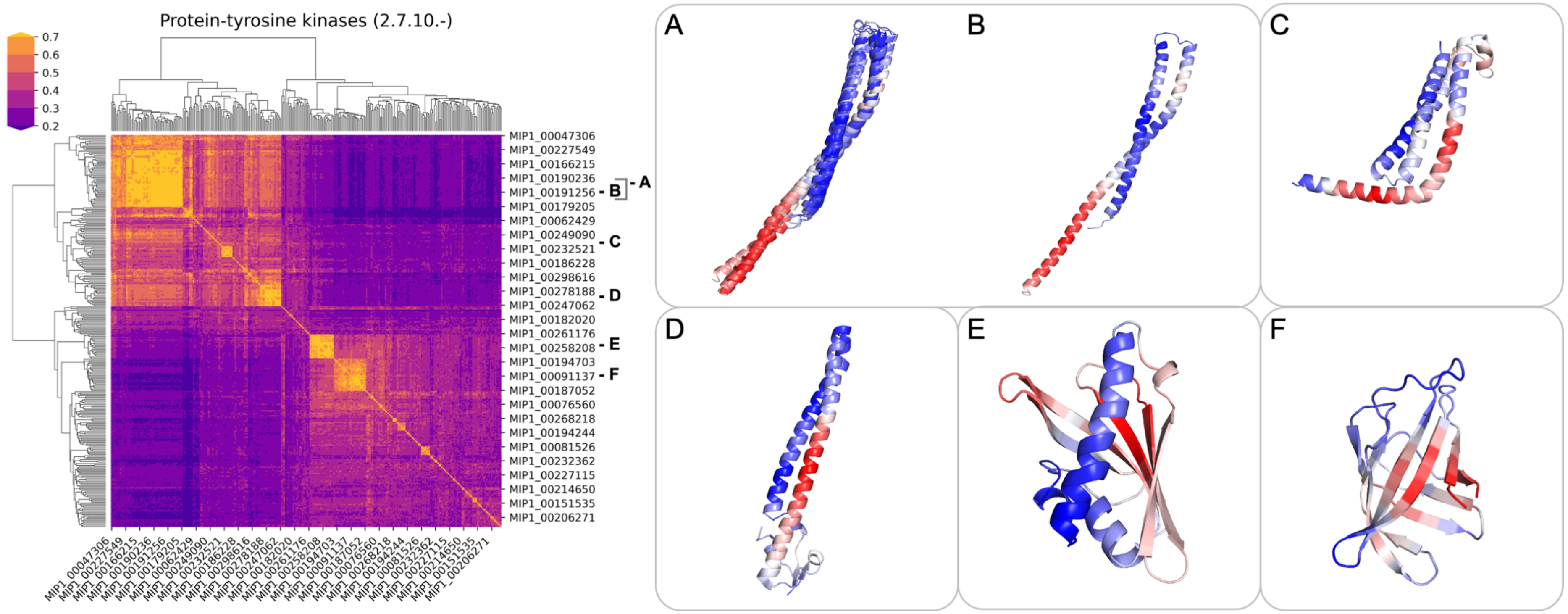

*Fig. S97: The figure shows a functional cluster with a heatmap of pairwise TM-scores of the structures in that cluster. The heatmap shows that the functional cluster is divided into two groups, predominantly helical, denoting the upper left yellow square and covering (A), (B), (C), and (D), and with predominantly $\beta$-sheet content, denoted by the cluster in the lower right and covering models (E) and (F). This structural diversity is corroborated by the generality of this particular function: Tyrosine kinases are a large and diverse group of enzymes that are involved in many key events in the body. They catalyze the reaction of transferring a phosphate group from ATP to tyrosine residues of specific proteins, essentially switching them "on" or "off". Phosphorylation of tyrosine residues in proteins controls functions such as subcellular localization, enzyme activity, and signal transduction. Tyrosine kinases come in two forms: transmembrane receptor tyrosine kinases (RTKs) and cytosolic non-receptor tyrosine kinases. RTKs are signalizing molecules that homo- and hetero-dimerize in the membrane and transduce the signal from outside of the cell to the intracellular region. RTKs also have a number of domains that dictate to which family they belong. Mutations in RTKs are implicated in a variety of cancers, making RTKs important drug targets.*

### EC 4.2.1.1 - carbonic anhydrase

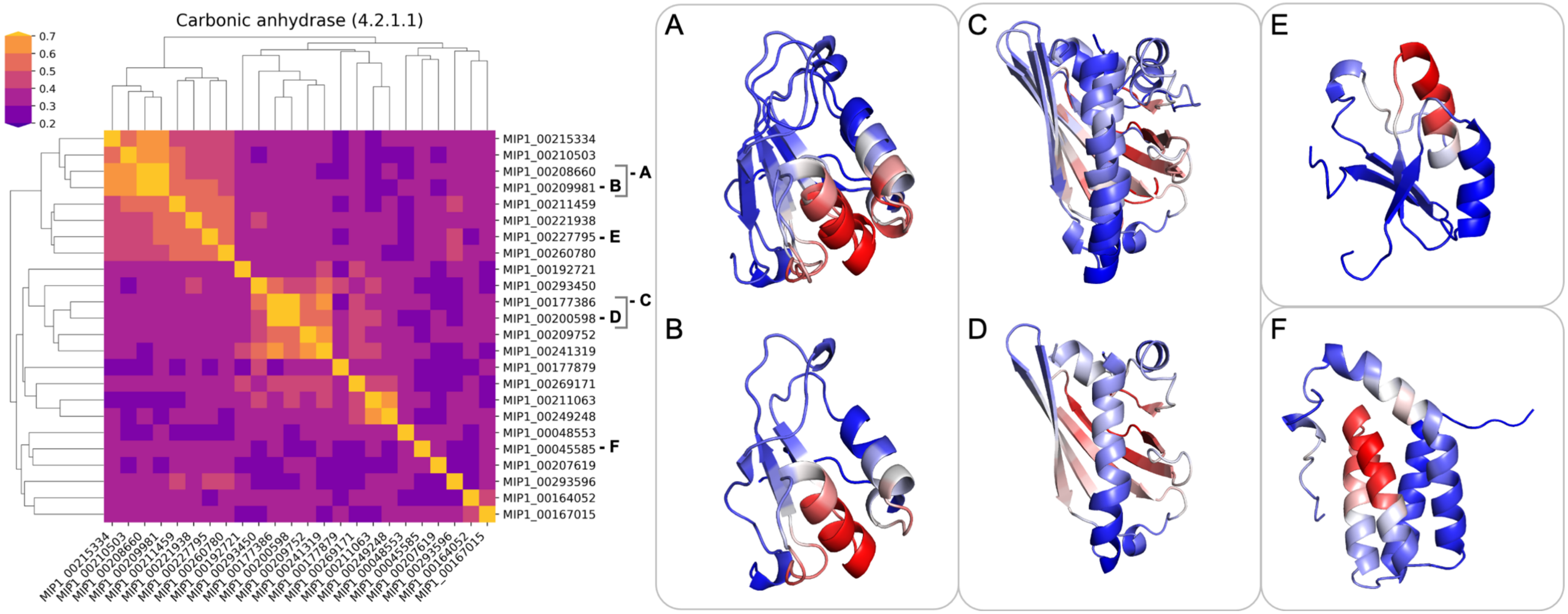

*Fig. S98: The figure shows a functional cluster with a heatmap of pairwise TM-scores of the structures in that cluster. In our MIP dataset, carbonic anhydrases are a smaller class of proteins with three small clusters, covering several different folds. Carbonic anhydrases are enzymes that catalyze the conversion of carbon dioxide to carbonic acid, often with the help of zinc in the active site, making them metalloenzymes. Carbonic anhydrases help regulate pH and fluid balance. Three main families ($\alpha$, $\beta$, $\gamma$) exist in addition to smaller families ($\delta$, $\zeta$, $\eta$, $\iota$) that are less well-studied: $\alpha$-CAs occur in mammals, $\beta$-CAs in bacteria and plants, and $\gamma$-CAs in methanogen bacteria in hot springs. The families are structurally different and appear to have evolved independently, which supports the structural diversity in this functional cluster in our MIP database. Interestingly, the fold in (C) superimposes perfectly with the structures of pathogenesis related proteins (for instance PDBID 4c94) for a variety of allergens and also cytokinin-specific binding proteins (for instance PDBID 2flh). The two latter proteins have an additional strand at the C-terminus that extends the twisted beta-sheet, which is not present in the MIP models.*

### EC 2.7.1.21 - thymidine kinase

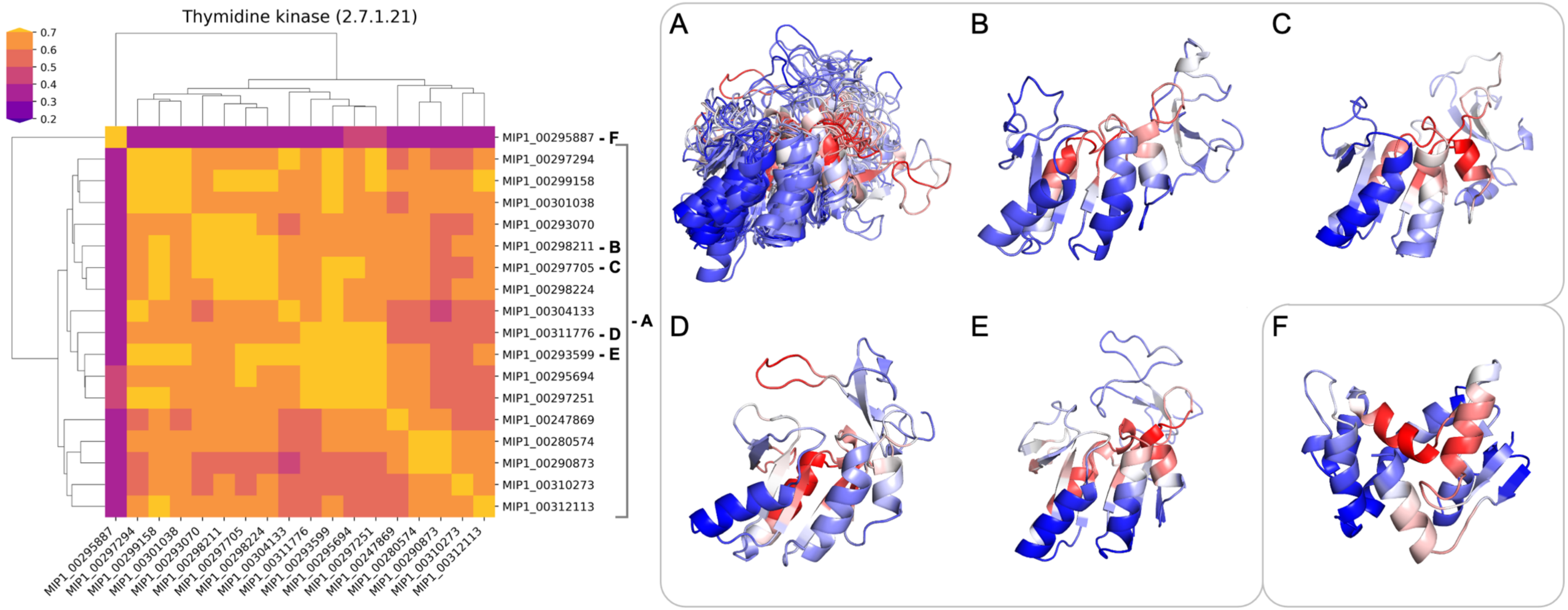

*Fig. S99: The figure shows a functional cluster with a heatmap of pairwise TM-scores of the structures in that cluster. The functional cluster shows one main fold (A) and one minor fold with a single representative (F). The salient residues in cluster (A) overlay nicely for all representatives and have similar sequence motifs for this cluster with sequences of GKST[SLIH]LL / [LFI][LIVC]DEAQFL / G[LI]RTD[FA]. Thymidine kinases are enzymes that catalyze the transfer of a phosphate group from ATP to thymidine, creating thymidine monophosphate and ADP. They are a key element in the synthesis of DNA, as they introduce thymidine into the DNA. There are two families, one found in herpesviruses, and one found in mammals, bacteria and viruses. The latter family encompasses two types, type I and type II, which differ structurally. The main cluster in the MIP dataset, shown in (A) superimposes almost perfectly with type II thymidine kinases, examples of which are PDB IDs 1w4r (human) and 2b8t (ureaplasma parvum). The structure is an alternating ($\beta\alpha$)x5($\beta$) fold containing six strands and five helices, in addition to a small sheet embedded in loop regions at the C-terminus. The structure in (F) does not overlay with any known thymidine kinase structures, so it is possible that this might be a divergent fold.*

### MF GO:0005125 - cytokine activity

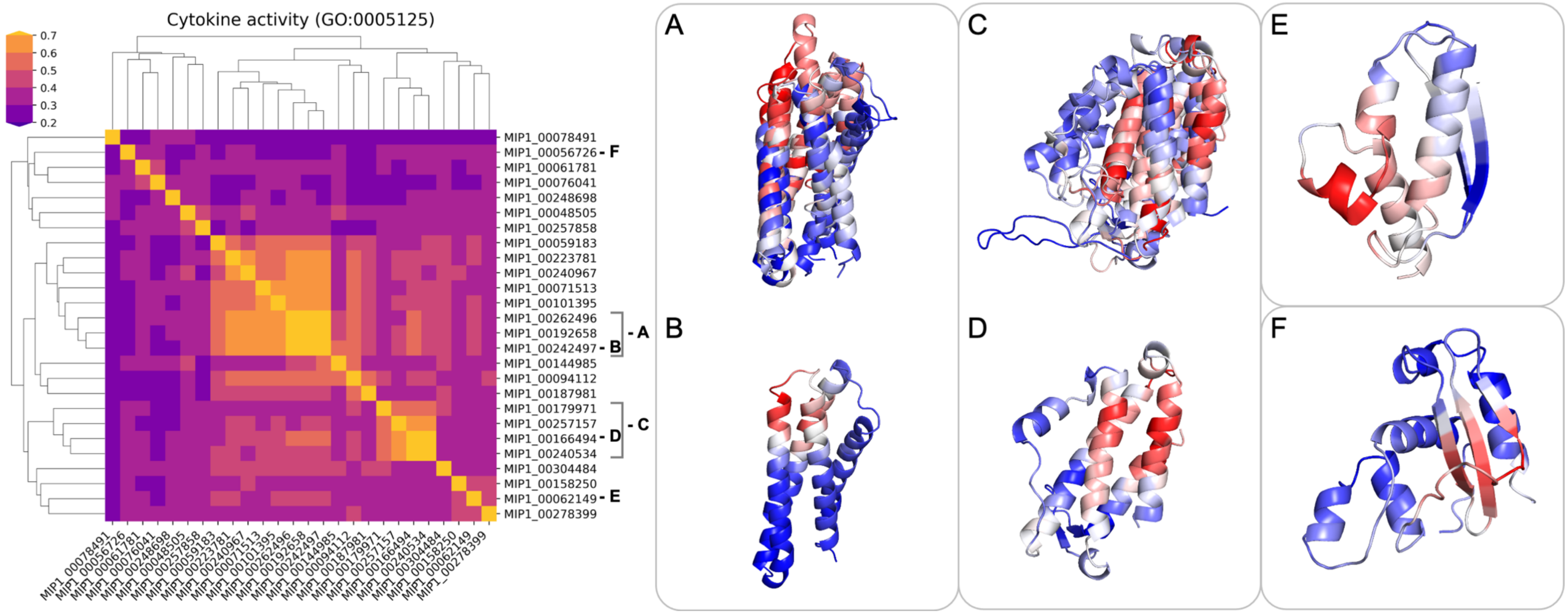

*Fig. S100: The figure shows a functional cluster with a heatmap of pairwise TM-scores of the structures in that cluster. Cytokines are small proteins involved in cell signaling and they cannot cross the membrane bilayer. Rather, they modulate the function of the receptors they interact with to control growth, survival, differentiation and effector functions of cells and tissues. Cytokines are immune-modulating agents and are therefore important for the immune system, yet they are not limited to it. Cytokines include interleukins, chemokines, interferons, lymphokines and tumor necrosis factors but generally do not include hormones or growth factors. Their terminology overlaps with hormones and the distinction between the two is still being researched. Hormones are important cell signaling molecules that act distantly from the production site, but usually circulate in much higher concentrations than cytokines (nanomolar vs. picomolar concentrations). In our MIP dataset, the functional cluster with predicted cytokine activity contains 26 members, covering two fold clusters and several other folds. The covered folds are often helical bundles or have high helical content. Given that cytokine activity is a broad function carried out by various proteins, their structures cover a variety of folds. Our MIP models look overall similar to known folds for interleukin or interferon, yet their helix/strand connections are different.*

### MF GO:0009881 - photoreceptor activity

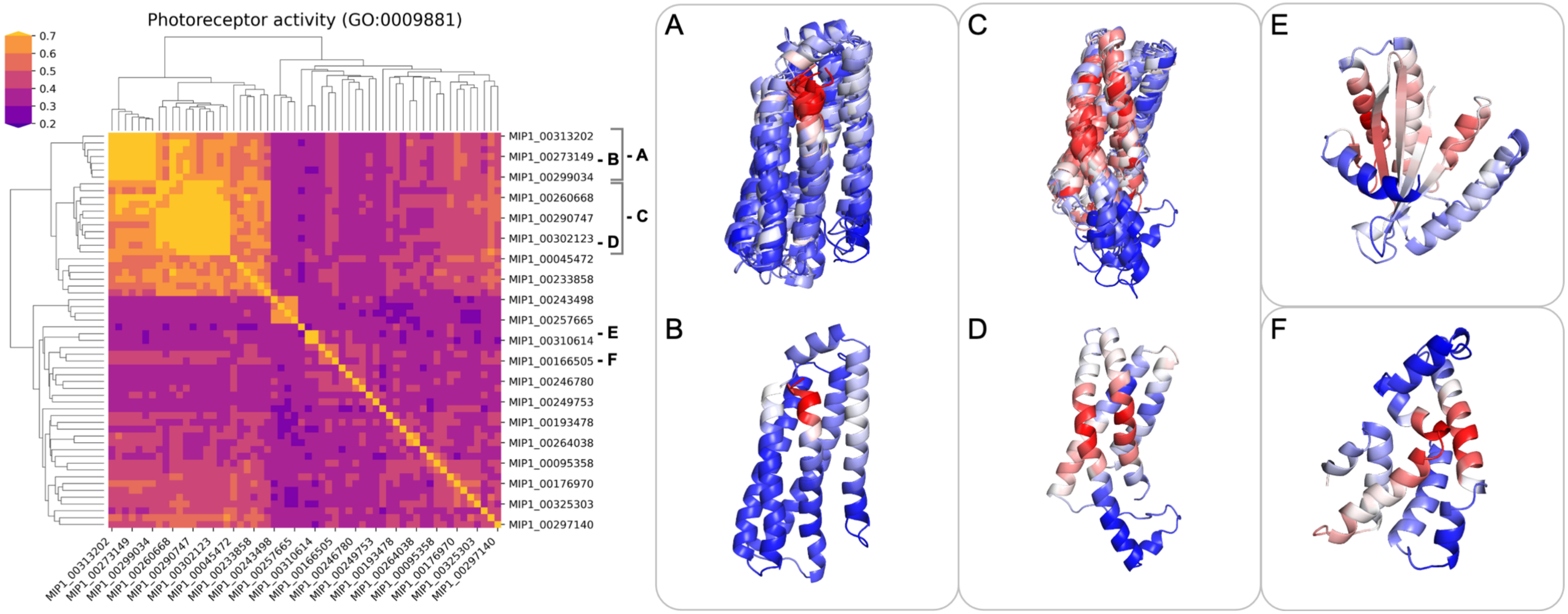

*Fig. S101: The figure shows a functional cluster with a heatmap of pairwise TM-scores of the structures in that cluster. In our MIP database, the functional cluster with predicted photoreceptor activity is an intermediate-sized cluster that has one large structural cluster (splitting into two smaller ones for 4 and 5-helix bundles), few smaller clusters and several other folds. The vast majority of structures in this functional cluster consists of helical bundles, covering different folds. Note that bacteriorhodopsin folds would not be covered here because they are with ~300 residues larger than cutoff of protein sizes we have predicted structures for. The clusters utilize similar salient residues to accomplish their function. One exception in terms of fold is the structure in (E) with a ($\alpha\alpha\alpha\beta\beta$)x2 fold. This structure looks similar to small $\alpha\beta$ photoreceptor proteins like LOV (light-oxygen-voltage-sensing) domains and photoactive proteins, but the helix/strand connections differ, making them a different fold. Photoreceptor activity is the response to light and examples of proteins with photoreceptor activity include rhodopsin in the retina of vertebrae, phytochromes in plants, bacteriorhodopsins and bacteriophytochromes in some bacteria. Photoreceptors usually have a photopigment ligand that reacts to light and induces a conformational change in the ligand, for instance through isomerization, which in turn triggers a conformational change in the receptor, causing a signaling cascade. Examples of pigments include retinal, flavin and bilin, even though some proteins work without pigments.*

### MF GO:0004930 - G protein-coupled receptor activity

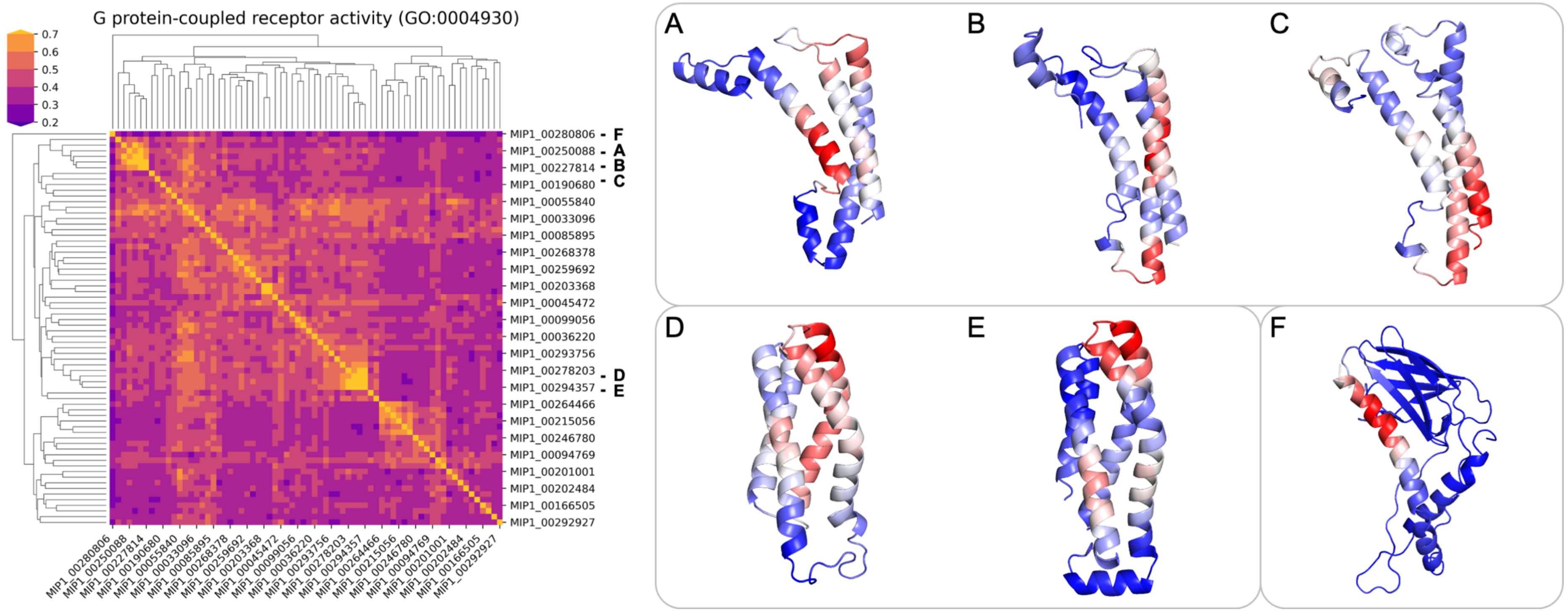

*Fig. S102: The figure shows a functional cluster with a heatmap of pairwise TM-scores of the structures in that cluster. The function GPCR activity covers an intermediate-sized cluster with several smaller clusters. The covered folds are vastly different, albeit mostly small, helical bundles and with very little strand content. The typical GPCR 7-helix bundle is not represented because these folds are with ~300 residues larger than the sequence-length cutoff of 200 residues we chose. QuickGO defines GPCR activity as "Combining with an extracellular signal and transmitting the signal across the membrane by activating an associated G-protein; promotes the exchange of GDP for GTP on the alpha subunit of a heterotrimeric G-protein complex."*
